# Species Tree Topology Impacts the Inference of Ancient Whole-Genome Duplications Across the Angiosperm Phylogeny

**DOI:** 10.1101/2024.01.04.574202

**Authors:** Michael T. W. McKibben, Geoffrey Finch, Michael S. Barker

## Abstract

**Premise:** The history of angiosperms is marked by repeated rounds of ancient whole-genome duplications (WGDs). Here we use state of the art methods to provide an up-to-date view of the distribution of WGDs in the history of angiosperms that considers both the uncertainty introduced by inference methods and alternative phylogenetic hypotheses.

**Methods:** Transcriptomic and genomic data were used to infer and place WGDs across two hypothesized angiosperm phylogenies. Initial WGD hypotheses were made using rate corrections to the distribution of synonymous divergences (K_s_) of paralogs and orthologs. WGD hypotheses were tested using syntenic inferences and Bayesian models of duplicate gene gain and loss across the phylogeny.

**Key results:** The number of ancient WGDs in the history of angiosperms (∼170) is largely similar across different inference methods, but there is often variation in the precise placement of WGDs on the phylogeny. K_s_ based methods often yield alternative hypothesized WGD placements largely due to variation in substitution rates among lineages. Phylogenetic models of duplicate gene gain and loss are more robust to topological variation, allowing for post hoc testing of WGD hypotheses. However, errors in species tree inference can still produce spurious WGD hypotheses regardless of method used.

**Conclusions:** Here we show that different WGD inference methods largely agree on an average of 3.5 WGD in the history of angiosperm species. However, the precise placement of WGDs on the phylogeny is subject to the inference method and tree topology. As researchers continue to test hypotheses regarding the impacts ancient WGDs have on angiosperm evolution, it is important to consider the uncertainty of the phylogeny as well as WGD inference methods.

## INTRODUCTION

Over 100 years after its discovery (Winkler, 1916; Winge, 1917), the number and distribution of ancient whole genome duplications (WGDs) or paleopolyploidization events across the angiosperm phylogeny remains uncertain. Regardless, the high frequency of ancient WGDs in angiosperms has spurred questions such as the association of WGD with the emergence of traits (Jiao et al., 2011; Soltis and Soltis, 2016; Moriyama and Koshiba-Takeuchi, 2018), timing of major environmental perturbations (Vanneste et al., 2014; Van de Peer et al., 2017; Wu et al., 2020), and diversification rate shifts (Vanneste et al., 2014; Tank et al., 2015; Landis et al., 2018). At the same time, increasing availability of genomic data has encouraged researchers to develop and improve methods for inferring and placing WGD on phylogenies. Despite these innovations, few studies have attempted to apply current tools across a broad phylogenetic context (One Thousand Plant Transcriptomes Initiative, 2019). Similarly, relatively few studies have explored the influence species tree and WGD inference have on each other (Shi, 2013; Struck, 2013; Smith and Hahn, 2022; Xiong et al., 2022), and how this shapes our estimation of the WGD history of flowering plants. When forming hypotheses for the role of WGD in plant evolution it is important to consider that the species tree itself is a hypothesis, to which any hypothesized WGD are nested within. This chain of inference can result in varying degrees of uncertainty that should be carefully considered when testing models of evolution. For all these reasons, there is a need for an updated description of the distribution of WGD across flowering plants that considers the sensitivity of inferences to variation in tree topology.

Methods for detecting and placing WGDs on phylogenies generally utilize three data types. The simplest of these is to estimate the synonymous divergence (K_s_) of duplicated genes in genomic or transcriptomic data (Barker et al., 2008; Ambrose and Purugganan, 2012; Vanneste et al., 2014; Tiley et al., 2018). When plotted as a histogram, WGDs show up in K_s_ distributions as peaks, and mixture models can be used to ascertain a median divergence among the paralogs (Tiley et al., 2018). The median K_s_ can then be compared to the ortholog divergences with other species to determine if the WGD is in the common ancestry of the species of interest. Inferences using K_s_ plots can also use rate corrections that account for variable rates of synonymous substitutions among lineages (Barker et al., 2008, 2009; Shi et al., 2010; Sensalari et al., 2022). If genomic data are available, K_s_ based methods are often accompanied with genome structural information. By making inter- or intra-genomic comparisons, regions where genes occur in the same order—collinearity—can be detected (Coghlan et al., 2005; Tang et al., 2008a). In the case of WGD, entire chromosomes are duplicated and often rearranged, forming groups of collinear genes commonly referred to as syntenic blocks (Renwick, 1971; Passarge et al., 1999; Kellis et al., 2004; Byrne and Wolfe, 2005; Jiao et al., 2014). Other processes can create syntenic blocks and plant species often experienced more than one WGD, sometimes making it difficult to ascertain if two species share the same WGD history (Tang et al., 2008a, 2011; One Thousand Plant Transcriptomes Initiative, 2019; Siddiqui and Conant, 2023). Despite these limitations, the availability of data and the accessibility of tools that can quickly generate K_s_ and syntenic inferences have made both approaches popular (Haas et al., 2004; Barker et al., 2010; Tang et al., 2011; Wang et al., 2012; Zwaenepoel and Van de Peer, 2019b; Henry et al., 2022; Sun et al., 2022).

In addition to methods focused on WGD inference in one or two species at a time, a variety of phylogenetic approaches have been developed to infer and place WGDs in the phylogenetic history of multiple species. When WGD occurs, the large influx of duplicate genes is expected to cause increased duplication rates along specific branches in the species tree. Several methods have been developed to estimate gene duplications along a phylogeny through gene counting (Rabier et al., 2014; Zwaenepoel and Van de Peer, 2020), using species trees to count duplicate subtrees in gene trees (Li et al., 2015, 2018), mapping clade specific duplications in gene trees onto a species tree (Yang et al., 2015), and gene tree-species tree reconciliations (Chen et al., 2000; Pfeil et al., 2005; McKain et al., 2016; Thomas et al., 2017; Zwaenepoel and Van de Peer, 2019a). The most recent iterations of these methods account for variation in the background rate of small scale duplications (SSD) across a phylogeny when testing for the presence of WGD (Zwaenepoel and Van de Peer, 2019a, 2020). Among these methods, WHALE offers a comprehensive framework to model and test hypothesized placements of WGD on a phylogeny. It does so by considering gene tree-species tree reconciliation in a Bayesian context, allowing for the uncertainty of gene tree topologies to be explicitly modeled (Zwaenepoel and Van de Peer, 2019a). One limitation of WHALE is it does not detect WGD itself, but rather allows users to test prior WGD hypotheses inferred using other methods. Due to this limitation, the hypothesis generation step is critical and should carefully be considered by researchers.

Here we applied state of the art methods on a sample of over 400 angiosperm species using two different species tree hypotheses. Initial WGD hypothesis generation was conducted using a K_s_ based approach with KsRates (Sensalari et al., 2022). Using this method, we were able to produce multiple alternative hypotheses for the placement of WGDs in the history of flowering plants. We further assessed the support for these placements using two models available in WHALE (Zwaenepoel and Van de Peer, 2019a). In doing so, we explicitly account for variability in the gene duplication and loss rates across our sampled species. To provide additional support for WGD inferences and placements, we compared patterns of synteny among more than 100 reference genomes. We conducted these analyses using two prior published species tree topologies (Kumar et al., 2017, 2022; Janssens et al., 2020) in order to better understand the sensitivity of current methods to topological variation and also give a broad overview of the WGD history across angiosperms that reflects the uncertainty of current methods and data. Given the increasing availability of genomic data, further exploration into how tree topologies and methods influence WGD inferences is critical as we form and test hypotheses of how WGDs have impacted plant evolution.

## MATERIALS AND METHODS

### Sequence Datasets

We collected 462 genomes and transcriptomes for the flowering plants and outgroups to infer and place WGDs across the angiosperm phylogeny. We downloaded 108 publicly available flowering plant genomes from Ensemble, Phytozome, NCBI, and several smaller databases ( (Appendix S1,Table S1; see Supplemental Data with this article). To fill in unsampled sections of the angiosperm phylogeny we included transcriptomes from the One Thousand Plant Transcriptomes Initiative (1KP)(2019). Missing gene annotations and inflated sample sizes can make the use of the Bayesian models of gene duplication inaccurate or computationally infeasible (Zwaenepoel and Van de Peer, 2019a; Chen et al., 2023). To accommodate these limitations, we filtered the 1KP dataset by only considering species that shared previously hypothesized WGD. For each of these previously inferred WGDs, we retained the two descendant species with the highest BUSCO score along with the next closest outgroup that did not have the WGD in its ancestry. After applying this procedure to the 1KP data set, we went through the dataset a second time, and for any family that was not represented in the post-filtering dataset we added back one representative species with the highest BUSCO score. Following this final addition of family representatives, we retained 1KP transcriptomes for 354 species in addition to the 108 reference genomes for subsequent analyses (Table S1-3).

### Phylogenetic Datasets

We used two different methods to obtain time calibrated phylogenies for WGD inference with WHALE. We obtained one set of time calibrated phylogenies from TimeTree.org (November 2022), a service that provides tree topologies and species divergence times conglomerated from the literature (Kumar et al., 2017, 2022). To explore how tree topology influences the placement of WGDs in the phylogeny we used a second time calibrated phylogeny (Janssens et al., 2020). In contrast to the novel time calibration approach used by TimeTree, the Janssens et al. (2020) tree is based on a more traditional dating approach using a phylogeny inferred from *matK* and *rbcl* sequences across 36,101 species and uses BEAST.v1.10 to develop a calibrated tree based on 56 fossil calibrations. Several species in our dataset were not present in one or both of the selected trees. For these species we assigned the leaf of the next closest species in the phylogeny as a representative sample (Table S2,3). To make computation time and analyses with WHALE feasible we split each of these phylogenies into 42 analyses of approximately 10–20 species each (Table S2,3). The placement of breaks between analyses were based on prior inferences in 1KP (2019) so that hypothesized duplications would be on internal branches of each analyses. For locations in the phylogeny where the duplications were near analysis breaks we included an overlap of two or more species between neighboring analyses.

### Hypotheses for WGD Placement

WHALE requires prior hypotheses for the placement of WGD on the phylogeny. To initially infer WGDs across our phylogenomic data sets, we used the KsRates package to generate Ks plots for each species and conduct rate corrections among species (Sensalari et al., 2022). For KsRates analyses we used the default settings and maximum number of outgroups equal to the number of species present in the focal analysis. A WGD was considered to be shared among species if the median K_s_ of the paralog peak was greater than the median K_s_ of the divergence among each species. As rate corrections were made for each species in each analysis, multiple hypothesized placements of each WGD could be inferred. Following inferences, we removed any hypothesized WGD that occurred prior to divergence with the outgroup or had a median Ks =< 0.1 (Tiley et al., 2018)(Table S2,3, Supplemental Data).

### WHALE Analyses

We used WHALE (Zwaenepoel and Van de Peer, 2019a) to test the inferences and phylogenetic placements of WGDs hypothesized by our KsRates analyses. WHALE requires a posterior distribution of gene tree topologies for each gene family in the phylogenetic sample (Zwaenepoel and Van de Peer, 2019a). Gene families were first inferred using OrthoFinder.v2.5.5 with default settings (Emms and Kelly, 2019), and filtered to remove extremely large and small gene families using orthofilter.py available from WHALE (Zwaenepoel and Van de Peer, 2019a). Gene families that passed the filtering step were aligned using Prank.v.170427 and analyzed with MrBayes.3.2.7a to generate a posterior distribution of gene trees under an LG + GAMMA model of sequence evolution (Ronquist et al., 2012; Emms and Kelly, 2019; Löytynoja, 2021). Each posterior distribution of trees was generated from 11,000 trees sampled every 10 generations from a model ran for 110,000 generations. From these distributions of trees we produced an Amalgamated Likelihood Estimation (ALE) object using ALEobserve, discarding the first 1,000 trees as burnin (Szöllõsi et al., 2013). Finally, we used the ccddata.py and ccdfilter.py scripts available from WHALE to filter out gene families with a high degree of uncertainty in their gene-tree topologies (Zwaenepoel and Van de Peer, 2019a)(Supplemental Data).

The rate of small-scale gene duplication and loss can vary across the phylogeny and impact the inference of whole-genome duplication events (Zwaenepoel and Van de Peer, 2019a; Chen et al., 2023). To accommodate this variation in duplication and loss rates, we ran two analyses on each analysis across the TimeTree (Kumar et al., 2017, 2022) and Janssens et al. (2020) topologies. The first of these—the “critical” model—allows the rate of gene duplication and loss to vary among branches, but the duplication and loss rates are constrained to be equal within branches. A second “relaxed” model allows the duplication and loss rates to be unconstrained and vary independently among and within branches, representing more biological realistic processes in exchange for increased parameterization. Following Chen et al. (2023), for both of these models we utilized a *Beta*(3,1) prior distribution for the number genes at the root of each tree (η). In the critical model, the mean duplication-loss branch rate (r) was set to an improper flat prior and each branch rate (λ,μ) was assumed to follow a multivariate Gaussian prior with a standard deviation (σ) from an exponential prior with mean equal to 0.1. In the relaxed model we allowed log-duplication-loss rates (r) to vary independently following bivariate normal prior with mean of 0 and standard deviation of 1. Within branches we assumed a *Uniform*(-1,1) prior for the correlation coefficient (ρ) between duplication and loss rates, and an exponential prior with mean of 1 for the standard deviation of branch rates (τ). For both models we sampled the posterior 500 times, allowing the ESS to reach a value of 100 or greater (average 513)(Supplemental Data). Finally, for each hypothesized WGD placement we calculated the log10 Bayes factor of the marginal posterior distribution of the retention rate parameter (q) under the critical and relaxed models (Supplemental Data).

### Syntenic Analyses

In addition to WHALE, we used syntenic inferences to further test WGD hypotheses proposed by KsRates. To detect synteny between the genomes in our dataset we used the python distribution of MCScan in a pairwise fashion (Tang et al., 2008b). For each WGD hypothesis we compared the syntenic depth ratio between ingroup species and outgroup species up to two nodes away from the WGD (Supplemental Data). For each ingroup species, the presence of the hypothesized WGD in its ancestry was considered supported if 50% or more of the syntenic depth comparisons with the outgroup species had a synthetic depth ratio greater than 1:1. The placement of the WGD on the phylogeny was then considered supported if all species daughter to it in the phylogeny met the criterion for the syntenic depth ratio test described. Finally, we used ete3.v3.1.3, matplitlib.v3.8, and seaborn.v0.13.0 to visualize syntenic bi-direction comparisons as a heat map arranged following Janssens et al. (2020) tree topology (Hunter, 2007; Huerta-Cepas et al., 2011; Waskom, 2021).

## RESULTS

### Ks Based WGD Hypotheses

Using KsRates, we inferred 246 and 244 hypothesized WGDs on the TimeTree (Kumar et al., 2017, 2022) and Janssens et al. (2020) topologies respectively (Table S2,3). Many of these placements represent the same duplication event shifted up or down one or more nodes depending on which species was used for the rate correction and how KsRates fit mixture models to the paralog distributions. For example, in analysis A1 of the TimeTree topology, the previously described ARTHβ duplication was variously placed to both include and exclude *Limnanthes douglasii* (Fig. 1). These alternative placements were due to some rate corrections, such as with *Batis maritima,* placing divergence with *L. douglassi* (2.05) prior to the paralog median K_s_ (1.98), whereas others placed it after (1.49 using *Salvadora persica*)(Supplemental Data). Similarly, we observed that the mixture models fit by KsRates occasionally placed the distributions of overlapping WGD closer to each other than prior work. For example in *Brassica rapa*, KsRates inferred a median K_s_ of 0.31 and 0.35 for the BRNIɑ and ARTHɑ WGD (Table S2,3). Previous analyses placed these duplications at median K_s_ of ∼0.3-0.4 and ∼0.6-0.7 respectively (Barker et al., 2009; Wang et al., 2013; Kagale et al., 2014; Jeong et al., 2016). All of these model differences were further exacerbated by topological disagreements between the two time calibrated phylogenies. Continuing the example from analysis A1, *Carica papaya* and *Moringa oleifera* were placed at a shallower node in the TimeTree topology compared to the Janssens et al. (2020) topology, resulting in disagreement whether they shared the ARTHβ duplication or not (Fig. 1). This shallower placement is in disagreement with the current understanding of the evolutionary history of the Brasicales (Barker et al., 2009; Edger et al., 2018; Mabry et al., 2020; Walden et al., 2020), whereas the Janssens et al. (2020) topology is more consistent with previous work in the group as are the WGD inferences.

**Figure 1:**
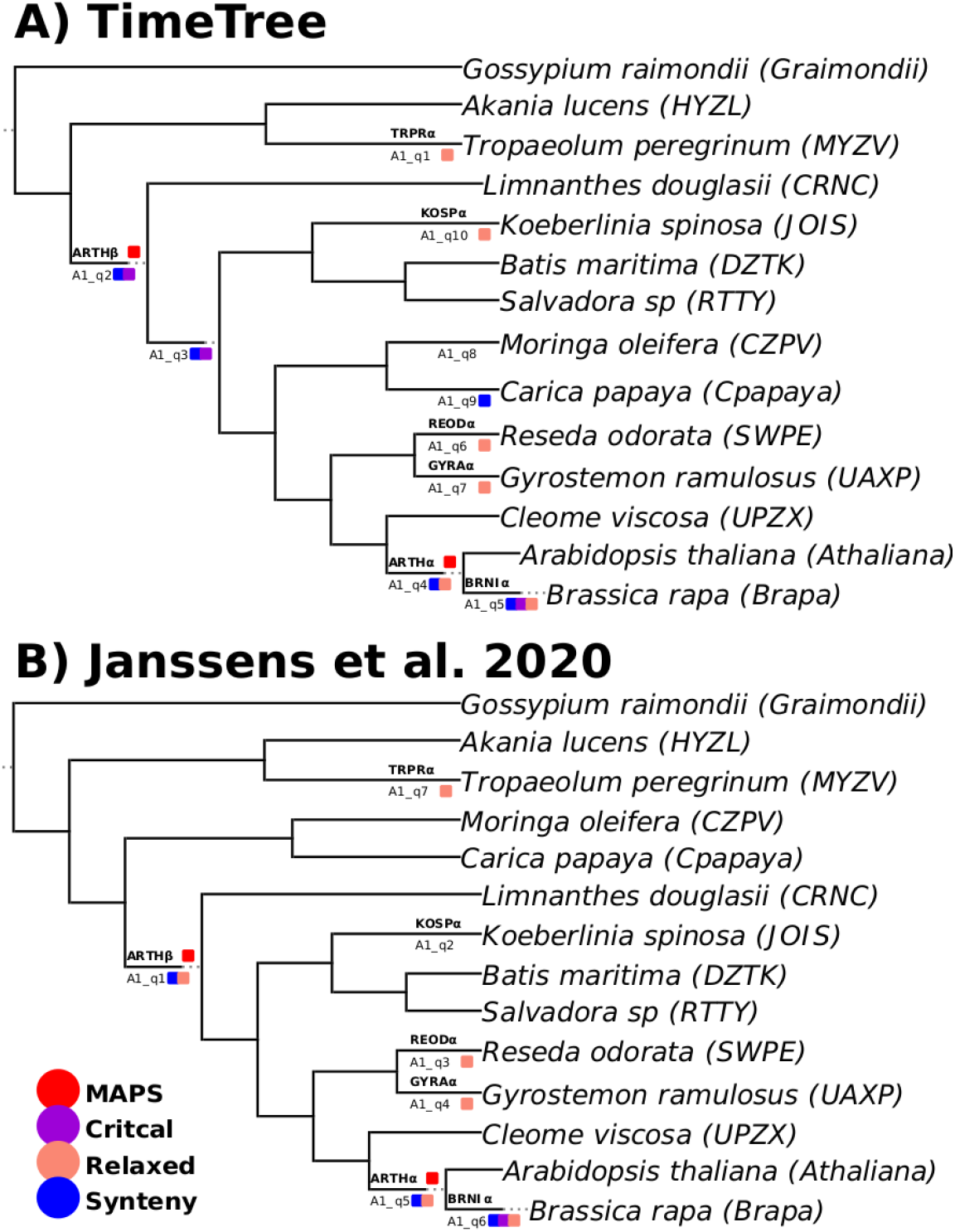
Comparison of support from KsRates (“analysis”_”WGD node”), syntenic (blue), WHALE Critical (purple), relaxed model (peach), and MAPS (red) for the placements of WGD within the Brassicales under the A) TimeTree and B) Janssens et al. (2020) tree topologies. The duplication naming convention from 1KP 2019 was used.

### WGD Hypothesis Testing with WHALE

We used WHALE to assess alternative hypotheses for the placement of WGD in the angiosperm phylogeny using paired analyses in the TimeTree (Kumar et al., 2017, 2022) and Janssens et al. (2020) phylogenies (Zwaenepoel and Van de Peer, 2019a). We conducted 42 analyses of an average species size of 12 for both the TimeTree (Kumar et al., 2017, 2022) and Janssens et al. (2020) species trees. For each analysis we used an average of 9,754 orthofamilies (4,804-12,180) as input data for WHALE (Supplementary Data). We used two models to assess our hypotheses, one that constrained rates of duplication and loss to be equal within branches but vary among branches, the critical model, and another that allowed duplication and loss rates to vary among and within branches, the relaxed model (Zwaenepoel and Van de Peer, 2019a; Chen et al., 2023). Across all of our analyses the median parameter ESS was 432 (513 average), suggesting a good approximation of the posterior using both models(Supplementary Data). In the TimeTree topology the critical WHALE model supported the inference and placement of 56 out of the 246 (∼23%) hypothesized placements made by KsRates (Fig. 2, Table S4). A much larger fraction of the hypothesized WGD inferences were supported by the relaxed model (167, ∼65%)(Fig. 2, Table S4). A similar pattern was observed in the Janssens et al. (2020) tree topology, where 47 (∼19%) and 154 (∼63%) of the 244 hypotheses were supported by the critical and relaxed models respectively (Fig. 2, Table S5). In total, 173 and 159 WGD hypotheses in the TimeTree and Janssens et al. (2020) analyses were supported by at least one of the WHALE models (Fig. 2, Table S4,5).

**Figure 2:**
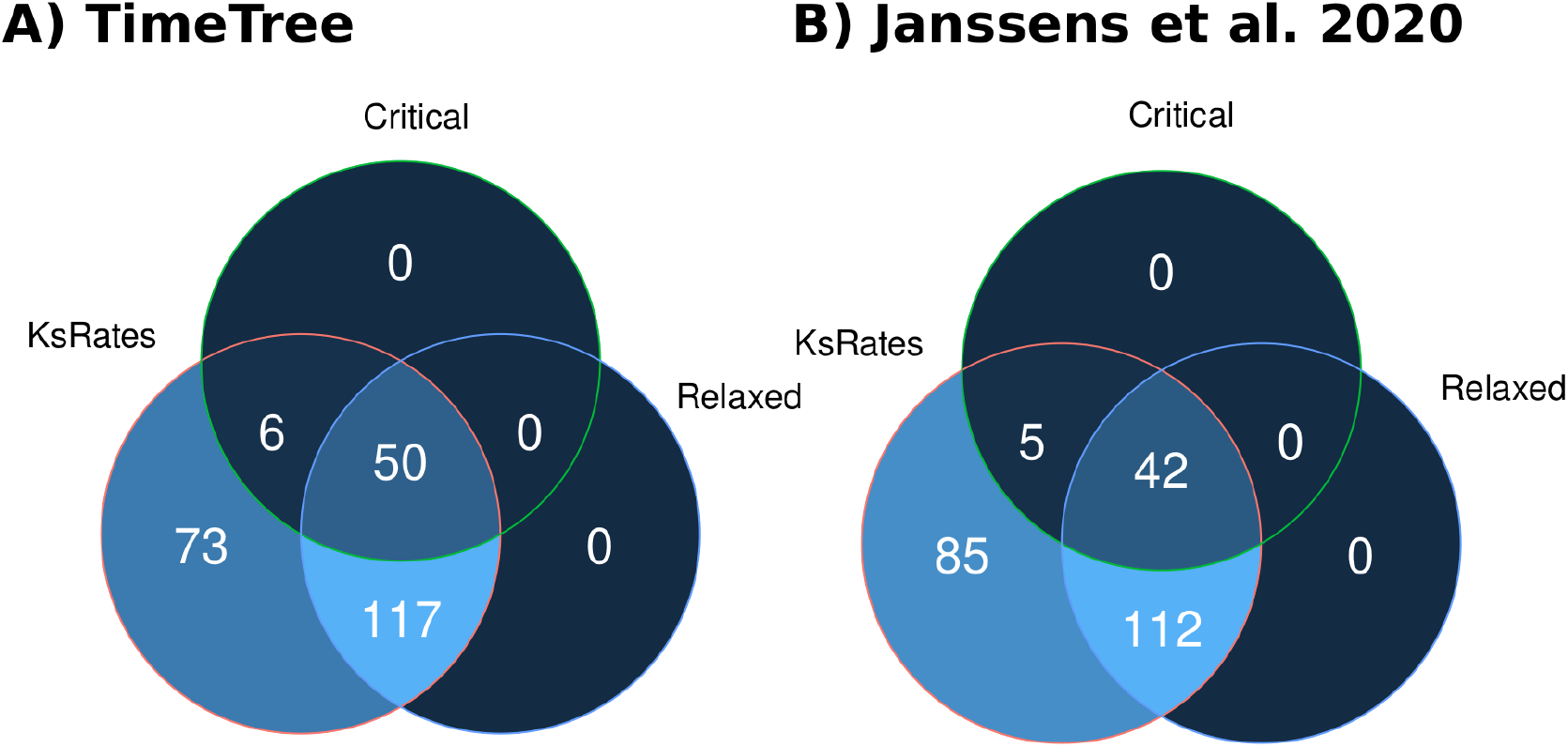
Venn diagrams of the total WGD hypotheses proposed by KsRates under the A) TimeTree and B) Janssens et al. (2020) topologies and the associated support from WHALE relaxed and Critical models.

KsRates and WHALE supported the inference of a similar number of WGDs in both the TimeTree (Kumar et al., 2017, 2022) and Janssens et al. (2020) topologies. However, the placements of these WGDs may represent different evolutionary hypotheses, as topological differences can shift which species are inferred to share a WGD event. It can be difficult to disentangle if a disagreement for the precise placement of WGD by WHALE under both topologies is due to model differences or topological non-equivalency between species trees. However, if a putative WGD is in the shared ancestry of an identical collection of species across different species trees, then we can consider the WGD placements to agree among those analyses. To make such a comparison between the TimeTree and Janssens et al. (2020) phylogenies, we compared the overlap of species descending from each WGD placement supported by WHALE. Between both sets of analyses across the two phylogenies, 124 of the hypothesized WGDs were inferred to be present in identical species, representing ∼72% and ∼78% of hypotheses supported by WHALE models in the TimeTree and Janssens et al. (2020) trees respectively. (Table S4,5). Of the remaining hypotheses in the TimeTree topology, 22 were unique and represented less than a 50% overlap with any WGD hypothesis in the Janssens et al. (2020) topology. A large fraction of these TimeTree specific hypotheses, 13, were WGD inferred to be present in only one species in the phylogeny (Table S4,5). A similar pattern was observed in the Janssens et al. (2020) topology, with the presence of 24 topology specific WGD hypotheses, 18 of which were species specific (Table S4,5). Notably, for 79% (33) of these unique species-specific WGD KsRates either did not infer WGD at all, or inferred the WGD to be shared between multiple species in the alternative tree.

### Syntenic inferences

We used additional syntenic information from the 108 genomes included in our analyses to further test hypothesized WGDs. Bi-directional intraspecific syntenic analyses were conducted using the pythonic version of MCscan (Tang et al., 2008b). MCscan classifies the syntenic depth ratio between two species, with higher syntenic depth ratios being indicative of the presence of a WGD. We visualized these inferences as a heatmap organized with the Janssens et al. (2020) topology (Fig. 3, Fig. S1). In the case of WGD presence, we expect higher syntenic depth ratios (brighter colors) in a band along the x-axis for all species sharing a given WGD and lower syntenic depth ratios (darker colors) vertically following up the y-axis in the same species. We have highlighted an example for the *Brassica* specific mesohexaploidy in Fig. 3 and supplemental Fig. 1 (Barker et al., 2009; Wang et al., 2013; Kagale et al., 2014; Jeong et al., 2016). To more formally test each of our hypothesized WGD placements, we selected all species with reference genomes up to two nodes up from the focal branch as outgroup species. For each species daughter to the hypothesized WGD position, we considered the WGD to be supported in the ancestry of that species if at least 50% of the outgroup species comparisons had a syntenic depth ratio greater than 1:1. The inference of a WGD on a given branch was considered supported if all daughter species passed the syntenic comparisons to the outgroup species. Due to limited genomic information, we were only able to test 74 and 75 WGD hypotheses under the TimeTree (Kumar et al., 2017, 2022) and Janssens et al. (2020) tree topologies (Table S4,5). Sixty-one WGD hypotheses from KsRates were supported by syntenic depth inferences in the TimeTree topology, 23 of which were not supported by either of the WHALE models (Table S4,5). Syntenic depth analyses supported 55 of the KsRates inferences of WGD in the Janssens et al. (2020) tree topology, and 18 of which were not supported by either WHALE model (Table S4,5).

**Figure 3:**
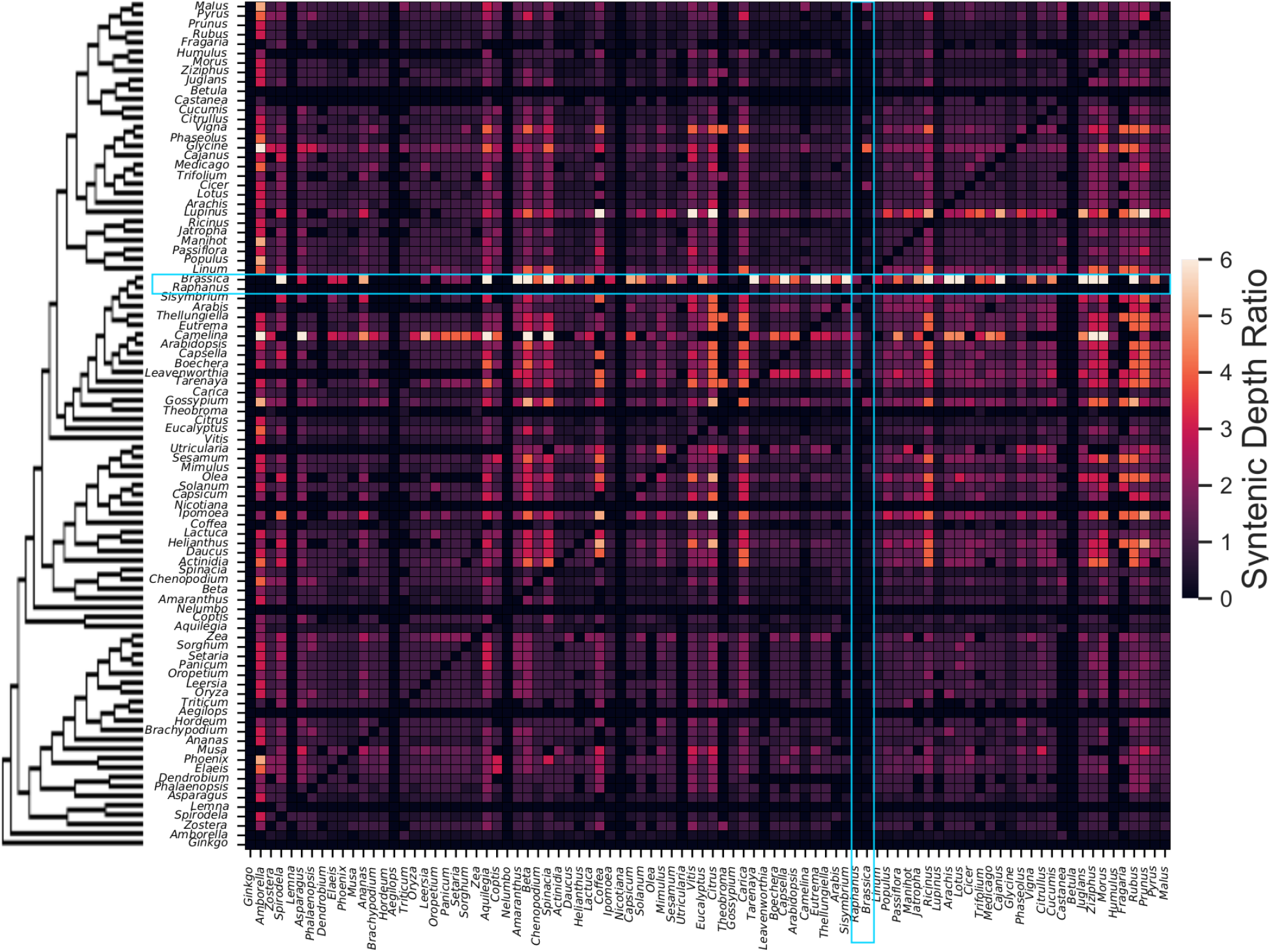
Heat map of the bi-direction syntenic depth inferences made by MCScan for each genera. The y-axis are in-group genera being analyzed ordered by the Janssens et al. (2020) tree topology. The x-axis represents the outgroup genera in the comparison arranged in the same order as the y-axis. We have highlighted an example of the expected pattern of syntenic depth ratios consistent with WGD in blue boxes. Our example shows the Brassica mesohexploidy having a consistently high syntenic depth ratio compared to outgroup species and a 1:1 or lower syntenic depth ratio when used as the outgroup for most other species.

### Merged Tree Inferences

WGDs were represented in multiple analyses due to their inherent overlap in our experimental design. We merged these inferences into the larger Janssens et al. (2020) tree to estimate the total number of supported WGD hypotheses across the entire phylogeny (Figure 4). This was not done for the TimeTree (Kumar et al., 2017, 2022) phylogeny because generating a single TimeTree with all species significantly changes the newly generated tree topology because the dating method used by TimeTree and yielded relationships inconsistent with the TimeTree phylogenies used in our WHALE analyses (unpublished data). To merge these inferences, we considered two hypothesized WGDs in different analyses as equivalent if there was only one WGD on the branches between the most recent common ancestor (MRCA) nodes of species sharing the duplication in each analysis. Following the merger, a total of 170 WGDs were inferred by KsRates and supported by WHALE, 32 of which were also supported by syntenic information (Fig. 4, Table S6). Our results are similar to prior work from 1KP (2019) that found 179 WGD in the angiosperms using K_s_ information, 38 of which were further supported by MAPS. Although the total number of WGD inferred in 1KP (2019) were similar to our inferences, their placement was not identical. Ten of these placements supported by MAPs did not have a phylogenetically equivalent branch in the Janssens et al. (2020) tree or were not tested due to KsRates not proposing their placement. Of the remaining 28 WGD hypotheses supported by MAPS we could test, 23 were also supported by WHALE models (∼82%)(Table S6). Regardless of which analysis is considered, these inferences result in an average of 3.5 WGD in the history of each angiosperm species when including WGD inferred deeper in the phylogeny than the angiosperm root (Figure 5) (Jiao et al., 2011; One Thousand Plant Transcriptomes Initiative, 2019).

**Figure 4:**
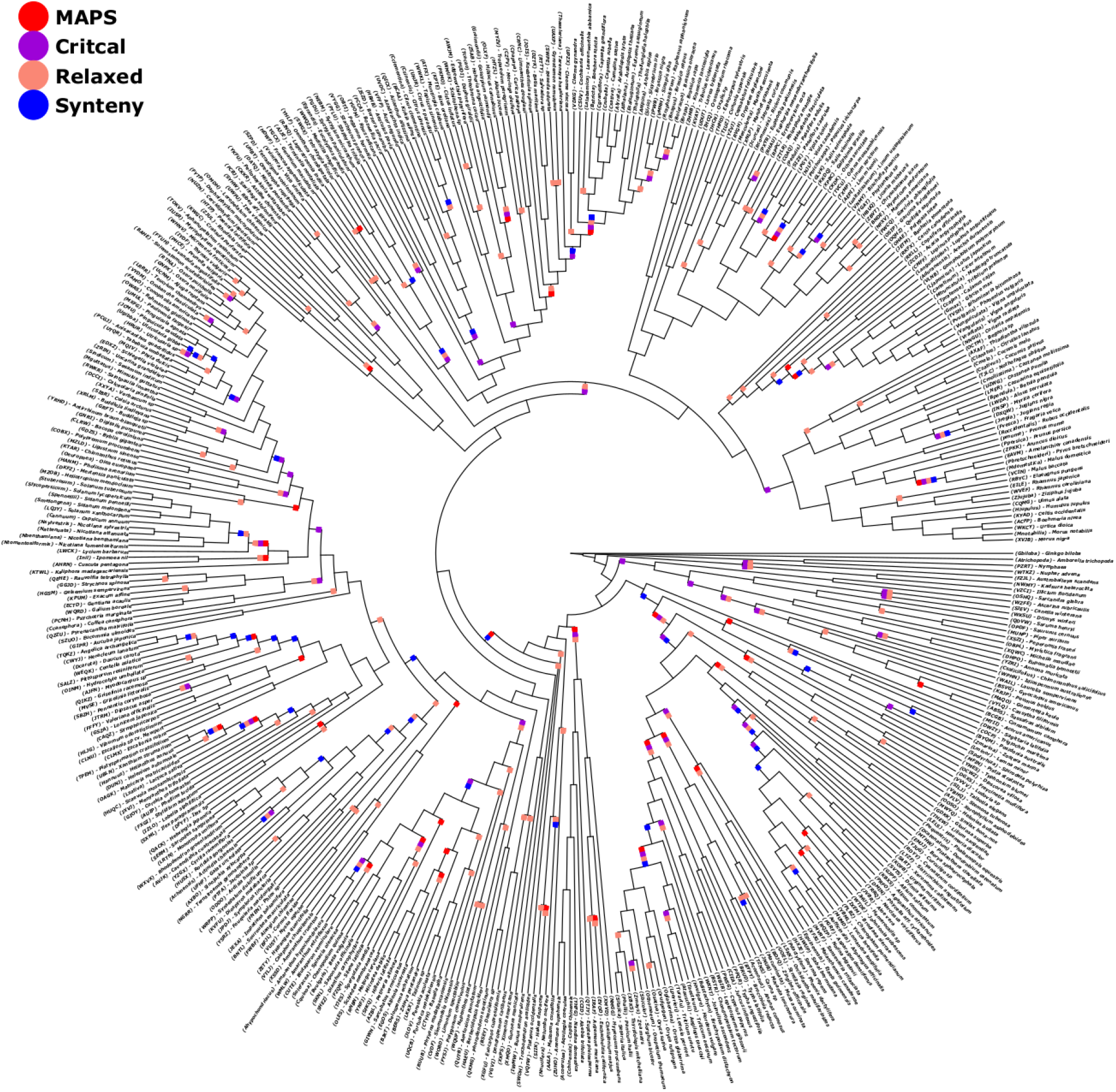
Final merged Janssens et al. (2020) phylogeny annotated with hypothesized WGD placements supported by synteny, WHALE models, or MAPS. MAPS inferences were based on the equivalent MRCA for species inferred to share the proposed WGD in the 1KP 2019 analysis. Annotations are as follows: MAPS (red), WHALE Critical (purple), relaxed model (peach), and the syntenic depth test (blue).

**Figure 5:**
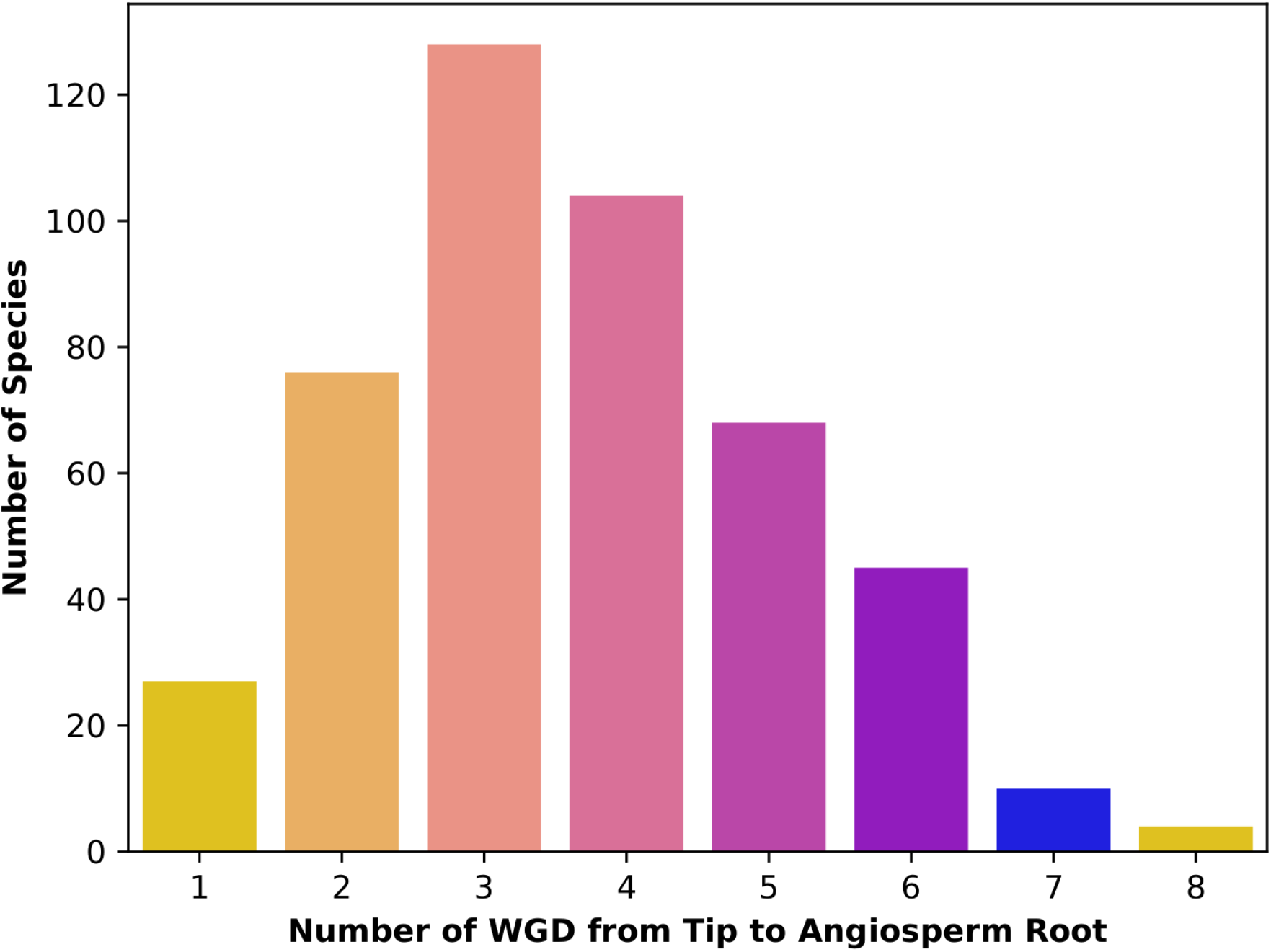
Histogram of the rounds of WGD in the history of each species included in the Janssens et al. topology. Rounds of WGD are based on the WGD hypotheses supported by WHALE models plus the AMBOα duplication shared by all angiosperms (Jiao et al., 2011; One Thousand Plant Transcriptomes Initiative, 2019).

## DISCUSSION

Our inferences of WGDs using state of the art methods and two different species tree topologies were largely consistent with previous analyses of WGDs across the angiosperms. We observed approximately 170 WGDs across our dataset, compared to 179 reported in the 1KP (2019) analysis. These observations result in an average of ∼3.5 WGD in the history of angiosperm species regardless of method or treetoplogy used. Although KsRates, WHALE, and the 1KP (2019) analysis largely agreed on the relative number of WGDs among lineages, their precise placement within the angiosperm phylogeny was variable. Approximately 80% of WGD hypotheses were inferred to be shared by the same species under both the TimeTree (Kumar et al., 2017, 2022) and Janssens et al. (2020) phylogenies. The remaining 20% of WGD hypotheses unique to each species tree were largely a result of alternative tree topologies altering hypothesis generation. Given this degree of uncertainty, and regardless of which WGD inference method employed, future researchers should carefully consider the limitations of methods used to produce hypothesized phylogenies, and placing hypothesized WGDs into those contexts.

We observed that KsRates proposed an inflated number of WGDs (∼245), many of which represented alternative hypotheses for particular WGDs. These alternative placements were largely due to differences in how synonymous substitution rate heterogeneity was estimated using different reference species. Additional variation in WGD placement was also caused by multiple or alternative mixture models fit to the paralog distribution. Mixture models are known to overestimate the number of components (Tiley et al., 2018; Sensalari et al., 2022) and can be biased by peak shape variation due to gene conversion (Qiao et al., 2019; Sensalari et al., 2022) and allopolyploidy (Marcet-Houben and Gabaldón, 2015; Sensalari et al., 2022). For example, within the Brassicales we observed a difference of ∼0.3 in the median K_s_ of the ARTHɑ WGD compared to previous reports (Barker et al., 2009; Wang et al., 2013; Kagale et al., 2014; Jeong et al., 2016). Even minor variation in the estimate of mean K_s_ of ortholog and paralog peaks can result in shifting WGD hypothesis up or down branches. Continuing our example in the Brassicales, our K_S_ inferences using the Janssens et al. (2020) phylogeny were largely consistent with prior reports of WGDs within this group (Barker et al., 2009; Wang et al., 2013; Kagale et al., 2014; Jeong et al., 2016). However, the TimeTree (Kumar et al., 2017, 2022) topology was different from previously inferred phylogenies (Barker et al., 2009; Edger et al., 2018; Mabry et al., 2020; Walden et al., 2020) and resulted in two alternative placements for the ARTHβ duplication (Barker et al., 2009; Wang et al., 2013; Kagale et al., 2014; Jeong et al., 2016; Mabry et al., 2020) that were also supported by syntenic and WHALE analyses. These results indicate that although K_s_ methods are an accessible and powerful tool for assessing the presence of WGD within species, they should be used carefully when placing WGDs on a phylogeny.

Bayesian gene tree-species tree reconciliation approaches for estimating the rates of gene duplication and loss are relatively robust to topological variation when initial WGD hypothesis generation is accurate. We used WHALE to remove a large fraction of alternative hypothesized WGD placements made by KsRates, resulting in a similar number of WGDs (∼170) inferred compared to previous reports (One Thousand Plant Transcriptomes Initiative, 2019). However, WHALE models often disagreed on the precise placement of WGDs under different species tree hypotheses. Our results suggest that duplication and loss rates among angiosperms are highly variable, and models that account for rate heterogeneity both among and within branches are less sensitive to topological variation than those that do not, consistent with prior reports (Zwaenepoel and Van de Peer, 2019a). In contrast, when holding duplication and loss rates constant within branches, WHALE models underestimated the number of WGD by almost 70% and often rejected well known WGDs such as the ARTHɑ WGD in the Brassicales (however supported in Zwaenepoel and Van de Peer, 2019a). Even when using the more robust relaxed model, 20% of hypothesized WGD placements were species tree specific, 76% of which were unique to single species. WHALE models also largely agreed with prior inferences from MAPS in the 1KP (2019) analyses. However, much like disagreements between WHALE models, a large fraction (∼67%) of disagreements with previous MAPS analyses were due to species tree topology differences between our study and the 1KP (2019) analyses.

We used syntenic analyses as an alternative method to testing hypothesized WGD placements. The availability of genomic data allowed us to test a total of 149 WGD hypotheses across our analyses, 116 of which were supported by syntenic depth inferences. Notably, a large fraction of these WGD hypotheses (∼35%) were not supported by either WHALE model. Three of these hypotheses were also supported by MAPs, however others may be due to limitations of our syntenic depth heuristic. Our syntenic analyses often supported both alternative hypotheses for a given WGD, as observed for the ARTHβ WGD in the A1 analysis under the TimeTree topology (Kumar et al., 2017, 2022). Many tools already exist for analyzing complex syntenic data, however they also require *a priori* knowledge of the shared WGD history among species (Tang et al., 2011; Parey et al., 2020; Siddiqui and Conant, 2023). Given the added power syntenic analysis can give to disentangling complex WGD histories, our analyses and other WGD focused syntenic tools highlight the need for formal tests to infer the shared polyploid ancestry of species using synteny.

The topology of species tree hypotheses are arguably more impactful on WGD inference than the inference method. Some methods used in this study were more robust to variation than others, however the most frequent cause of disagreements among methods in this study, as well as the results of prior work (One Thousand Plant Transcriptomes Initiative, 2019), were topological differences between species trees. Here we have only evaluated WGD inferences using two alternative topologies, making it difficult to discern how branching patterns and branch lengths impact WGD inferences respectively. Branch lengths directly determine both synonymous rate corrections and estimated rates of gene duplication and loss (Zwaenepoel and Van de Peer, 2019a; Sensalari et al., 2022), however it is unclear how sensitive WGD inferences made using these methods are to variation in branch lengths between alternative phylogenies. Error in the inferred evolutionary relationships among species likely represents an even larger challenge to WGD inference. It is well known that paralogy reconciliation and species tree inferences influence each other (Shi, 2013; Struck, 2013; Smith and Hahn, 2022; Xiong et al., 2022), leading some to estimate them concurrently (Boussau et al., 2013; Szöllősi et al., 2014). In the case when gene trees are reconciled with a prior curated species tree, the estimation of duplication and loss rates are constrained to the species tree topology and inherit any errors therein. Inferring accurate species trees has long been a central challenge in evolutionary biology (see Anderson et al. 2012, Szöllősi et al. 2014, and Morel et al. 2022), and WGD inferences ultimately depend on a solid phylogenetic foundation. A firmer understanding of the sensitivity of different WGD inference methods to different species tree inference errors is important for future efforts to untangle the complex evolutionary history of ancient WGDs across eukaryotes.

In this study we explored the distribution of ancient WGDs across the angiosperm phylogeny. We found that a variety of current methods are robust in detecting the presence of WGDs across lineages. However, we observed significant sensitivity to species tree topology when placing WGDs within the angiosperm phylogeny. Given that species trees are hypotheses themselves, to which we map WGD hypotheses onto, it is critical to carefully consider the uncertainty introduced by different WGD inference methods. Many fundamental questions in polyploid evolution are beginning to be addressed with the increased availability of genomic data (Jiao et al., 2011; Vanneste et al., 2014; Tank et al., 2015; Soltis and Soltis, 2016; Landis et al., 2018; Smith et al., 2018; Bohutinska et al., 2020; Hao et al., 2021; Li et al., 2021; Qi et al., 2021; Marchant et al., 2022; Qiao et al., 2022; Almeida-Silva and Van de Peer, 2023; Bhadra et al., 2023; Blischak et al., 2023). Accurate inferences for the shared polyploid ancestry among angiosperm species are necessary when forming predictions of polyploid plant evolution. For these reasons, future work characterizing the impacts of variation in species tree topology on methods for inferring ancient WGDs is necessary. Methods such as syntenic analyses may offer additional evidence for disentangling complex WGD histories, however more formal methods are required. As we expand the availability of data across the tree of life, the goal of accurate WGD inferences will likely reveal new insights into the profound impacts polyploidy has had on the evolution of plants and other lineages alike.

## Supporting information

Appendiz S2

## ACKNOWLEDGEMENTS

The authors would like to thank the University of Arizona TRIF, UITS, Research, Innovation, and Impact (RII), and Research Technologies departments for supporting the High Performance Computing (HPC) resources used to conduct our analyses. We would also like to thank Dr. Justin Conover, Dr. Sylvia Kinosian, Seongyeon Kang, and Qiuyu Jiang for their feedback on this manuscript and during the development of the project. Finally, we thank Dr. Katrina Dlugosch for assistance with running analyses on the HPC.

## AUTHOR CONTRIBUTIONS

M.T.W.M and M.S.B. designed the project, analyzed the data, and wrote the manuscript. G.F. assisted with preparing the Janssens et al. (2020) sub-analyses.

## DATA AVAILABILITY STATEMENT

Data and scripts used to analyze these data are available at the following git repository: https://gitlab.com/barker-lab/ajb_special_issue_2023.git.

