## Appendiz S2 for "Species Tree Topology Impacts the Inference of Ancient Whole-Genome Duplications Across the Angiosperm Phylogeny"

#### Appendix S2 - Supporting Figures - S1-S85

**Figure S1:** Heat map of the bi-direction syntenic depth inferences made by MCScan.

**Figure S2-85:** Phylogenies for each analysis under the TimeTree and Janssens et al. (2020) phylogenies.

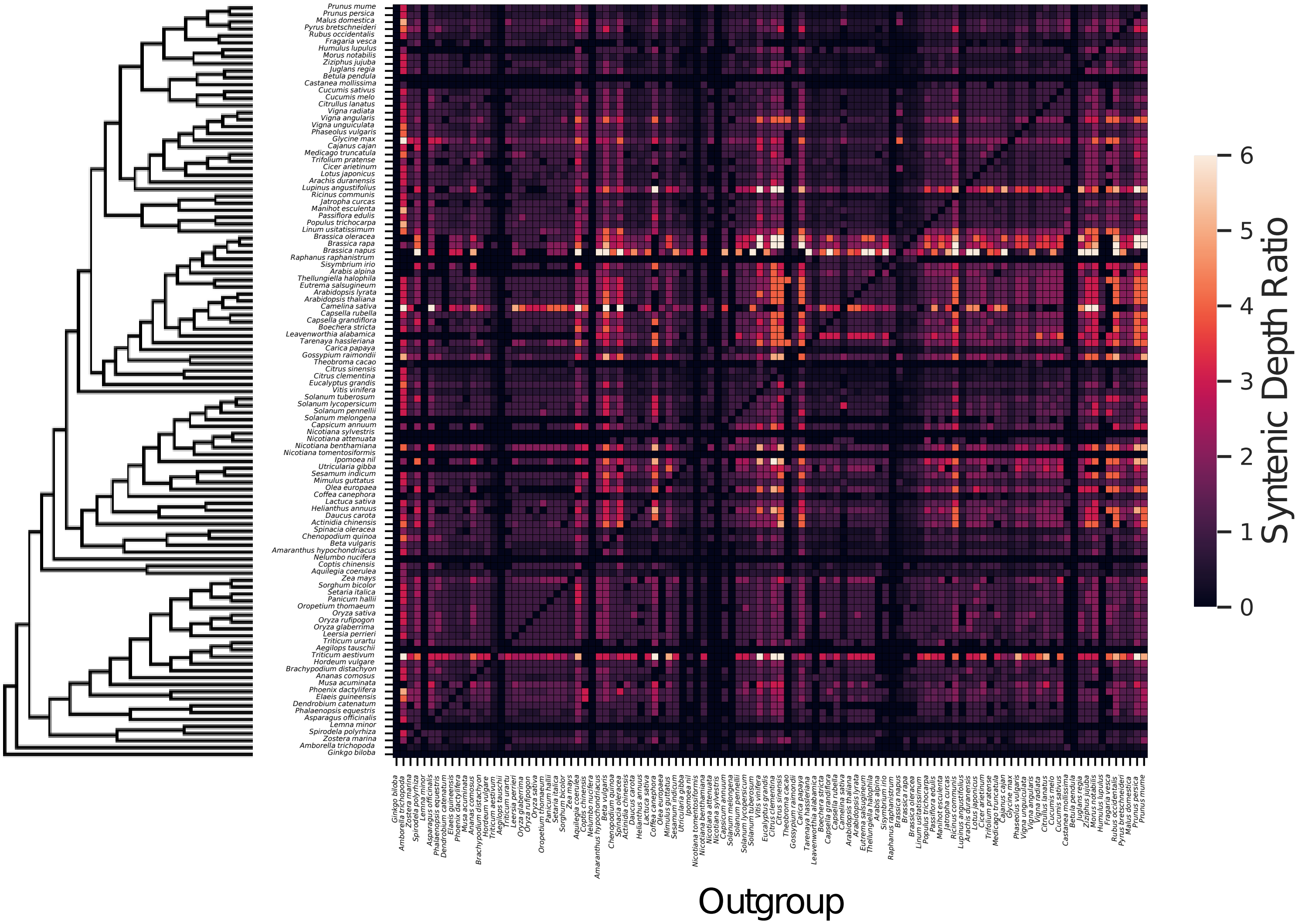


**Figure S1:** Heat map of the bi-direction syntenic depth inferences made by MCScan for each species. The y-axis are in-group species being analyzed ordered by the Janssens et al. (2020) tree topology. The x-axis represents the outgroup species in the comparison arranged in the same order as the y-axis. We have highlighted an example of the expected pattern of syntenic depth ratios consistent with WGD in blue boxes. Our example shows the Brassica mesohexploidy having a consistently high syntenic depth ratio compared to outgroup species and a 1:1 or lower syntenic depth ratio when used as the outgroup for most other species.

### A1 TimeTree

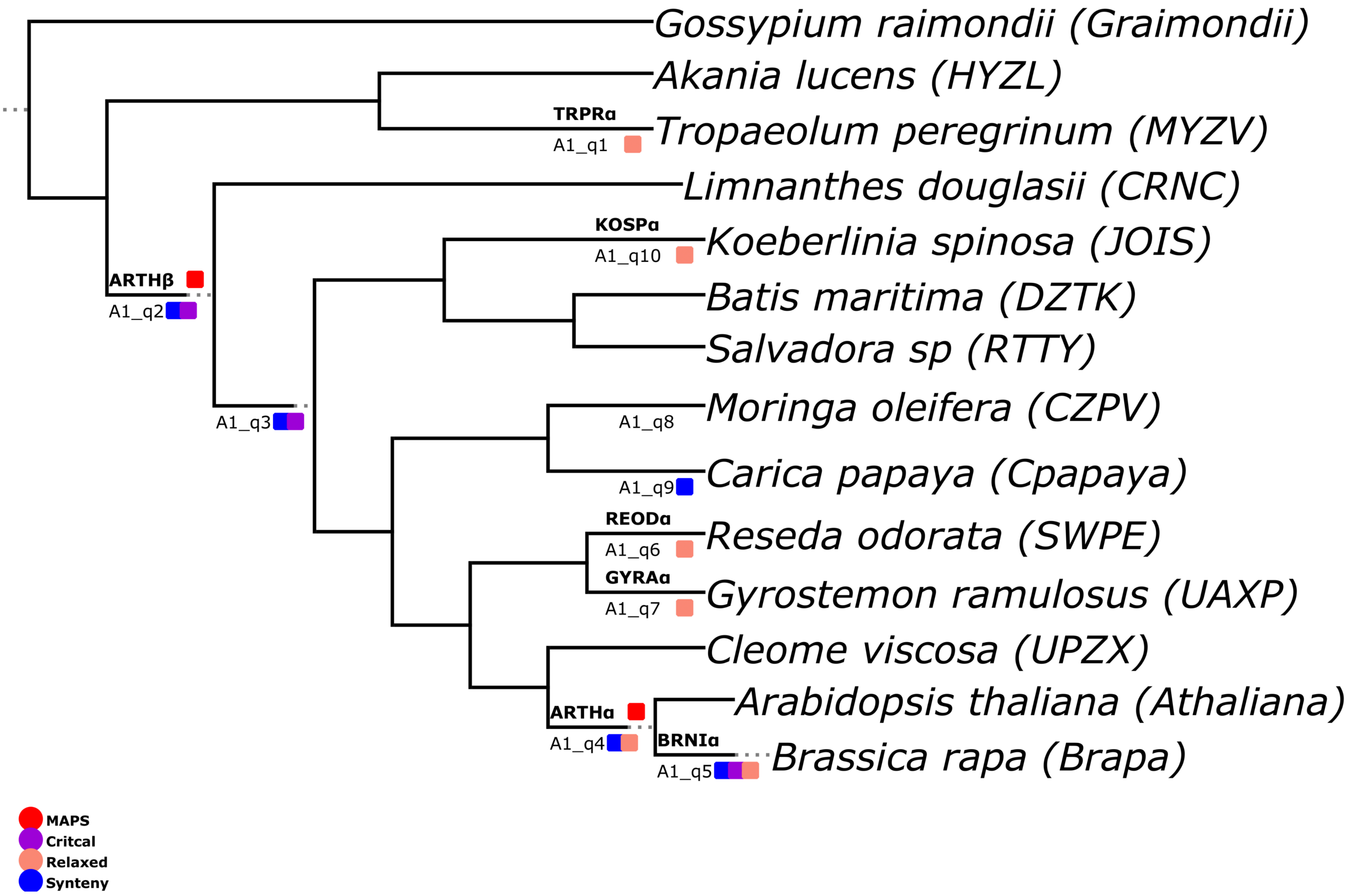


### Figure S2

### A2 TimeTree

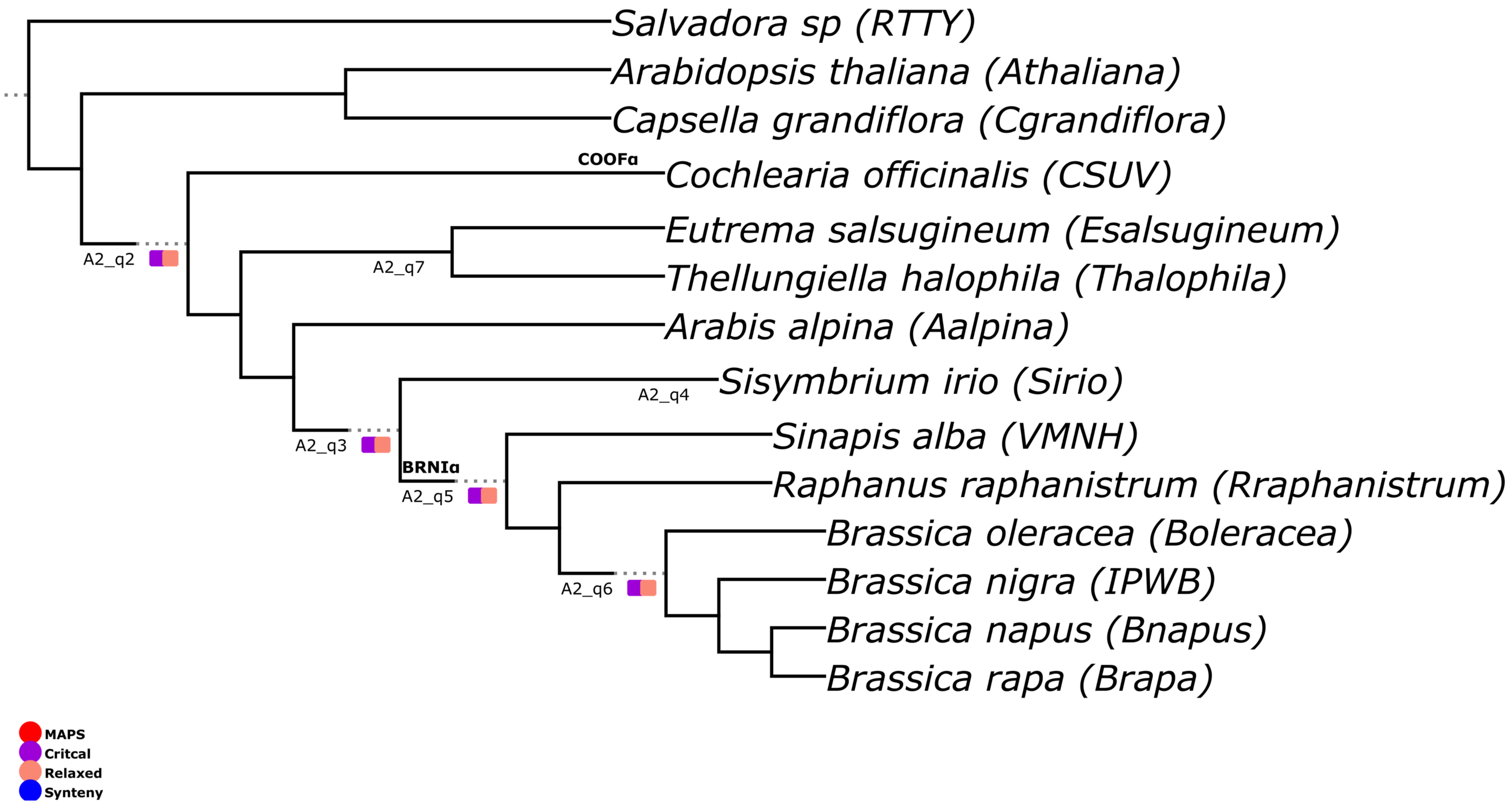


### Figure S3

### A3 TimeTree

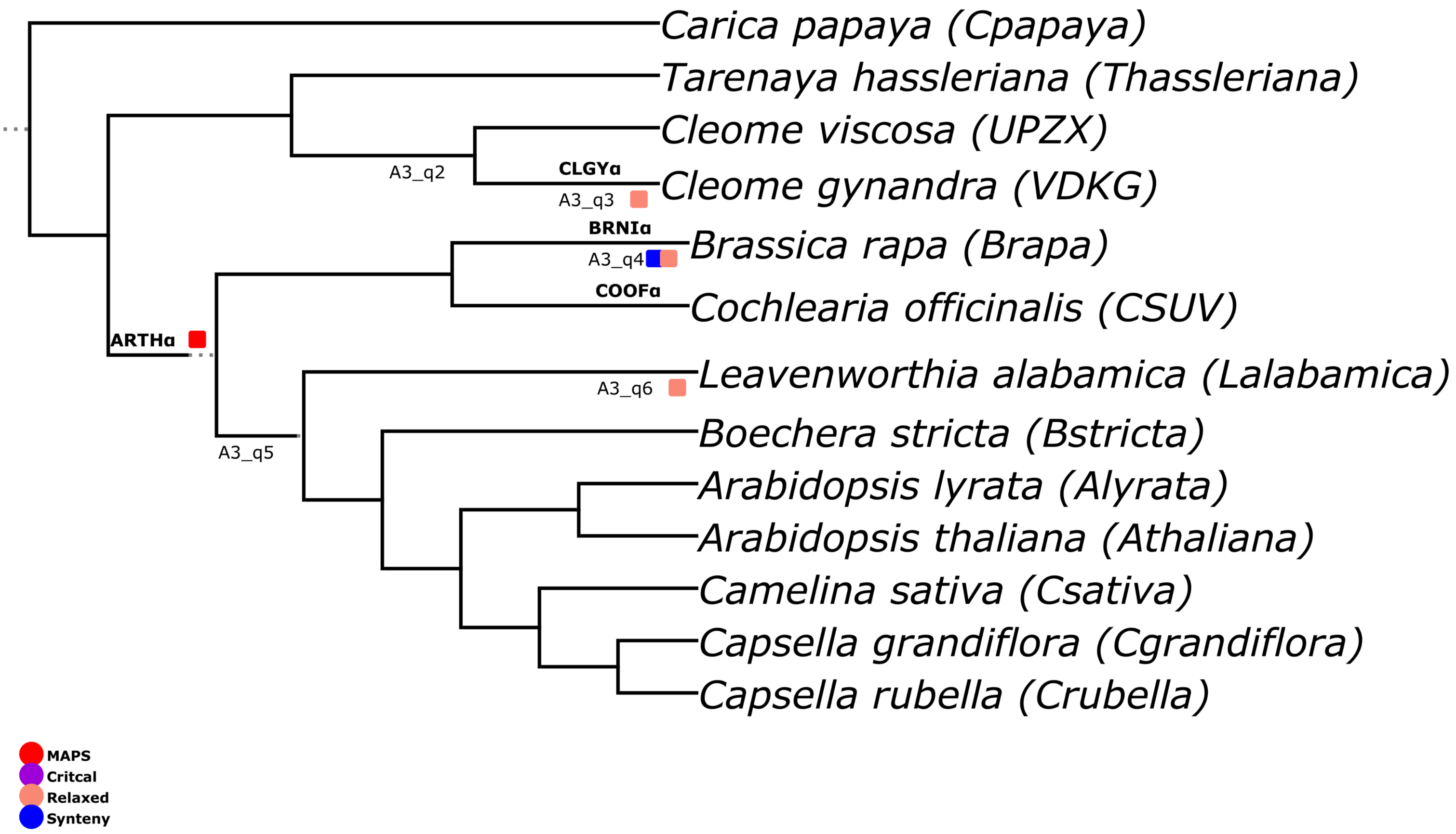


### Figure S4

### B1 TimeTree

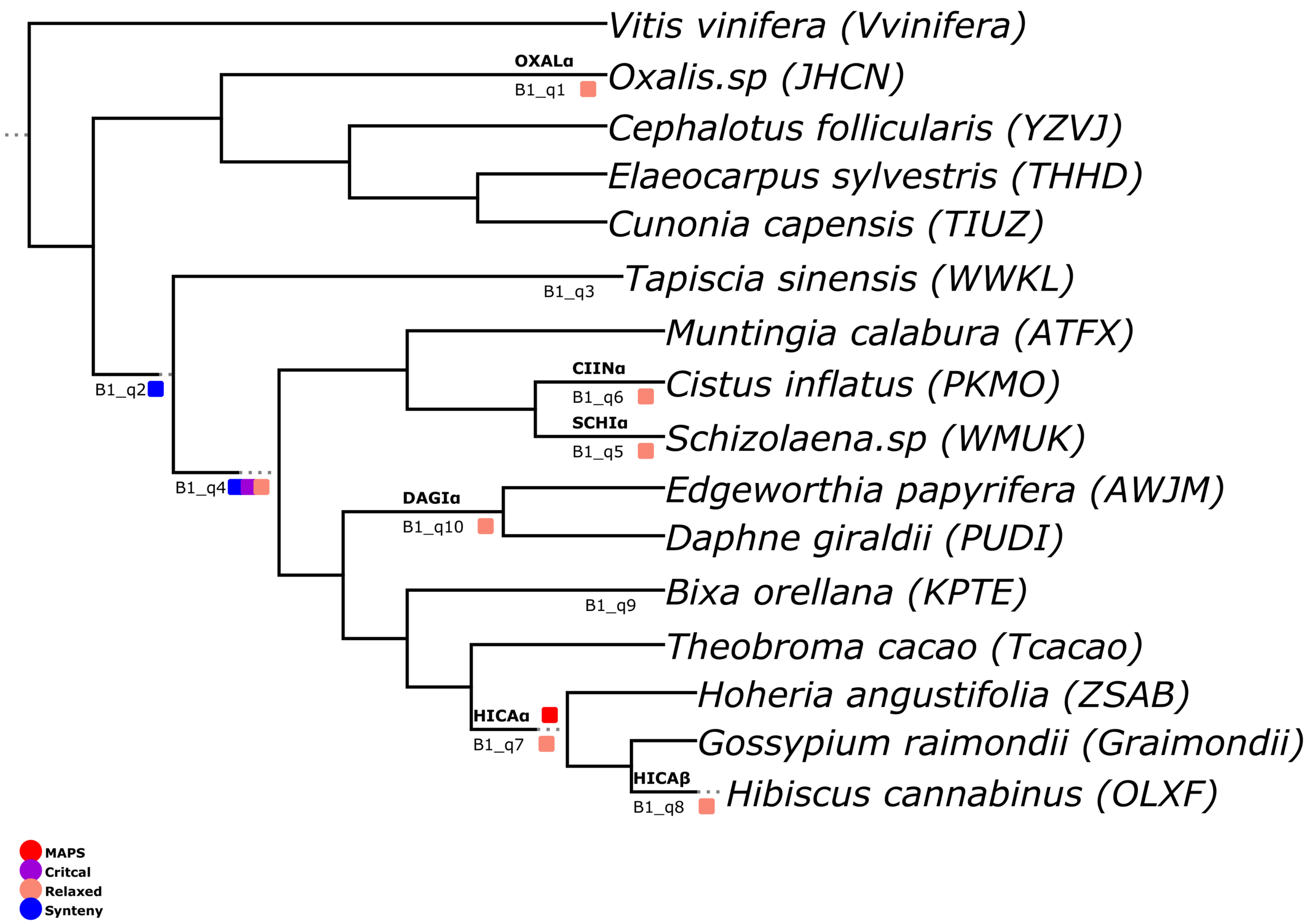


### Figure S5

### B2 TimeTree

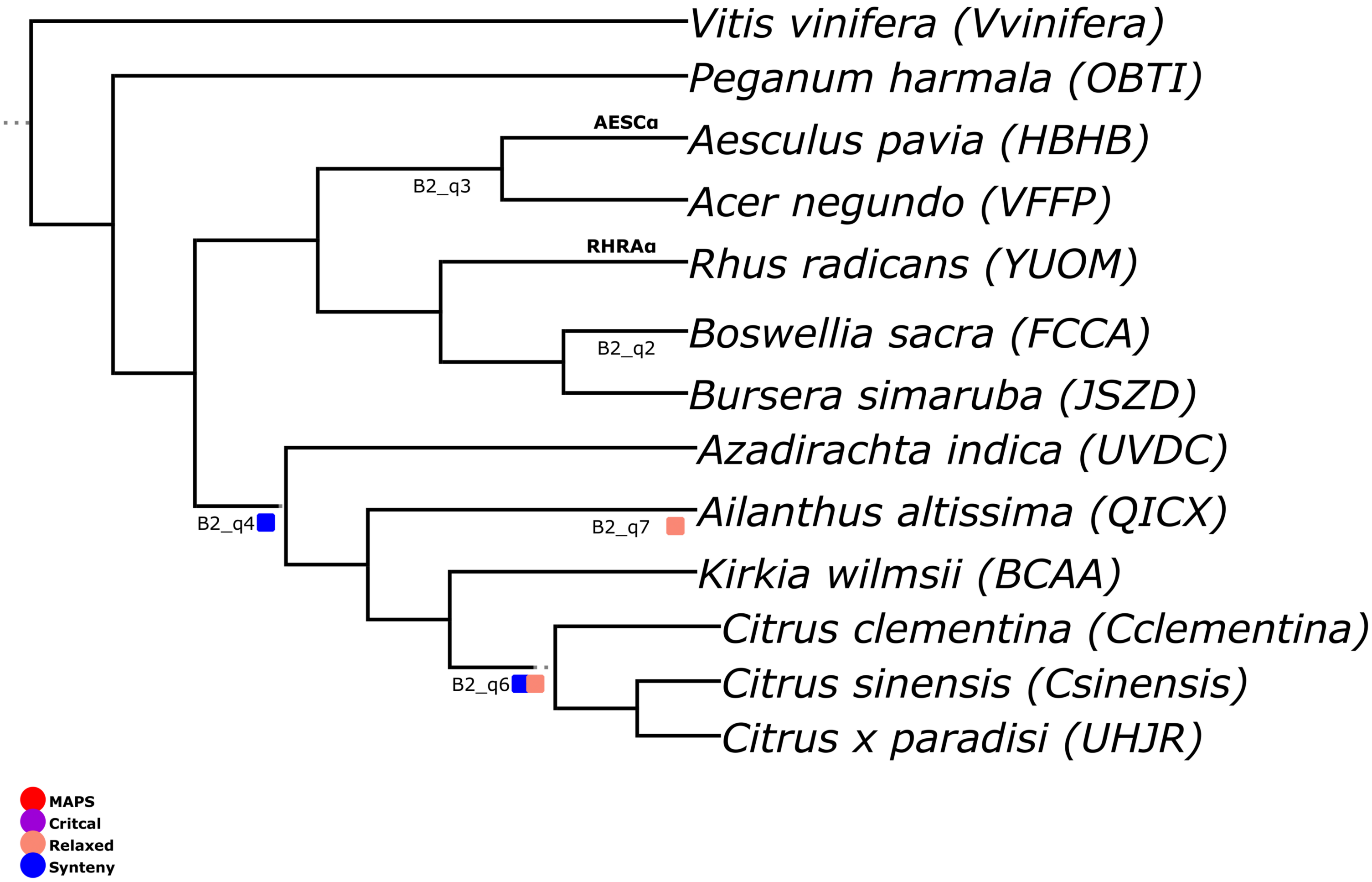


### Figure S6

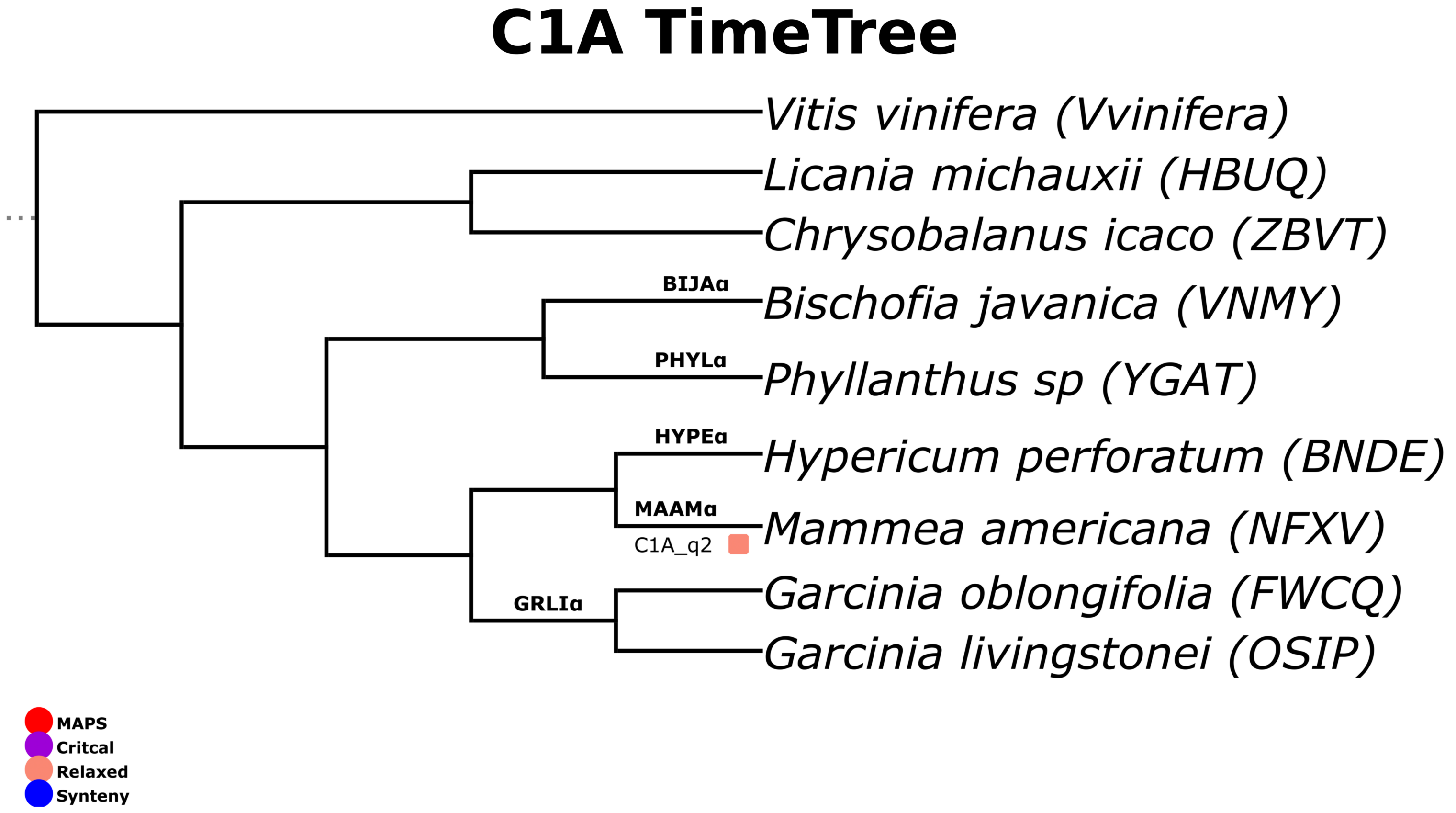
C1A TimeTree
Vitis vinifera (Vvinifera)
Licania michauxii (HBUQ)
Chrysobalanus icaco (ZBVT)
BIJAα
Bischofia javanica (VNMY)
PHYLα
Phyllanthus sp (YGAT)
HYPEα
Hypericum perforatum (BNDE)
MAAMα
Mammea americana (NFXV)
C1A_q2
GRLIα
Garcinia oblongifolia (FWCQ)
Garcinia livingstonei (OSIP)
MAPS
Critcal
Relaxed
Synteny

**Figure S7**

### C1B TimeTree

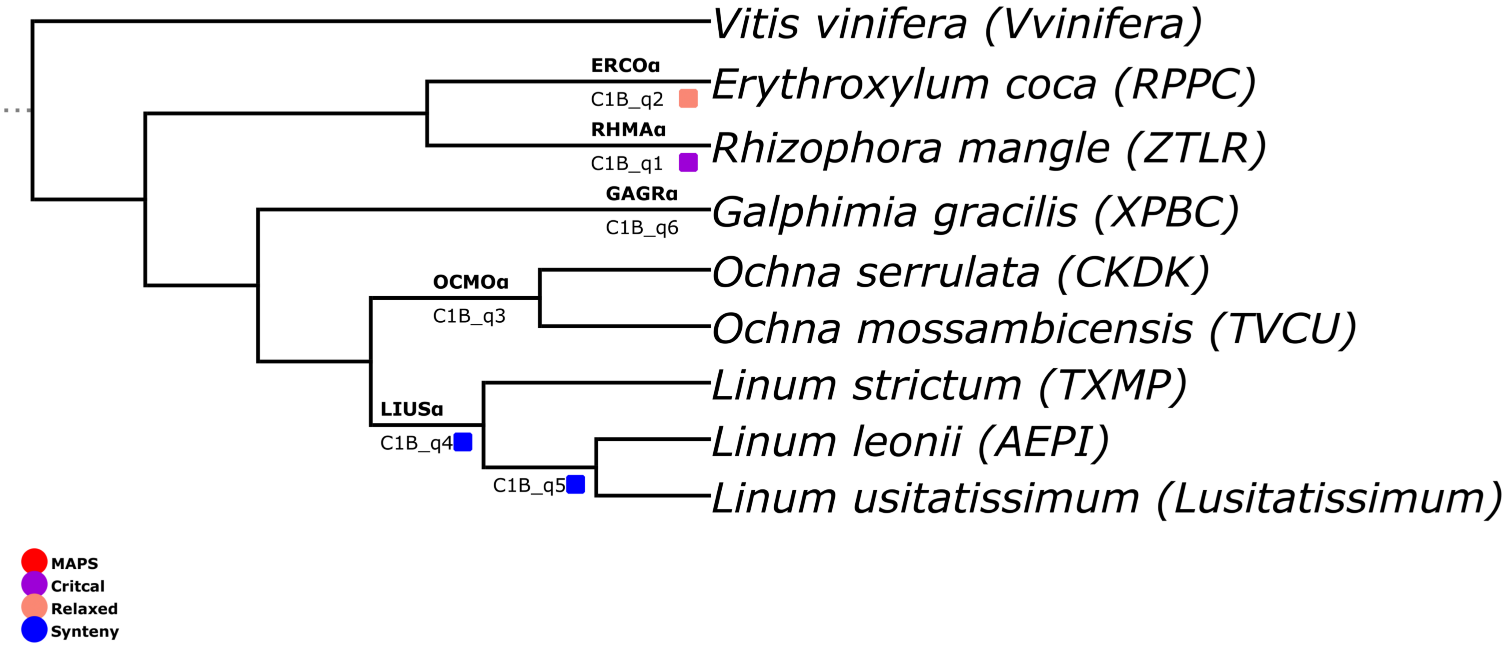


### Figure S8

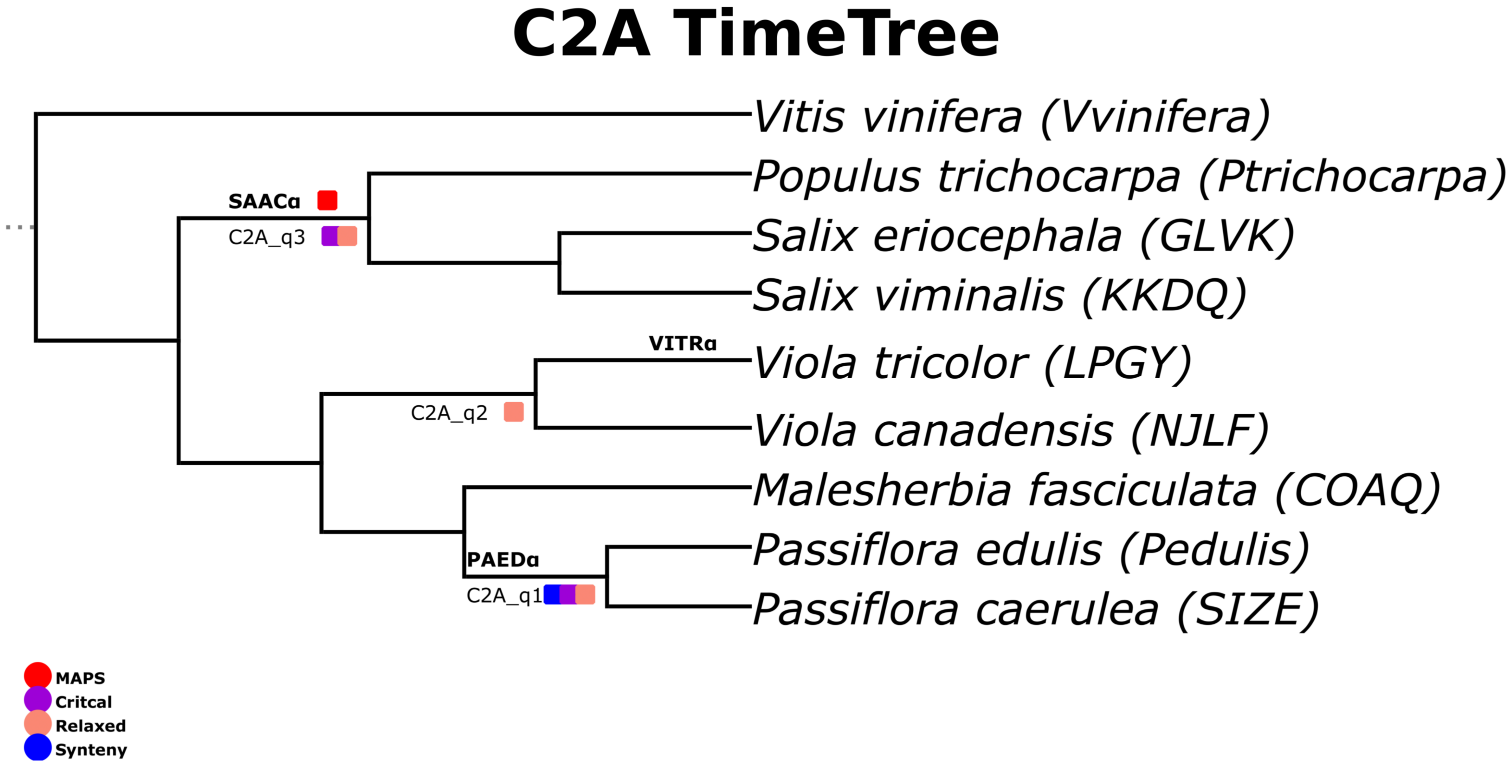
C2A TimeTree
Vitis vinifera (Vvinifera)
Populus trichocarpa (Ptrichocarpa)
Salix eriocephala (GLVK)
Salix viminalis (KKDQ)
Viola tricolor (LPGY)
Viola canadensis (NJLF)
Malesherbia fasciculata (COAQ)
Passiflora edulis (Pedulis)
Passiflora caerulea (SIZE)
SAACα
C2A_q3
VITRα
C2A_q2
PAEDα
C2A_q1
MAPS
Critcal
Relaxed
Synteny

### Figure S9

### C2B TimeTree

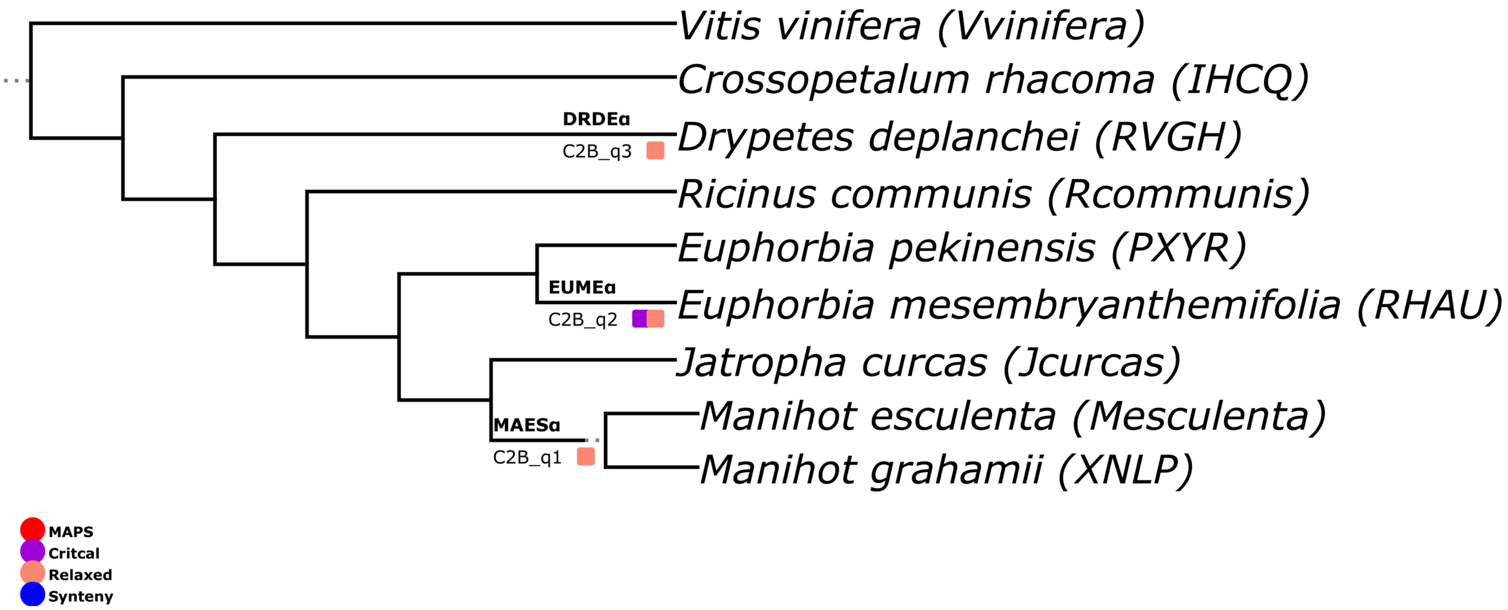


**Figure S10**

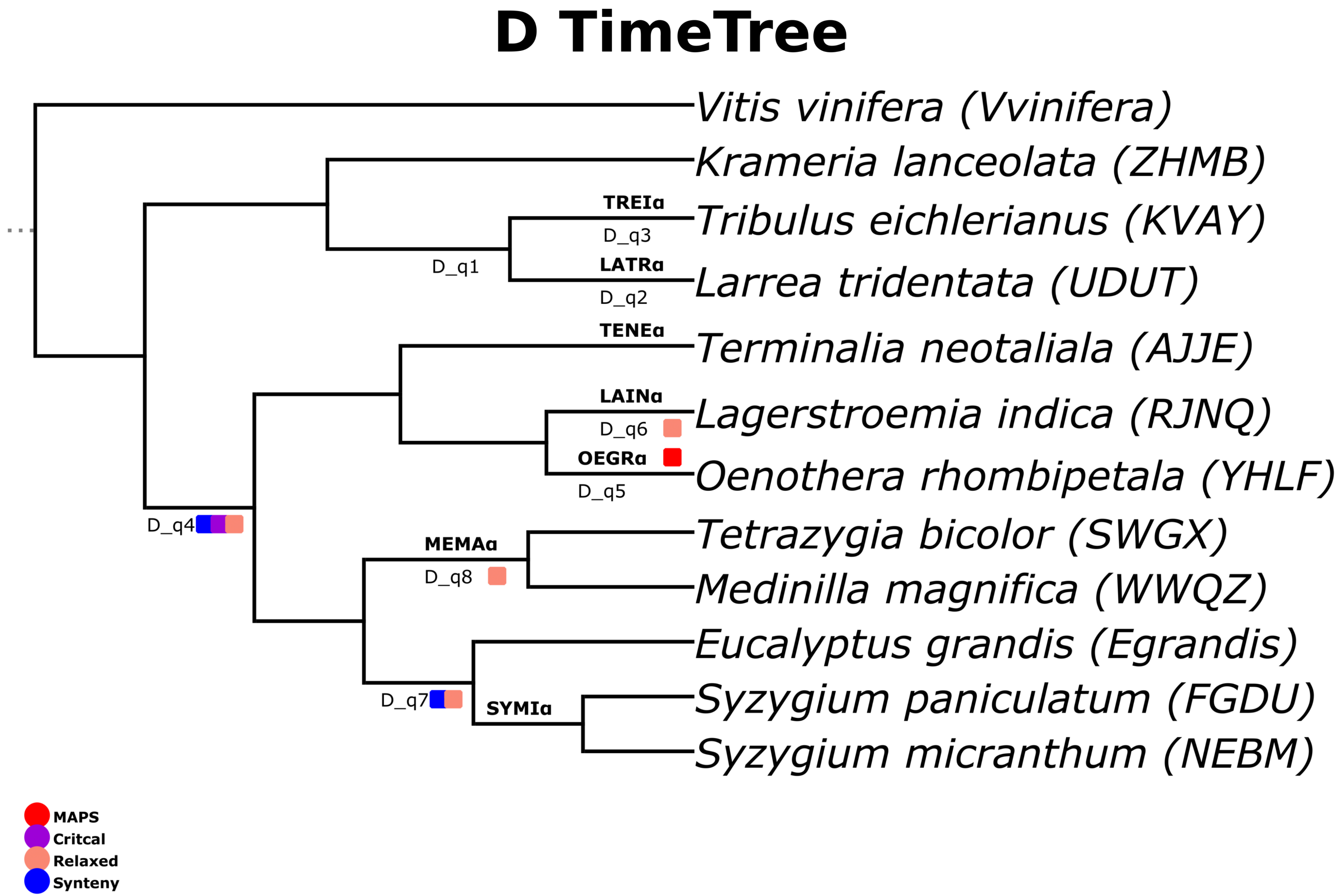

D TimeTree
Vitis vinifera (Vvinifera)
Krameria lanceolata (ZHMB)
TREIα
D_q3
Tribulus eichlerianus (KVAY)
D_q1
LATRα
Larrea tridentata (UDUT)
D_q2
TENEα
Terminalia neotaliala (AJJE)
LAINα
Lagerstroemia indica (RJNQ)
D_q6
OEGRα
Oenothera rhombipetala (YHLF)
D_q5
D_q4
Tetrazygia bicolor (SWGX)
MEMAα
D_q8
Medinilla magnifica (WWQZ)
Eucalyptus grandis (Egrandis)
D_q7
SYMIα
Syzygium paniculatum (FGDU)
Syzygium micranthum (NEBM)
MAPS
Critcal
Relaxed
Synteny


### Figure S11

### E1 TimeTree

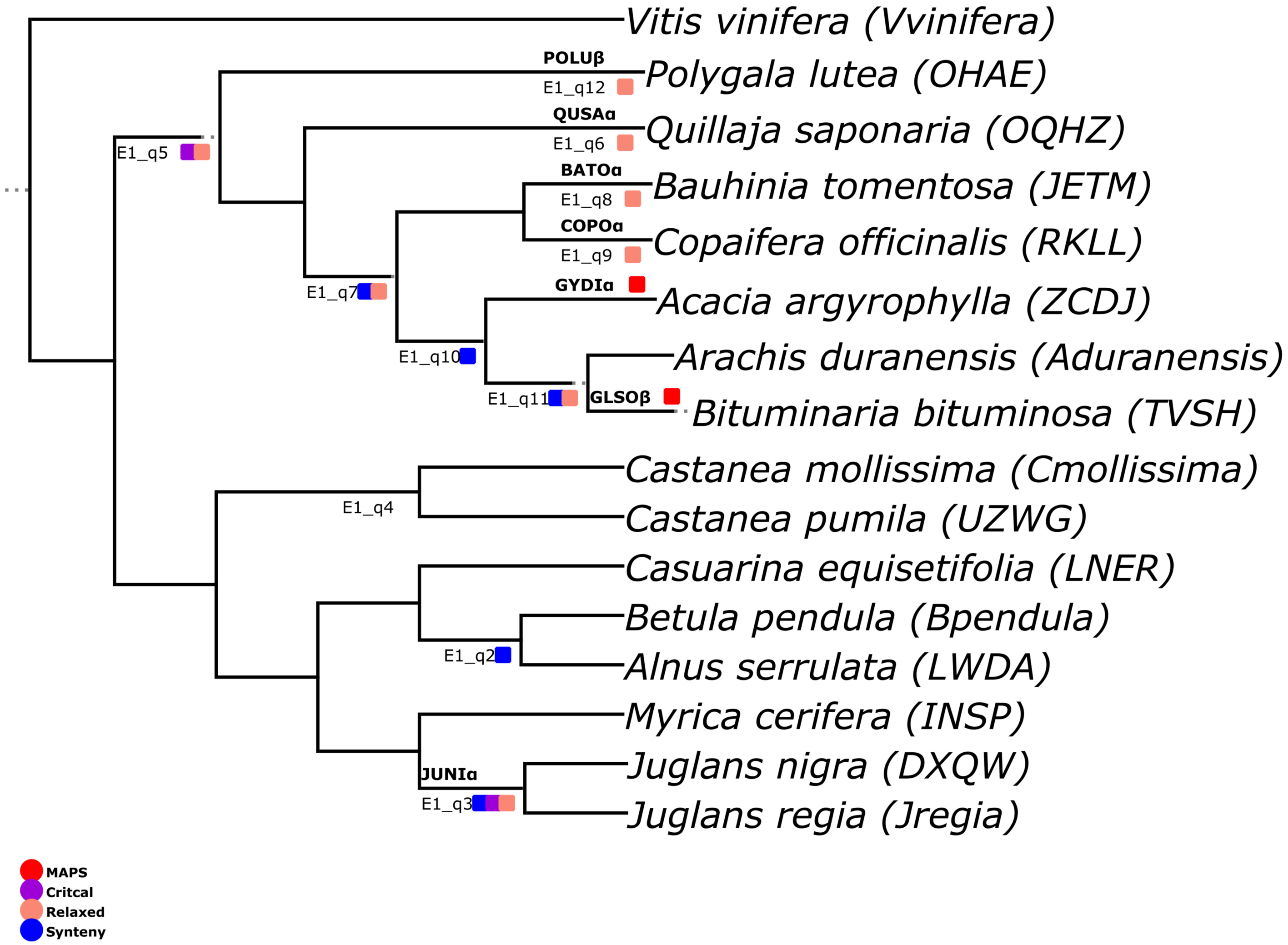


### Figure S12

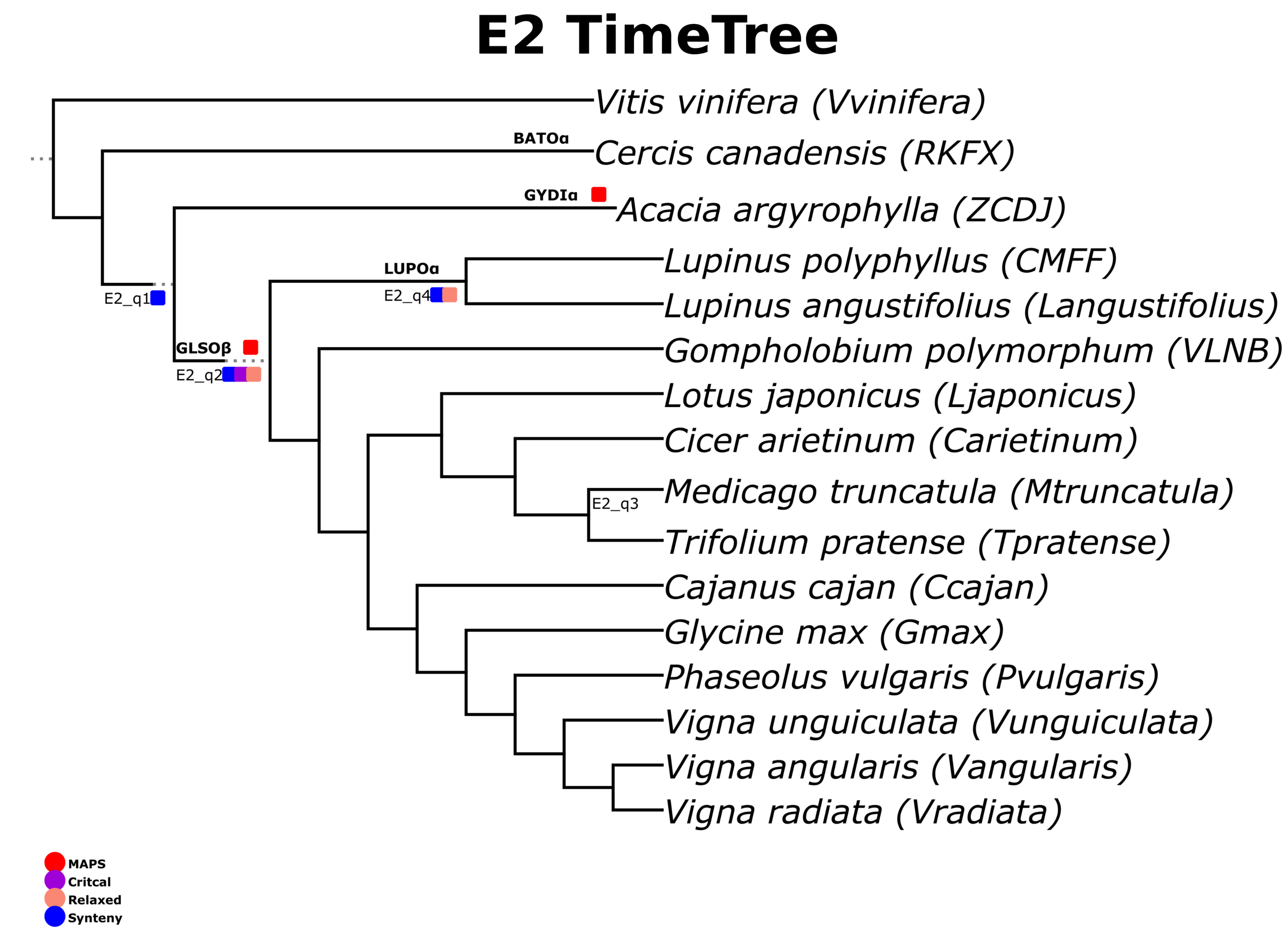
E2 TimeTree
Vitis vinifera (Vvinifera)
BATOα
Cercis canadensis (RKFX)
GYDIα
Acacia argyrophylla (ZCDJ)
LUPOα
Lupinus polyphyllus (CMFF)
E2_q4
Lupinus angustifolius (Langustifolius)
E2_q1
GLSOβ
E2_q2
Gompholobium polymorphum (VLNB)
Lotus japonicus (Ljaponicus)
Cicer arietinum (Carietinum)
E2_q3
Medicago truncatula (Mtruncatula)
Trifolium pratense (Tpratense)
Cajanus cajan (Ccajan)
Glycine max (Gmax)
Phaseolus vulgaris (Pvulgaris)
Vigna unguiculata (Vunguiculata)
Vigna angularis (Vangularis)
Vigna radiata (Vradiata)
MAPS
Critcal
Relaxed
Synteny

### Figure S13

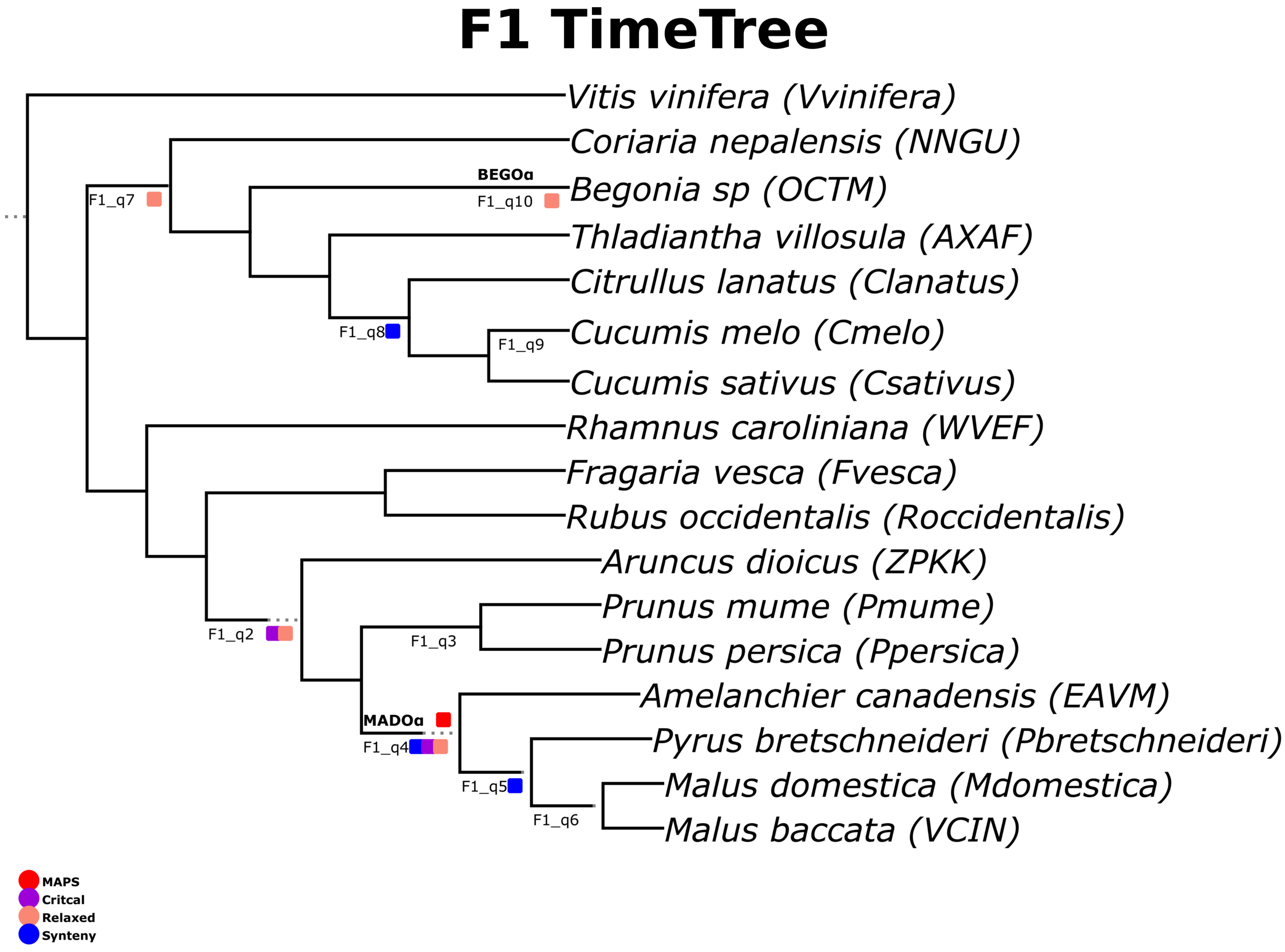
F1 TimeTree
Vitis vinifera (Vvinifera)
Coriaria nepalensis (NNGU)
BEGOα
Begonia sp (OCTM)
F1_q10
F1_q7
Thladiantha villosula (AXAF)
Citrullus lanatus (Clanatus)
Cucumis melo (Cmelo)
F1_q8
F1_q9
Cucumis sativus (Csativus)
Rhamnus caroliniana (WVEF)
Fragaria vesca (Fvesca)
Rubus occidentalis (Roccidentalis)
Aruncus dioicus (ZPKK)
Prunus mume (Pmume)
F1_q2
F1_q3
Prunus persica (Ppersica)
Amelanchier canadensis (EAVM)
MADOα
F1_q4
Pyrus bretschneideri (Pbretschneideri)
F1_q5
Malus domestica (Mdomestica)
F1_q6
Malus baccata (VCIN)
MAPS
Critcal
Relaxed
Synteny

### Figure S14

### F2 TimeTree

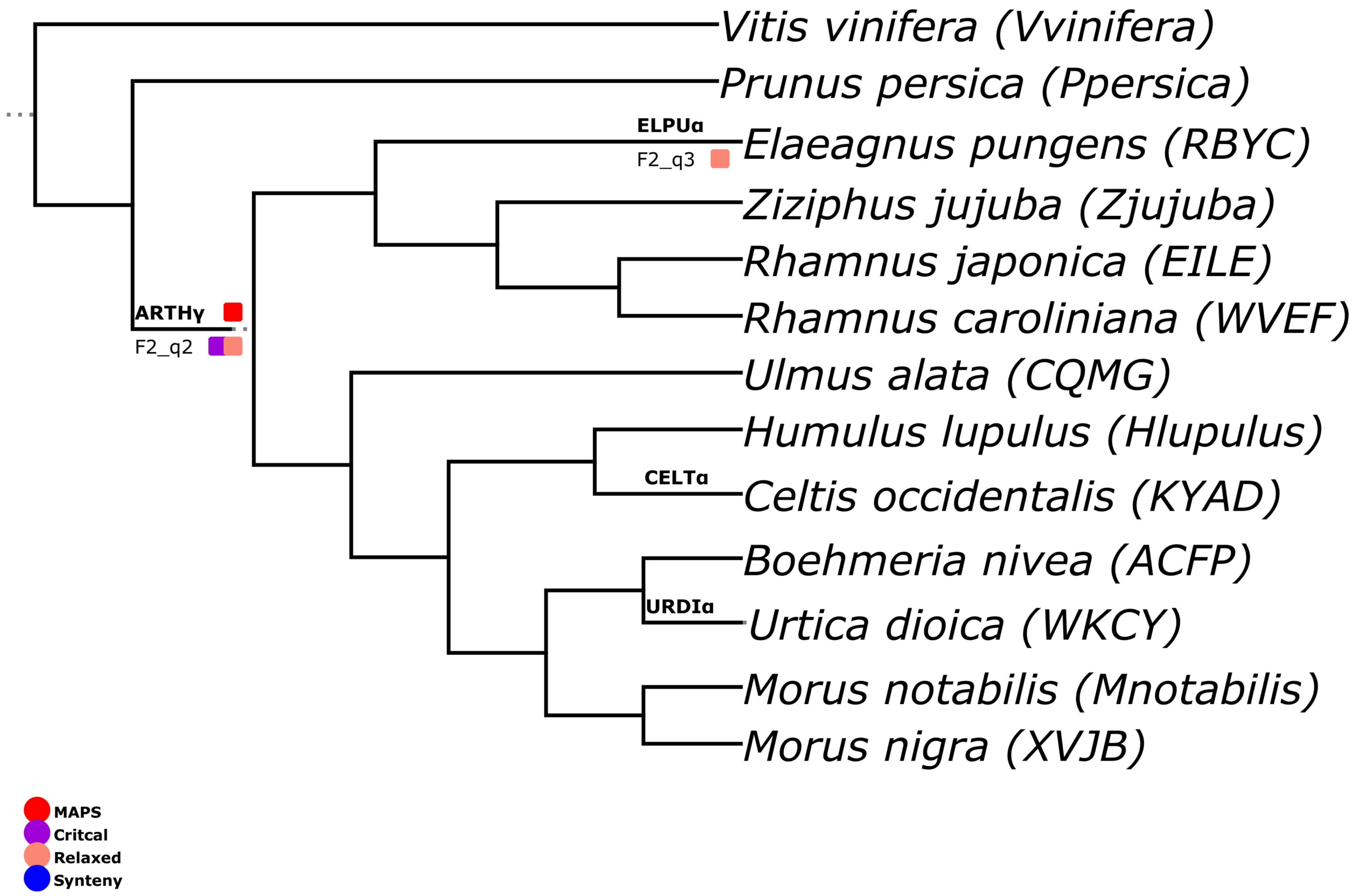


**Figure S15**

### G1 TimeTree

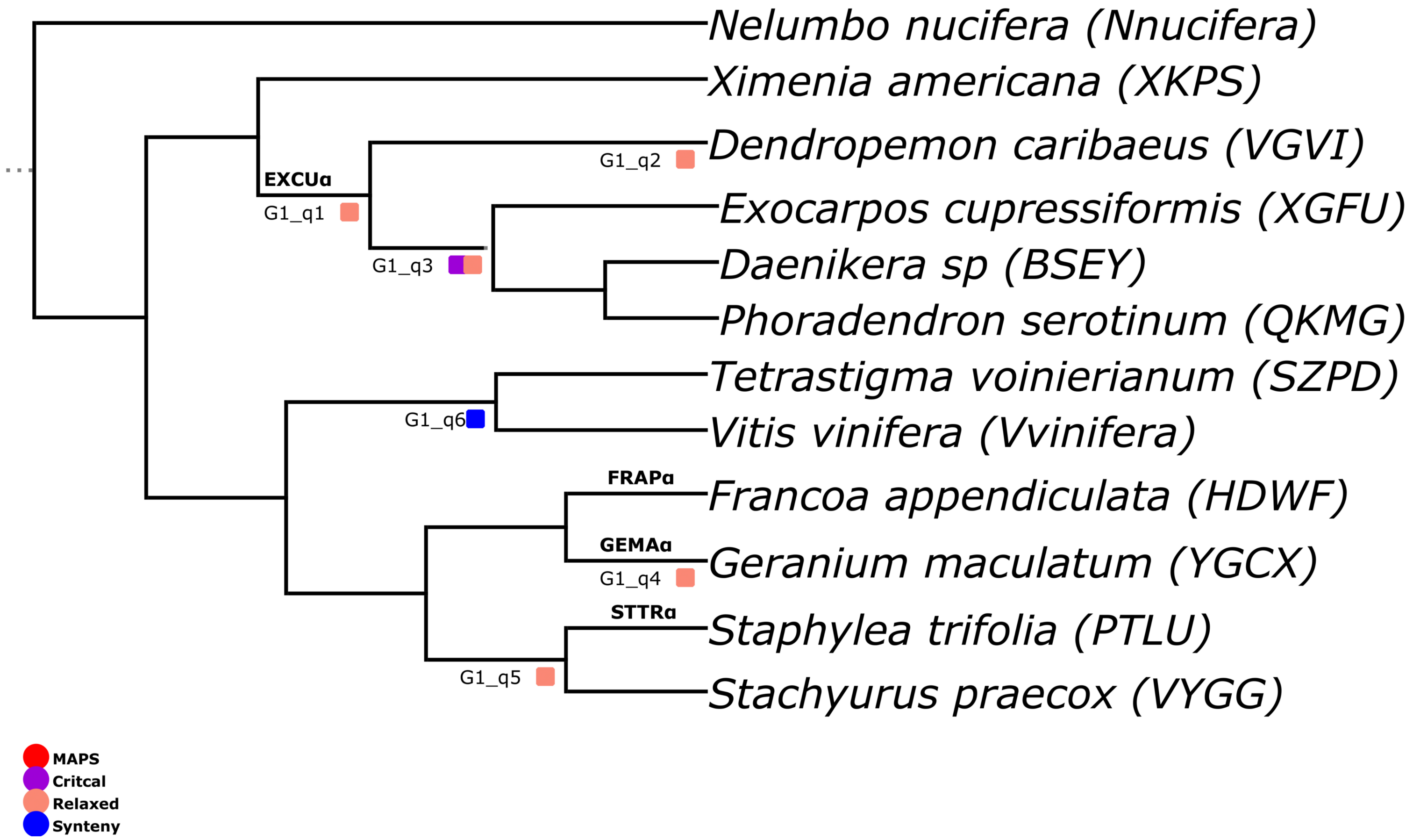


### Figure S16

### G2 TimeTree

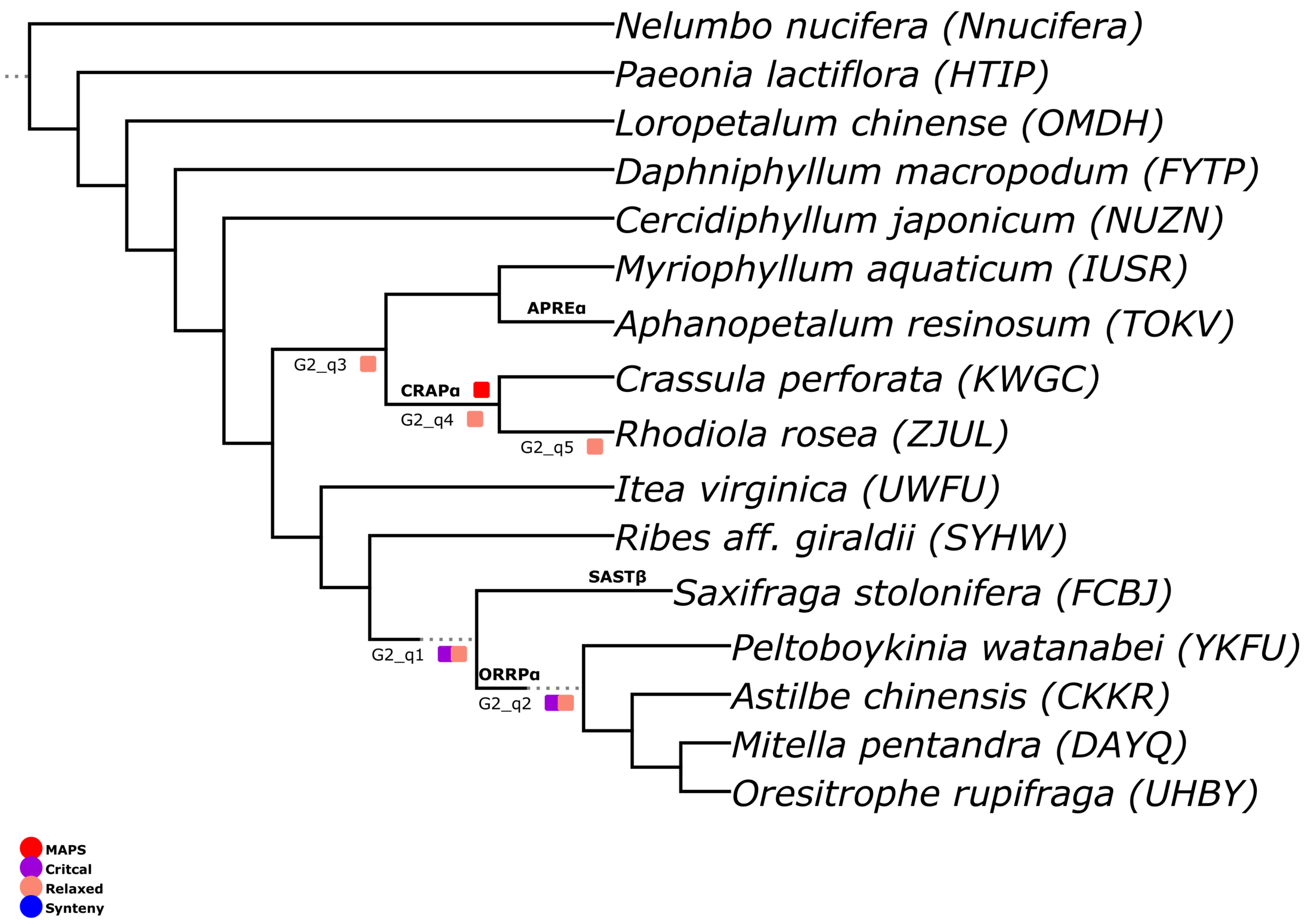


### Figure S17

### H1 TimeTree

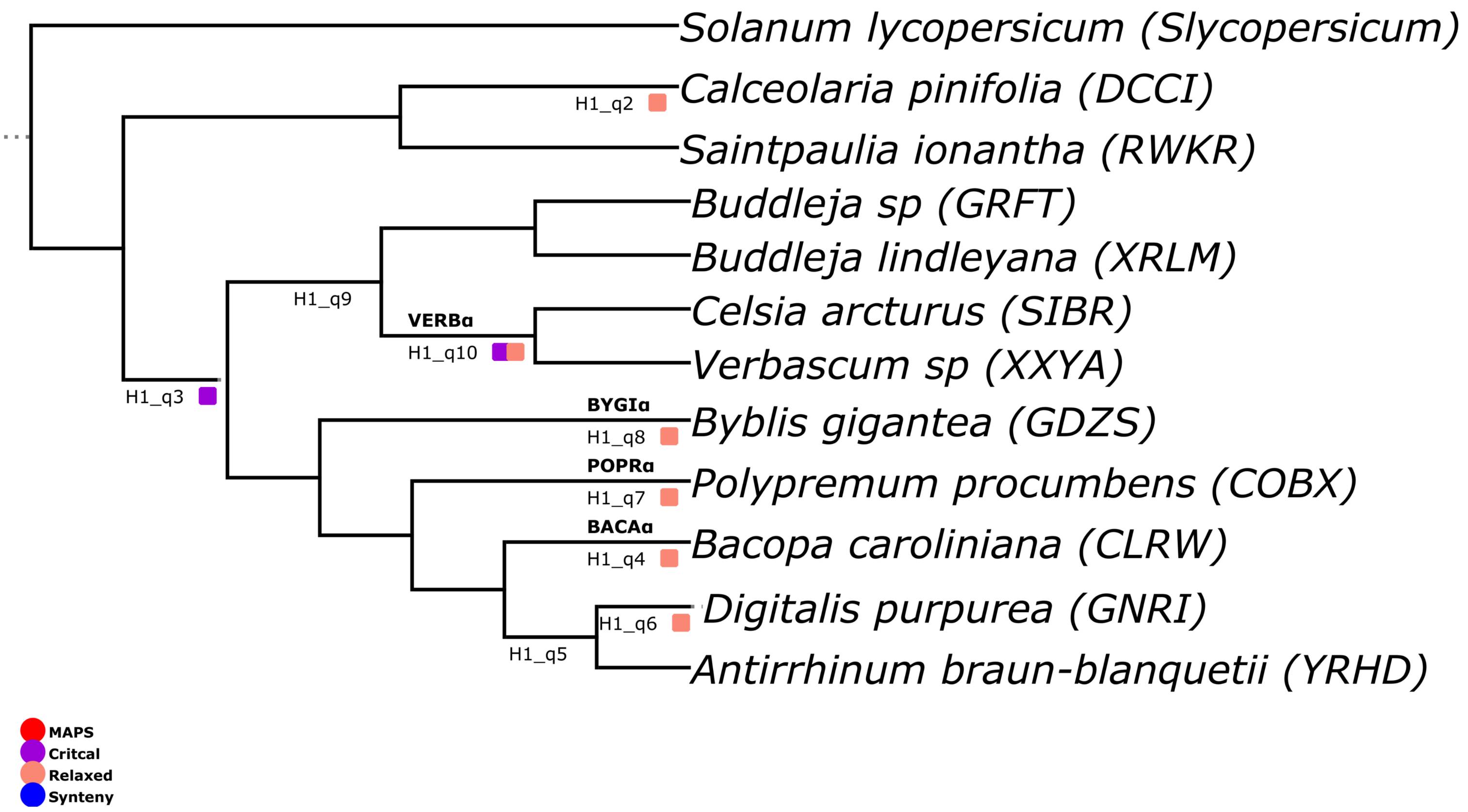


### Figure S18

### H2 TimeTree

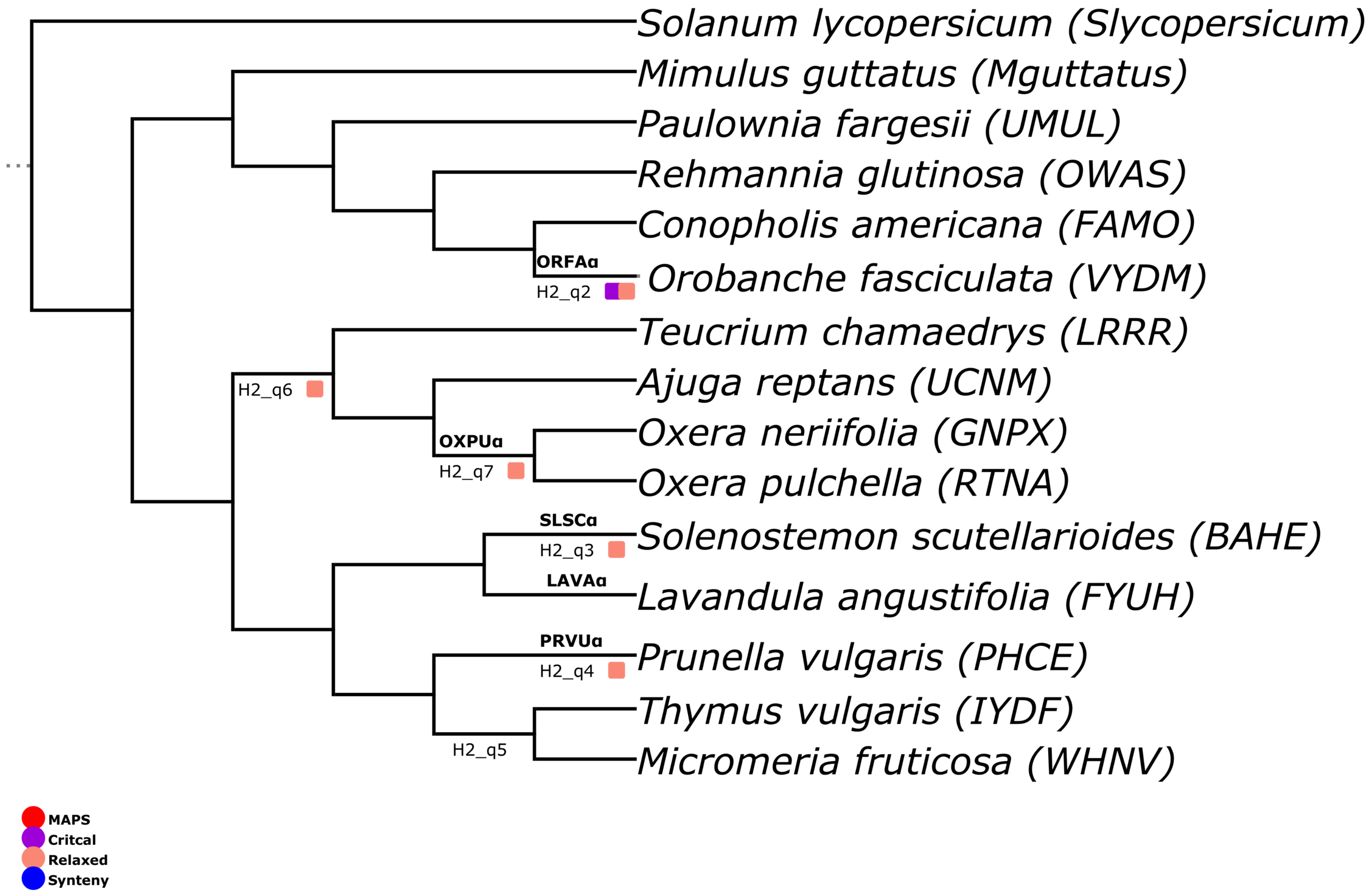


### Figure S19

### H3 TimeTree

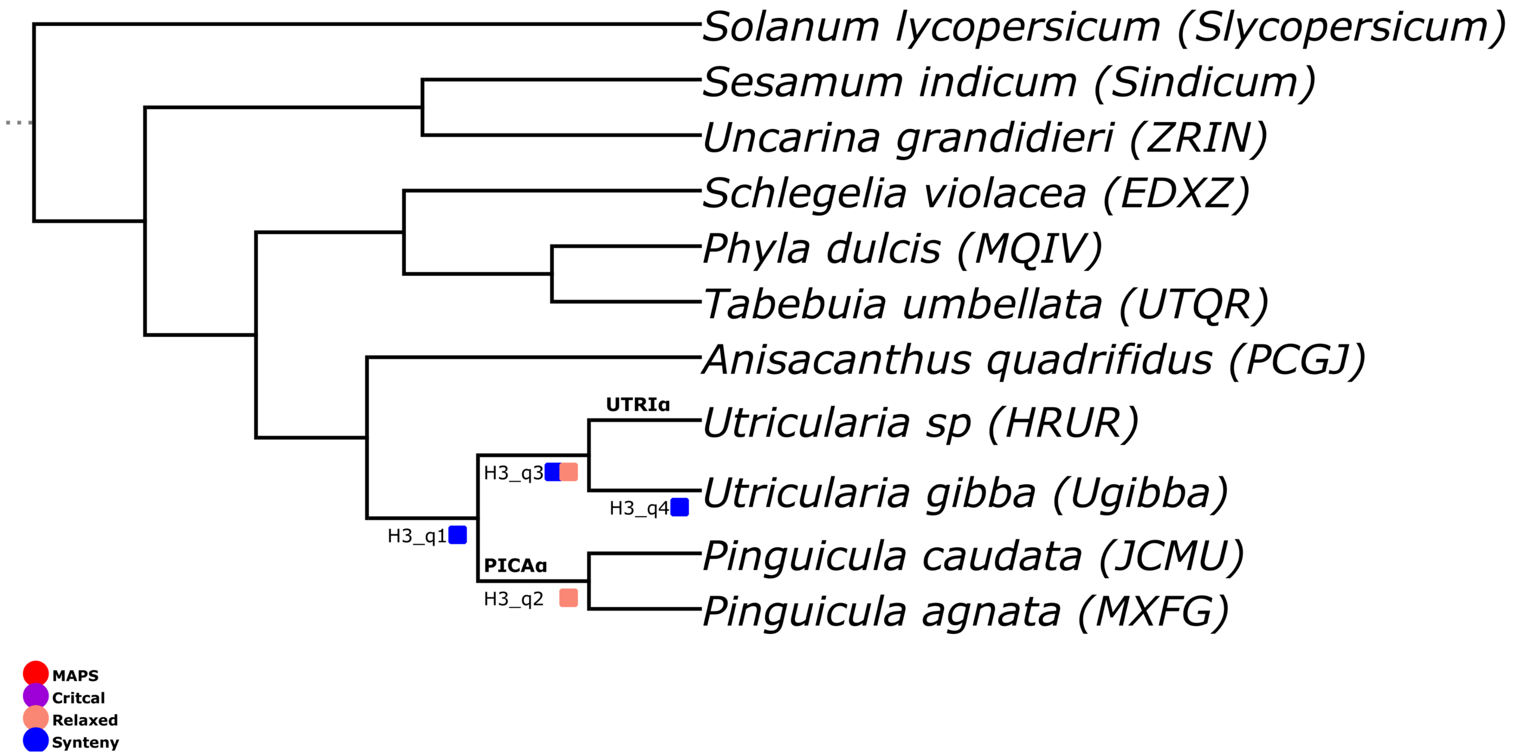


### Figure S20

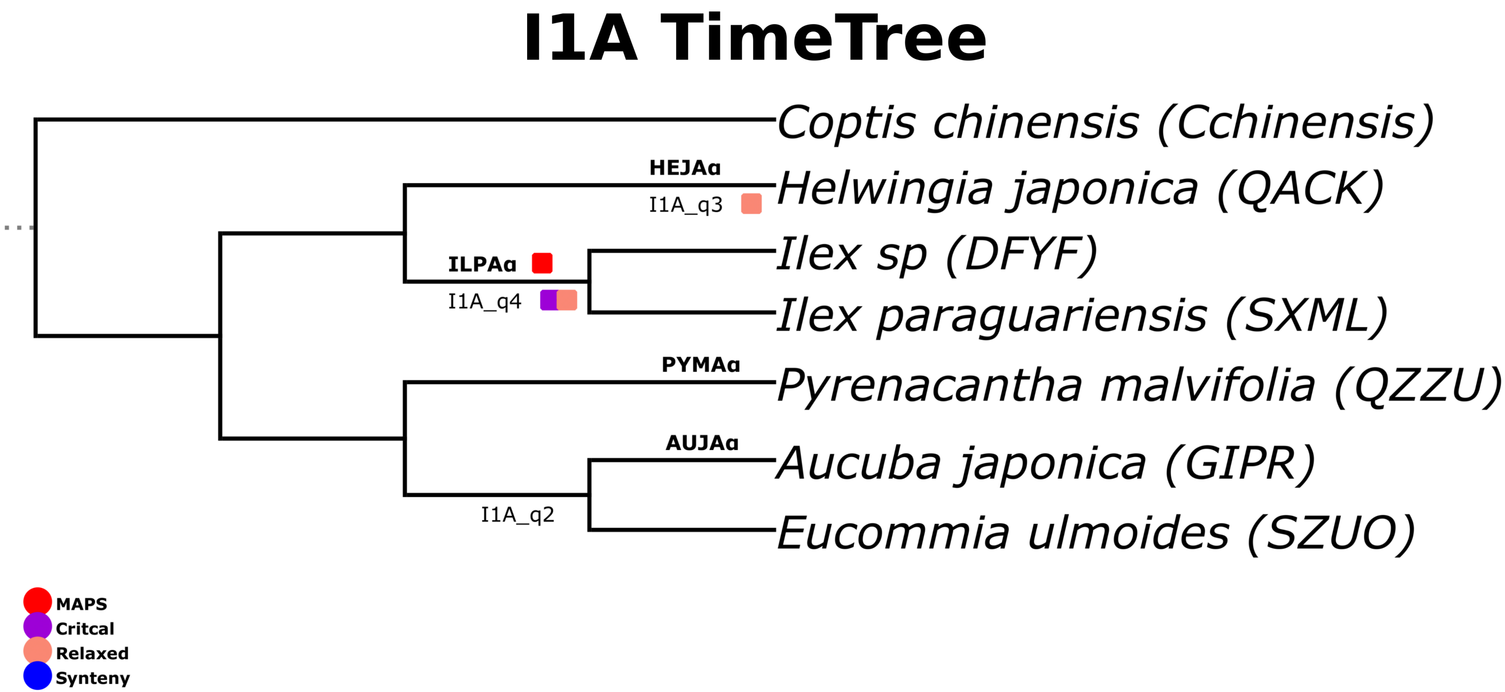

I1A TimeTree
Coptis chinensis (Cchinensis)
HEJAα
I1A_q3
Helwingia japonica (QACK)
ILPAα
I1A_q4
Ilex sp (DFYF)
Ilex paraguariensis (SXML)
PYMAα
Pyrenacantha malvifolia (QZZU)
AUJAα
Aucuba japonica (GIPR)
I1A_q2
Eucommia ulmoides (SZUO)
MAPS
Critcal
Relaxed
Synteny


### Figure S21

### I1B TimeTree

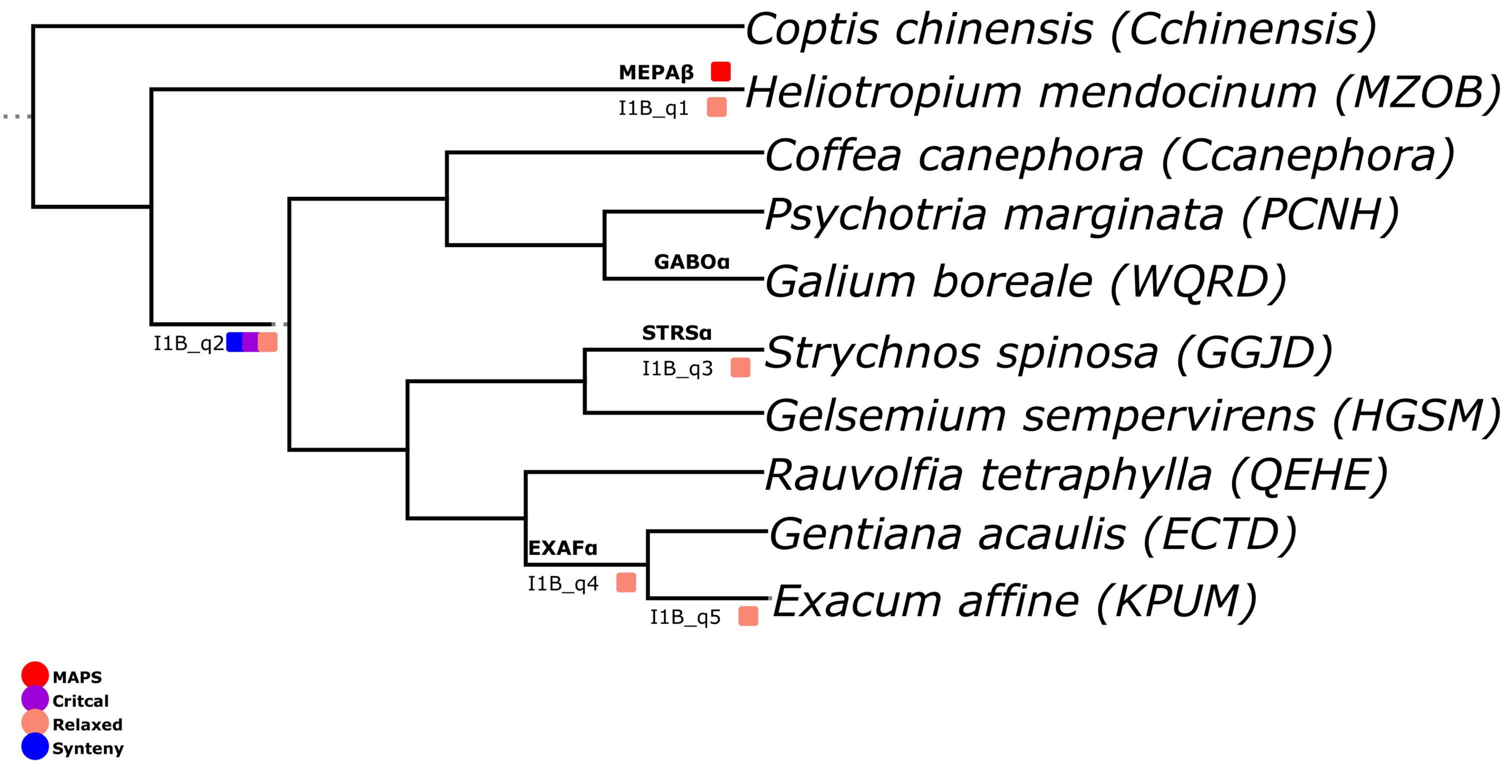


**Figure S22**

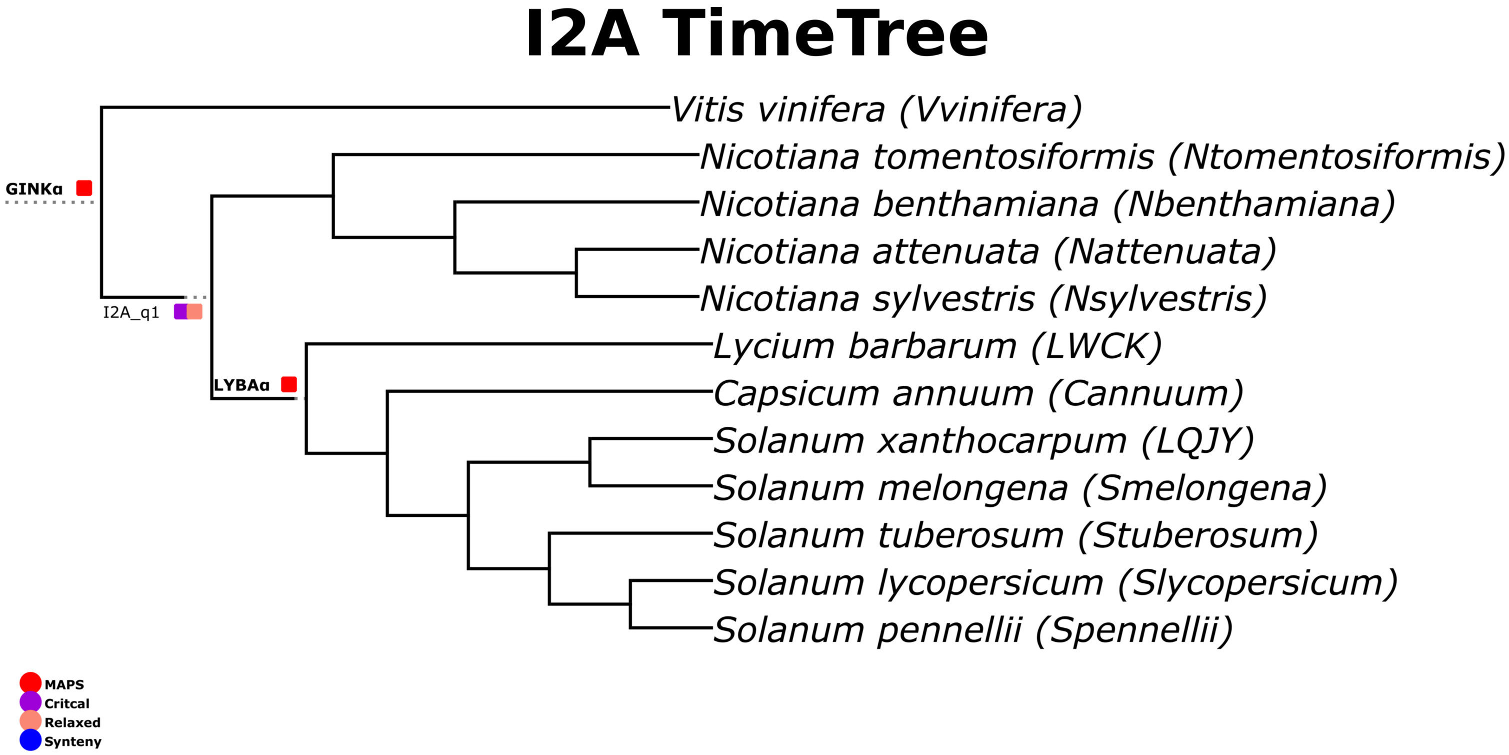

I2A TimeTree
Vitis vinifera (Vvinifera)
Nicotiana tomentosiformis (Ntomentosiformis)
Nicotiana benthamiana (Nbenthamiana)
Nicotiana attenuata (Nattenuata)
Nicotiana sylvestris (Nsylvestris)
Lycium barbarum (LWCK)
Capsicum annuum (Cannuum)
Solanum xanthocarpum (LQJY)
Solanum melongena (Smelongena)
Solanum tuberosum (Stuberosum)
Solanum lycopersicum (Slycopersicum)
Solanum pennellii (Spennellii)
GINKα
I2A_q1
LYBAα
MAPS
Critcal
Relaxed
Synteny


### Figure S23

### I2B TimeTree

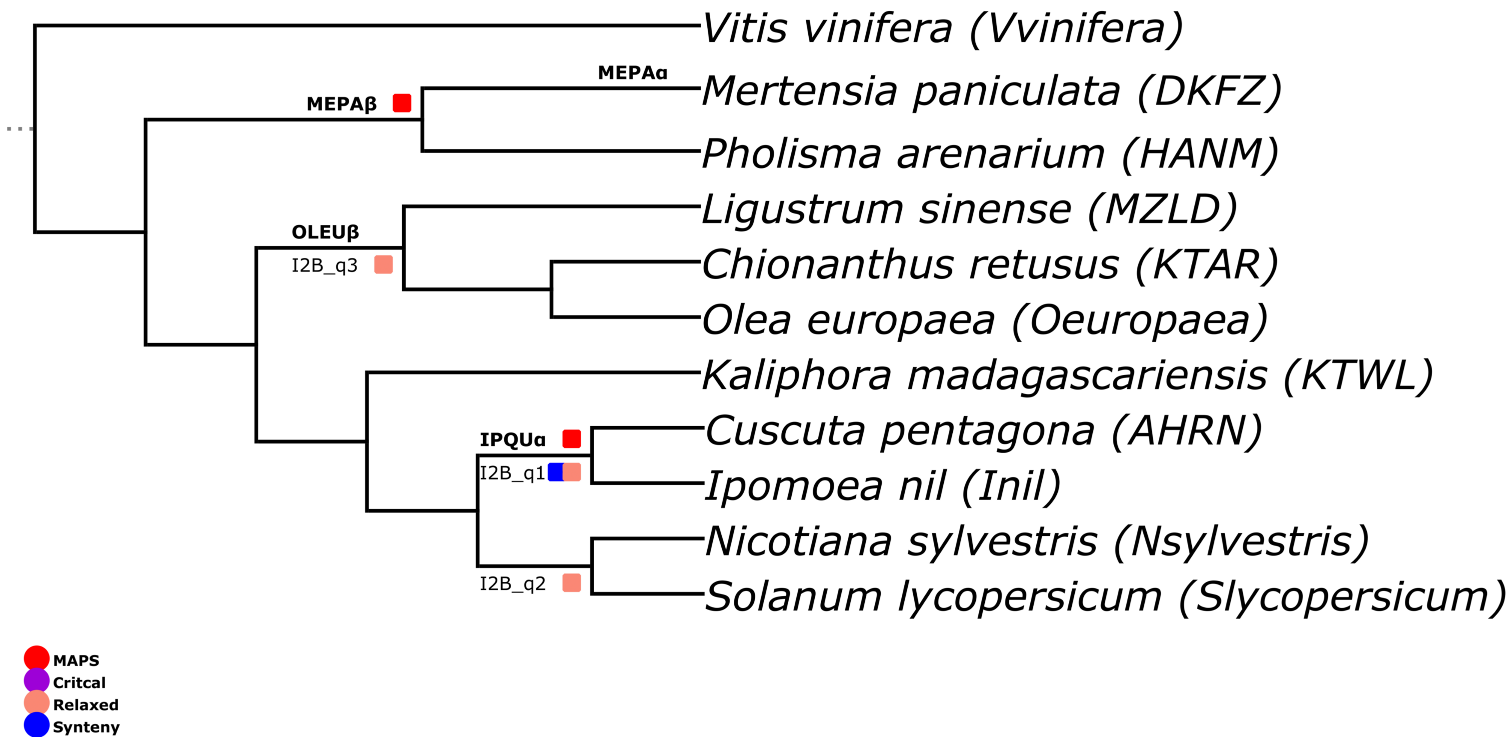


### Figure S24

### J1 TimeTree

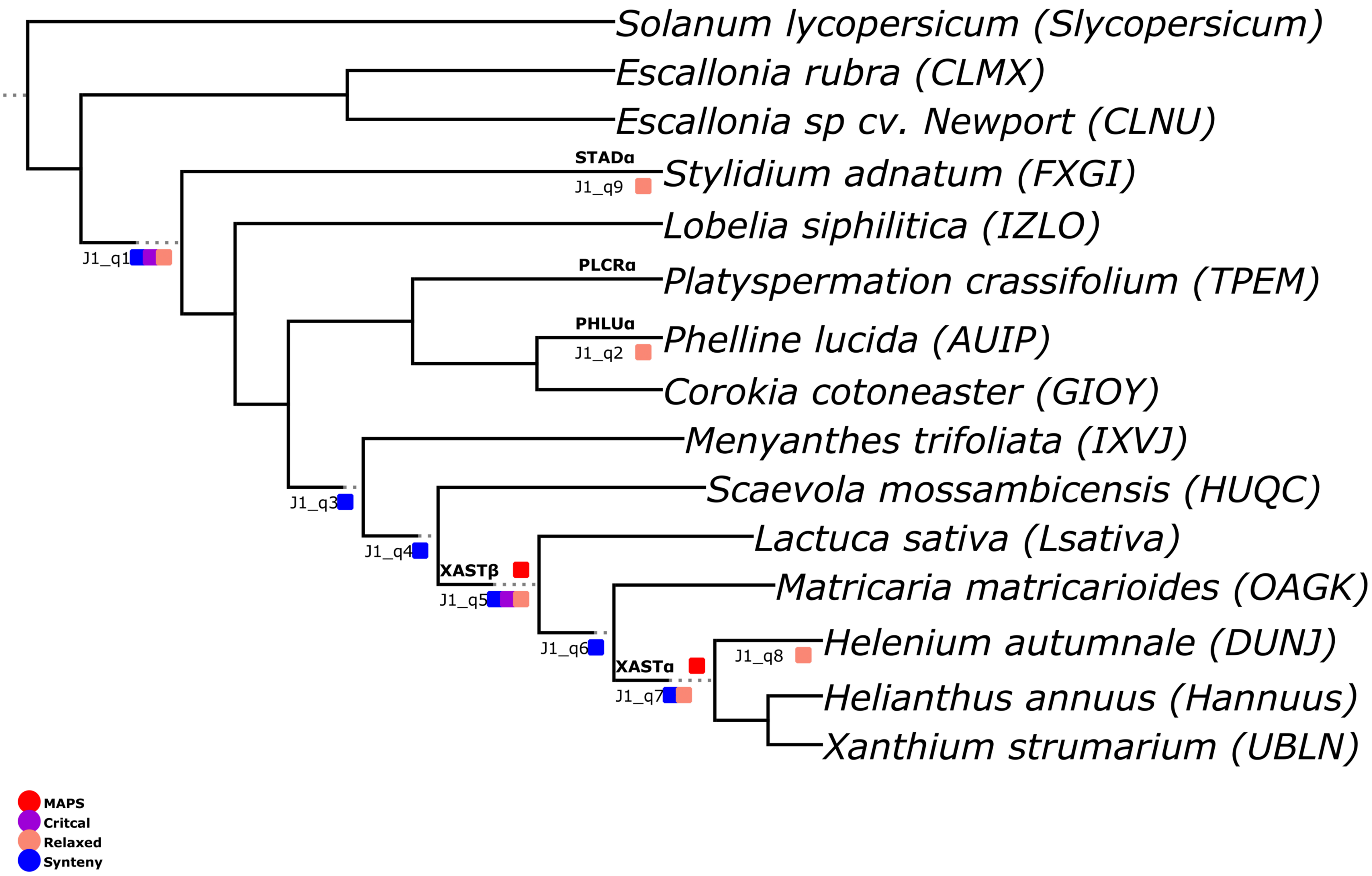


### Figure S25

### J2 TimeTree

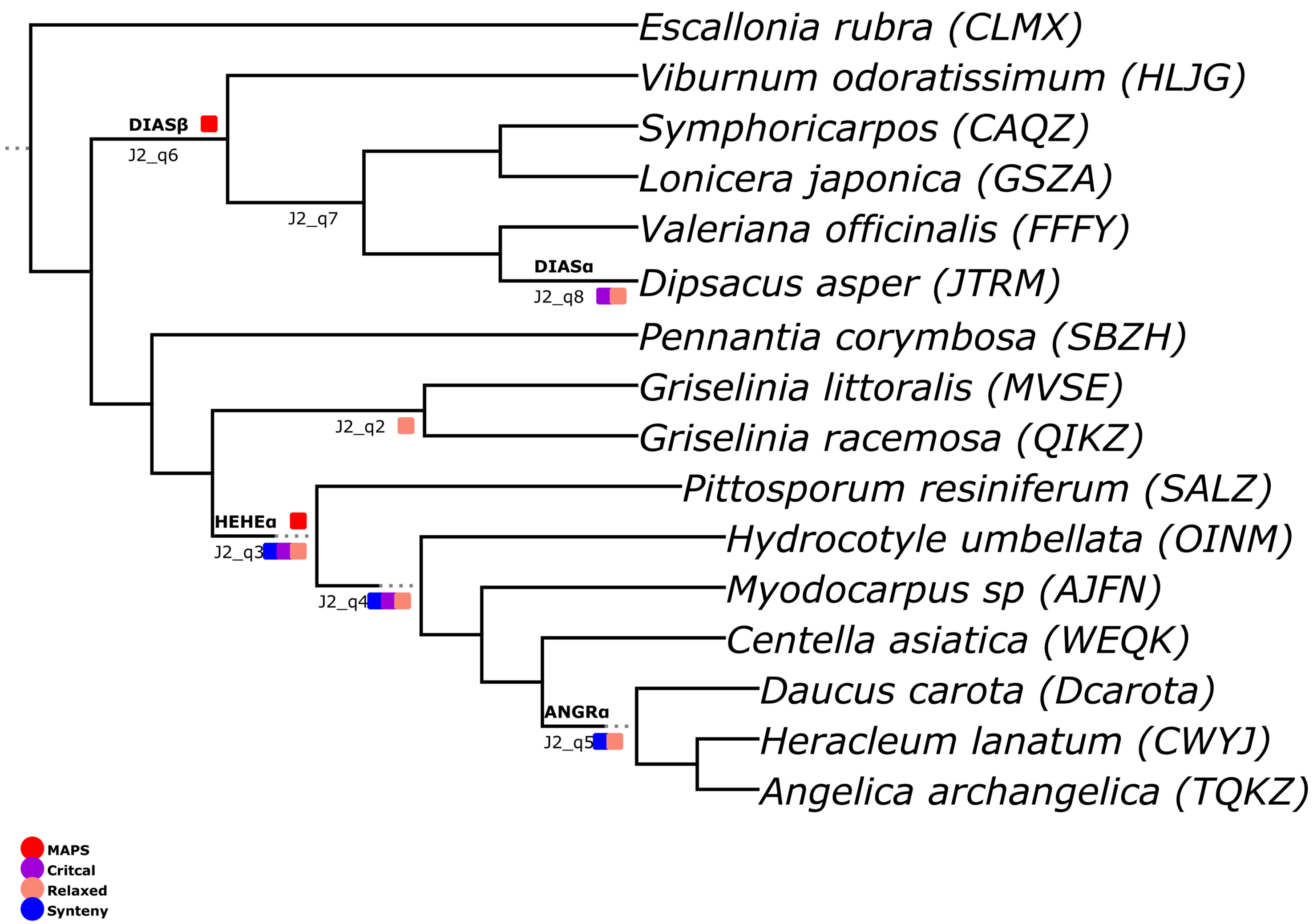


### Figure S26

### K1 TimeTree

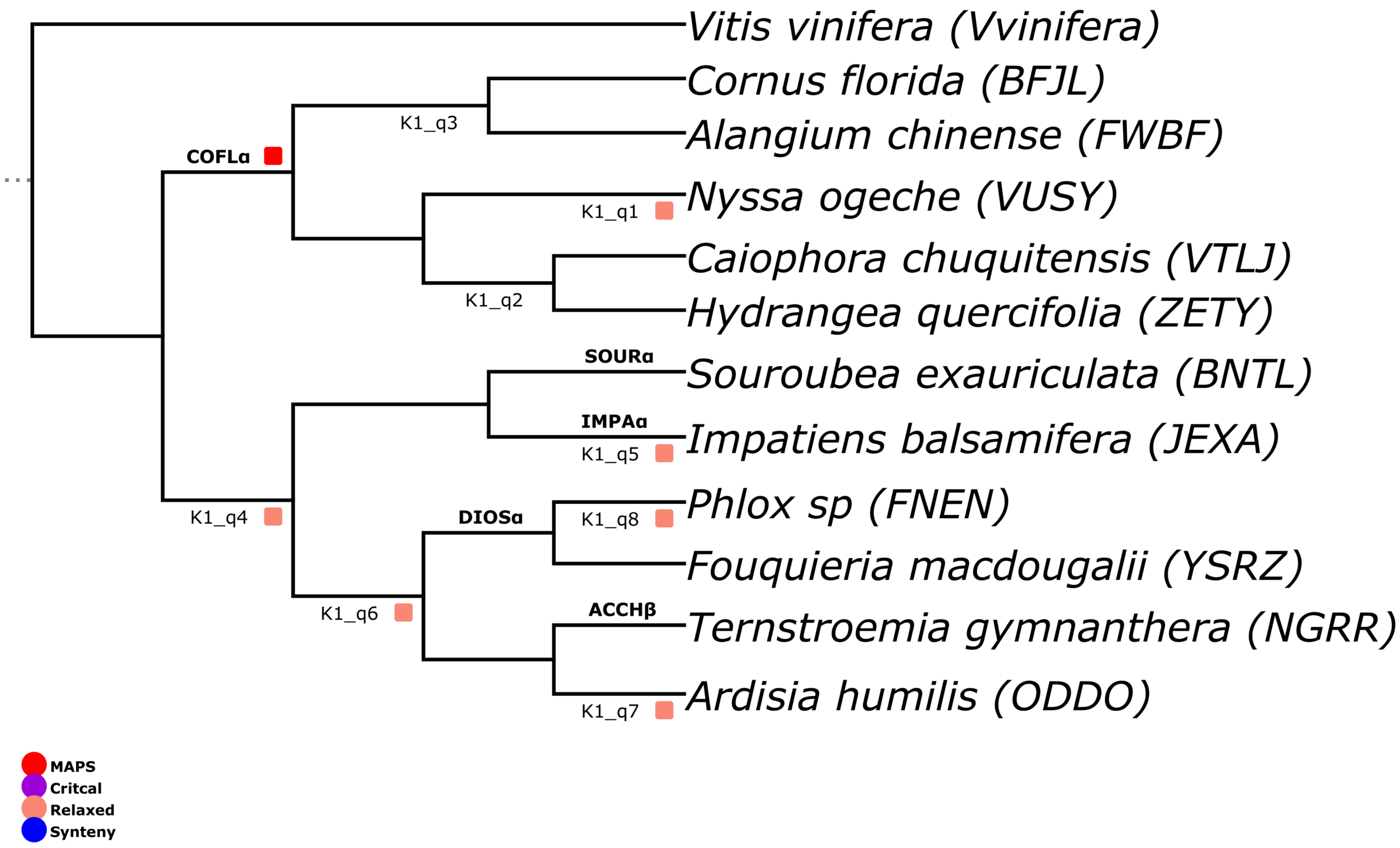


### Figure S27

### K2 TimeTree

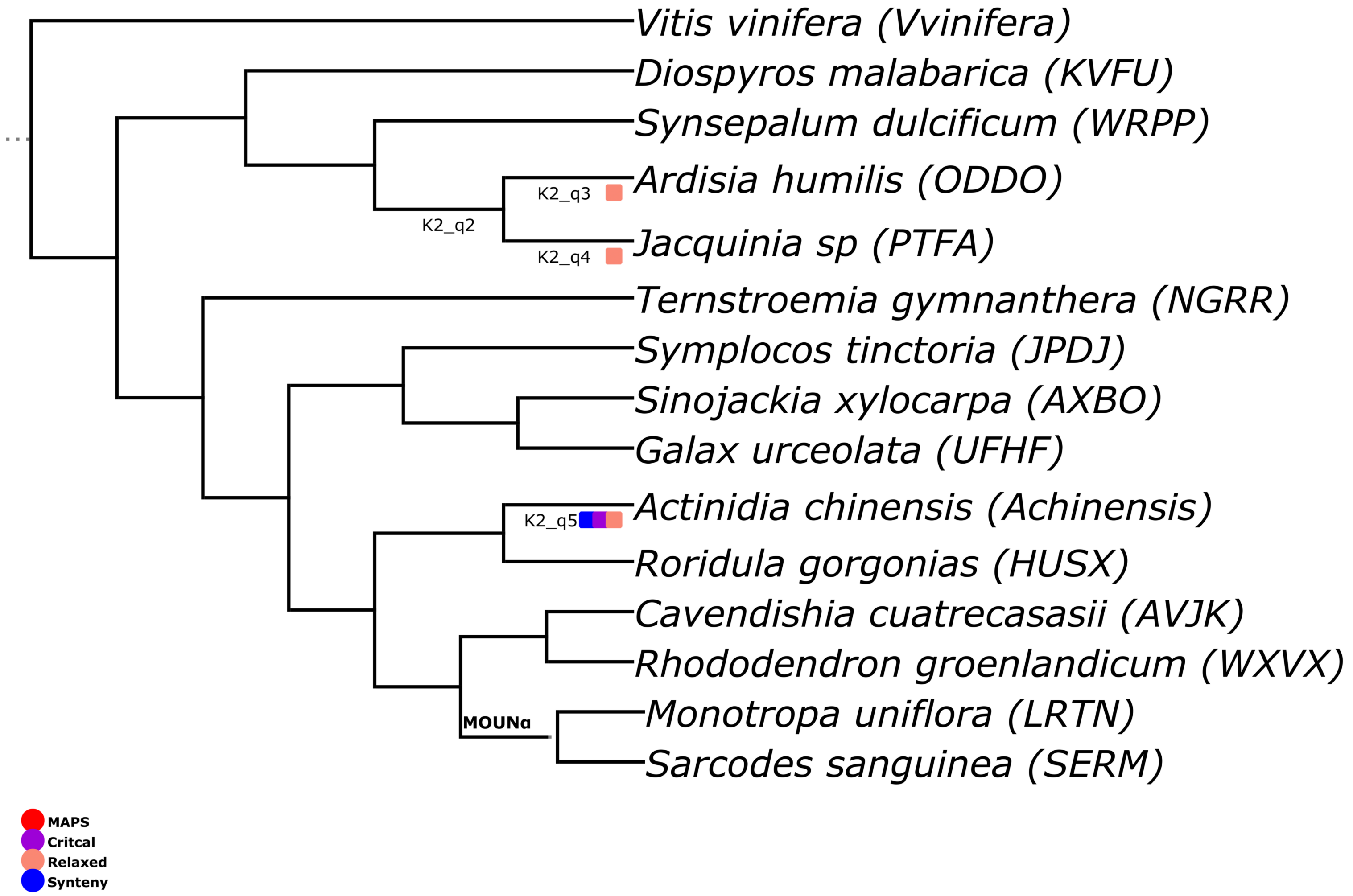


### Figure S28

### L1 TimeTree

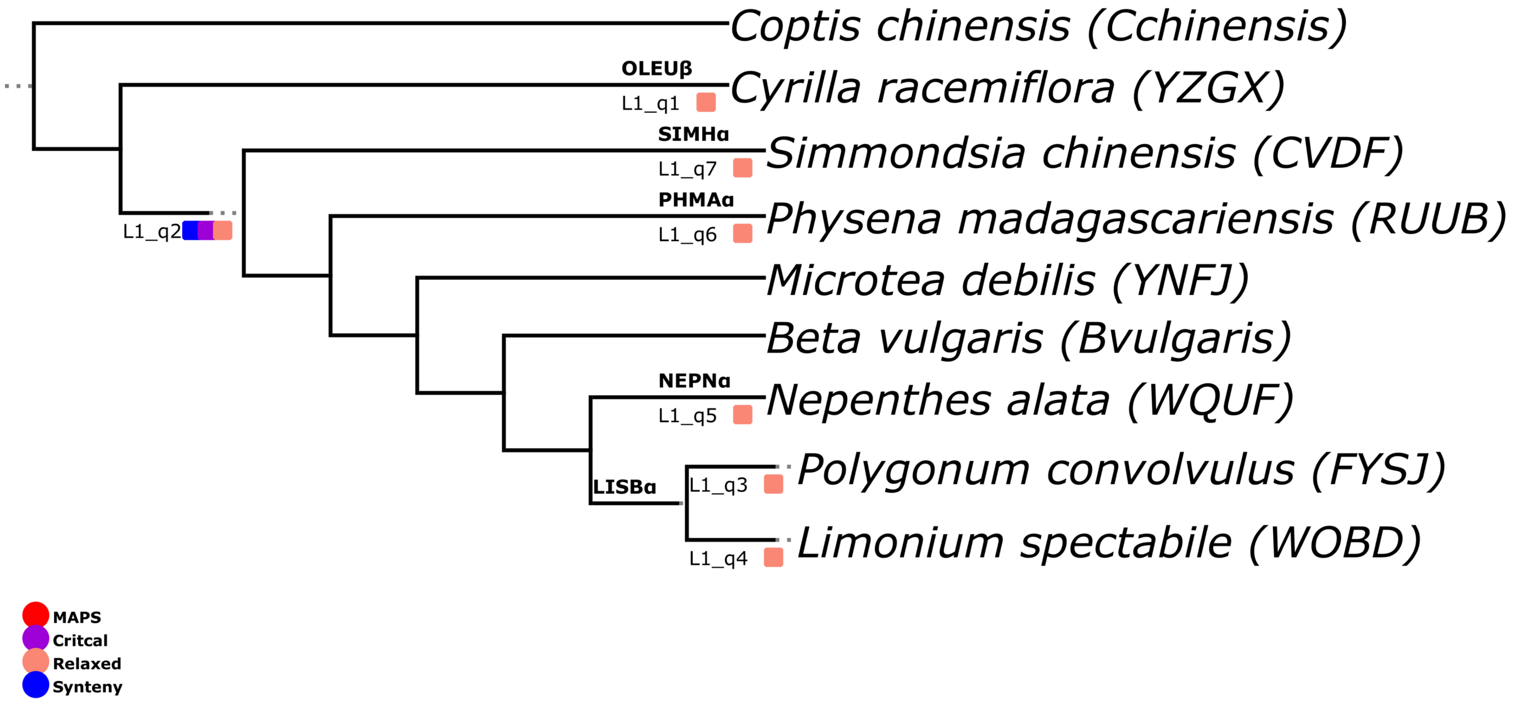


### Figure S29

### L2 TimeTree

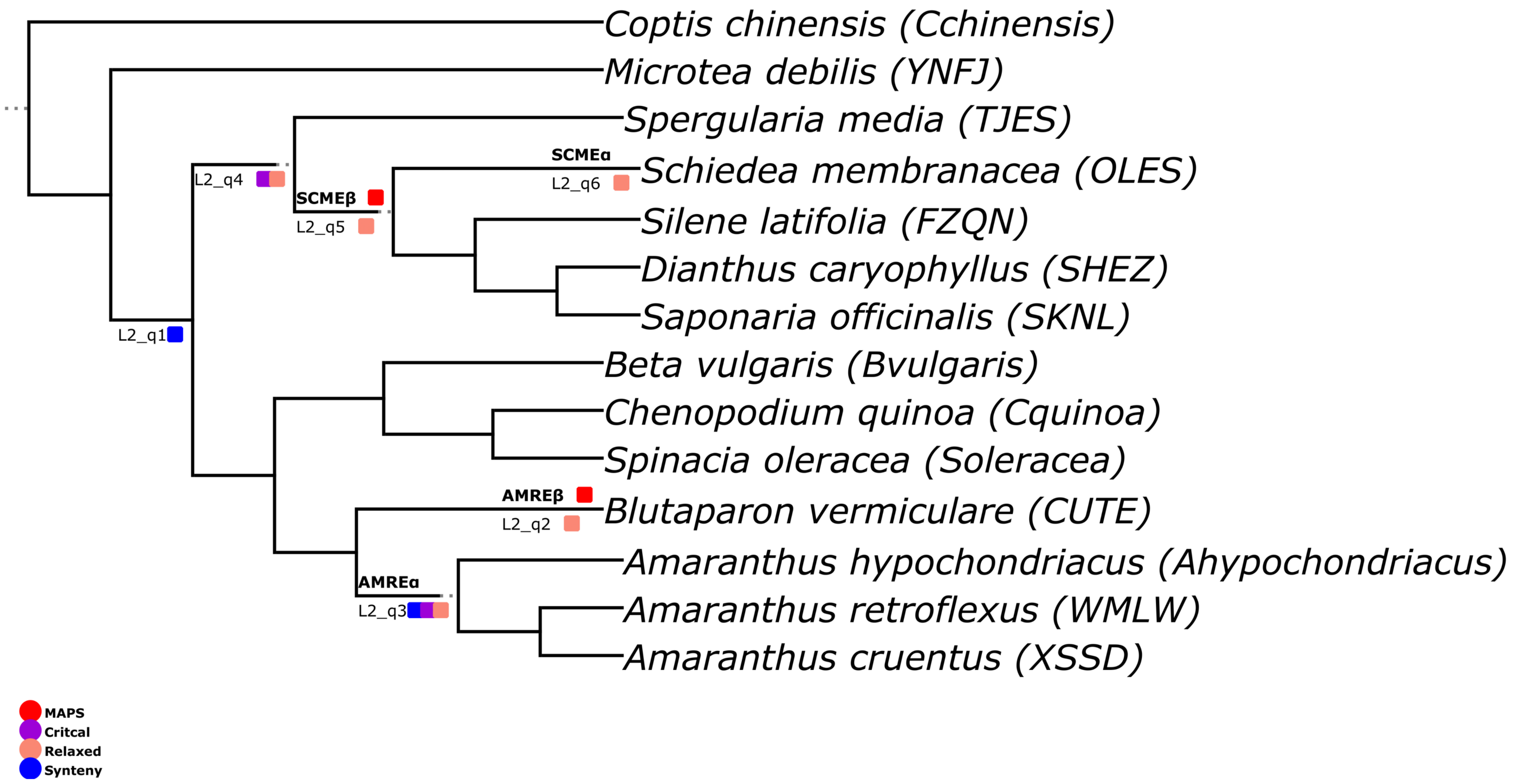


### Figure S30

### L3 TimeTree

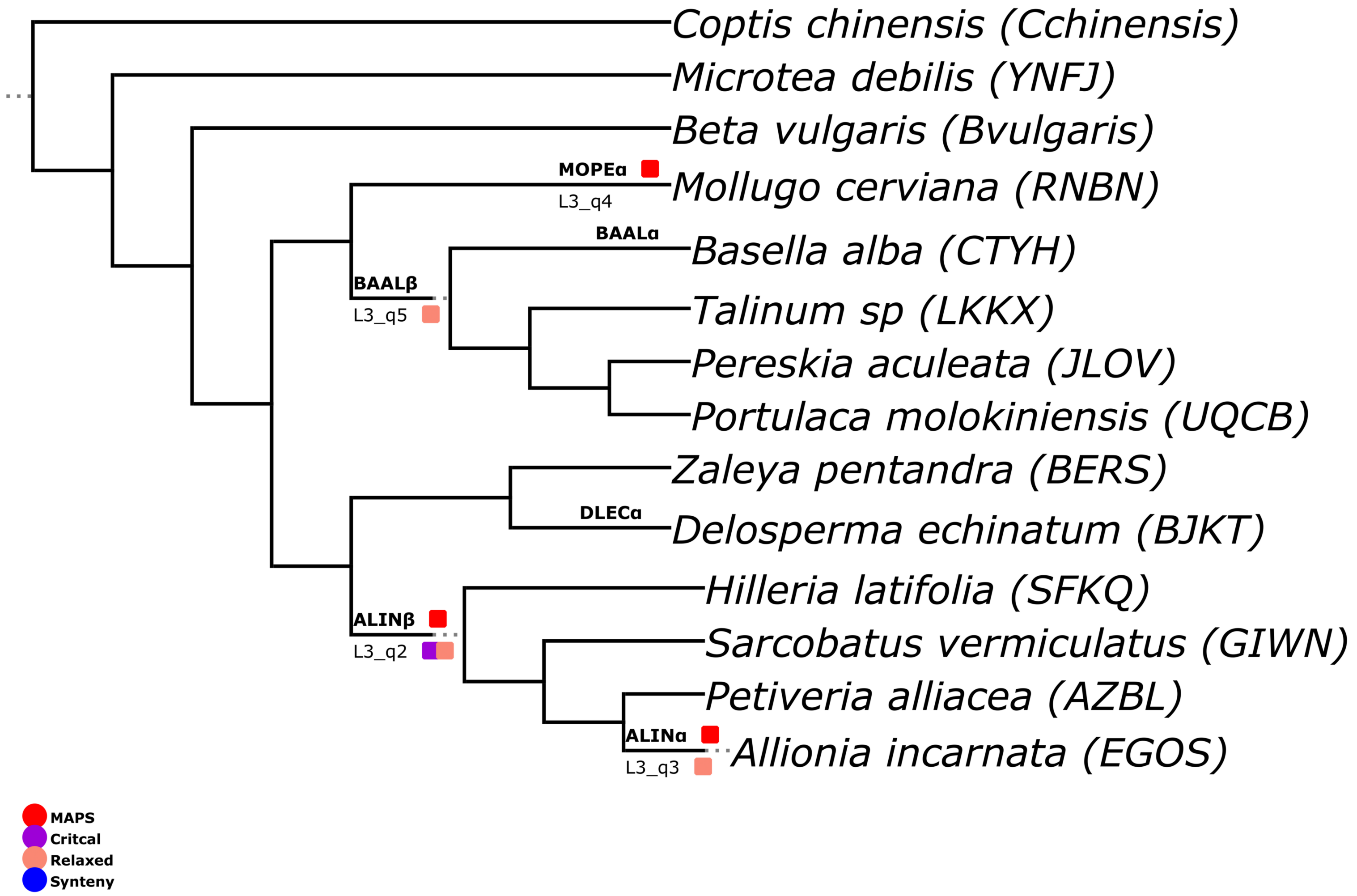


### Figure S31

### M1 TimeTree

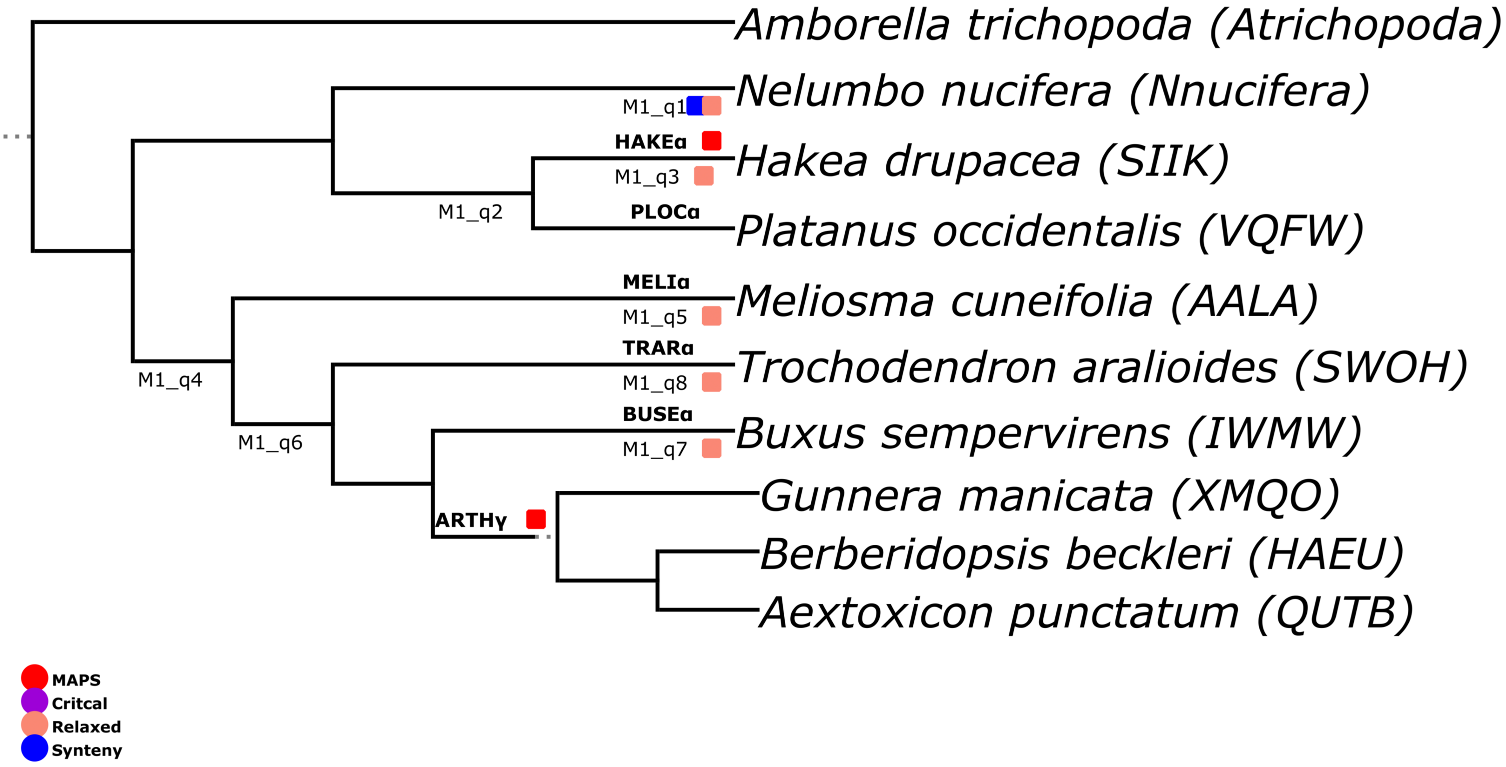


### Figure S32

### M2 TimeTree

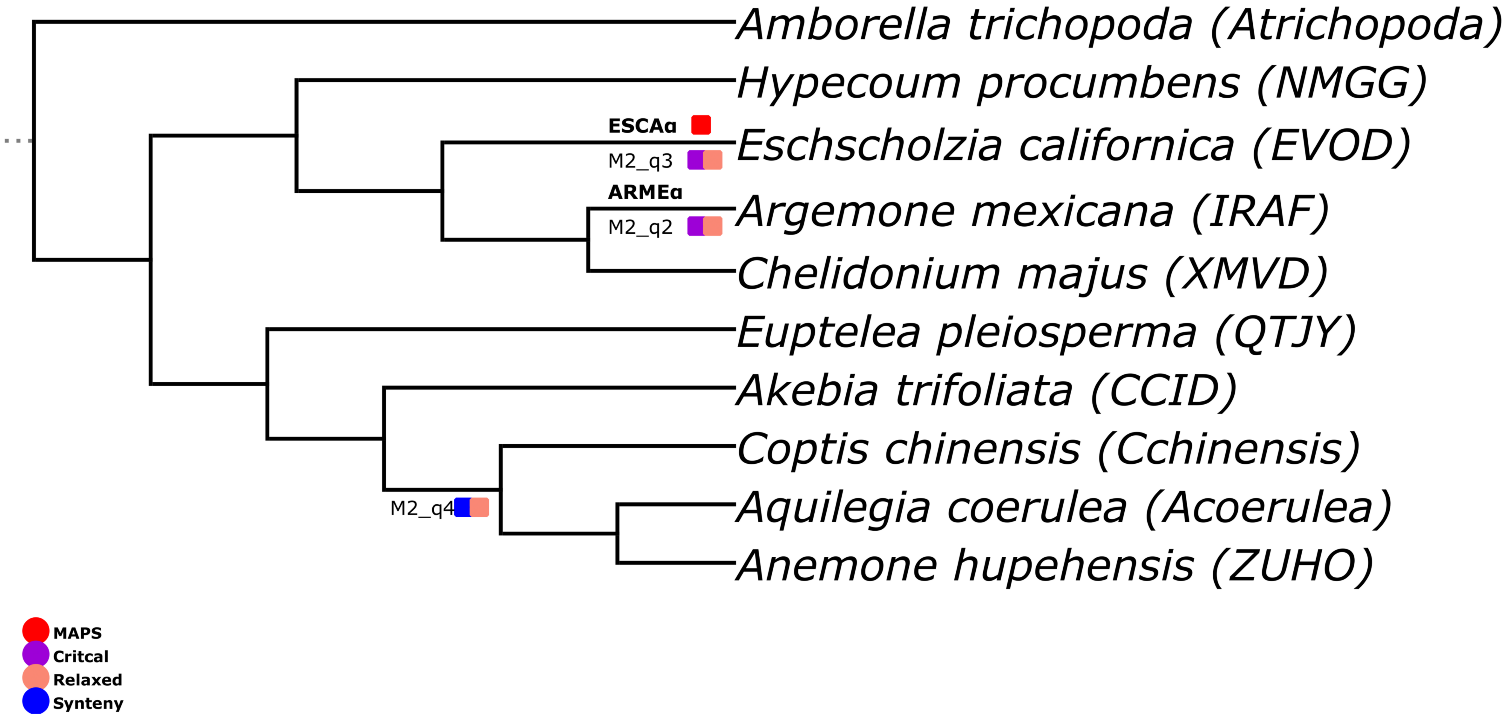


### Figure S33

### N1 TimeTree

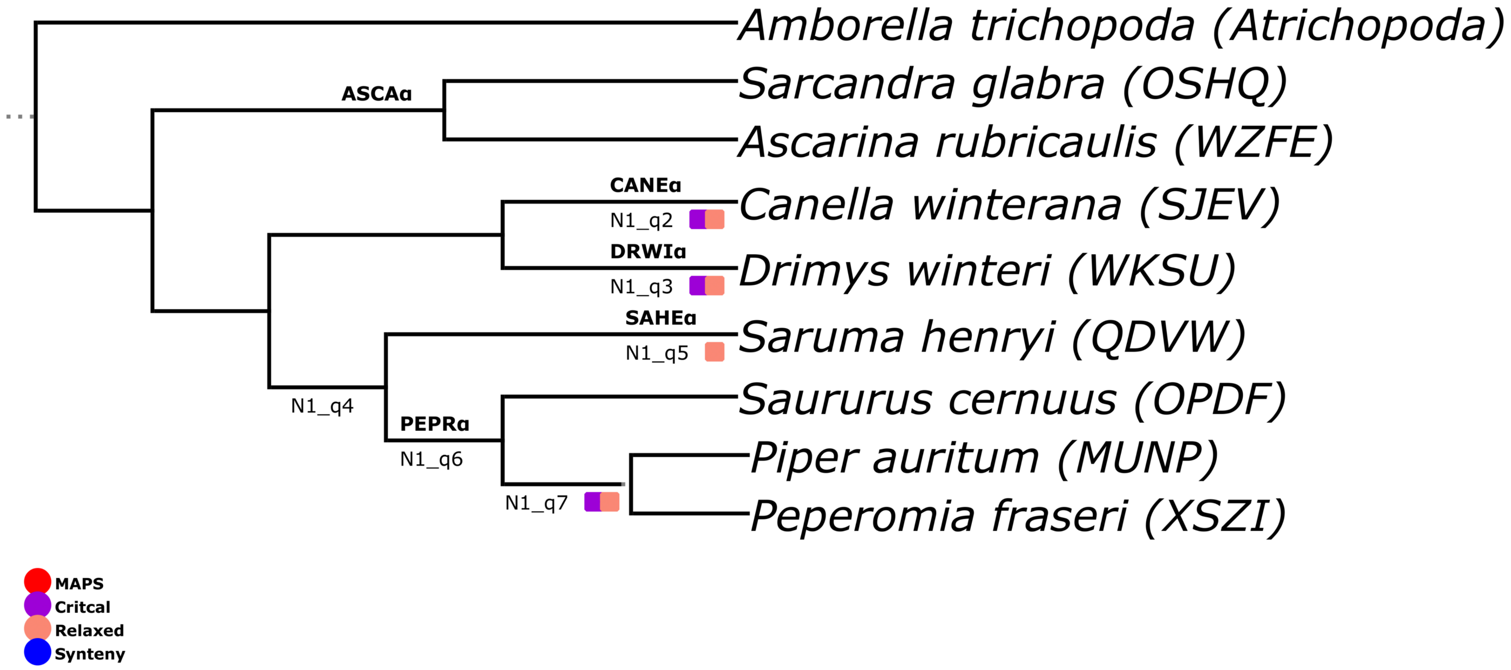


**Figure S34**

### N2 TimeTree

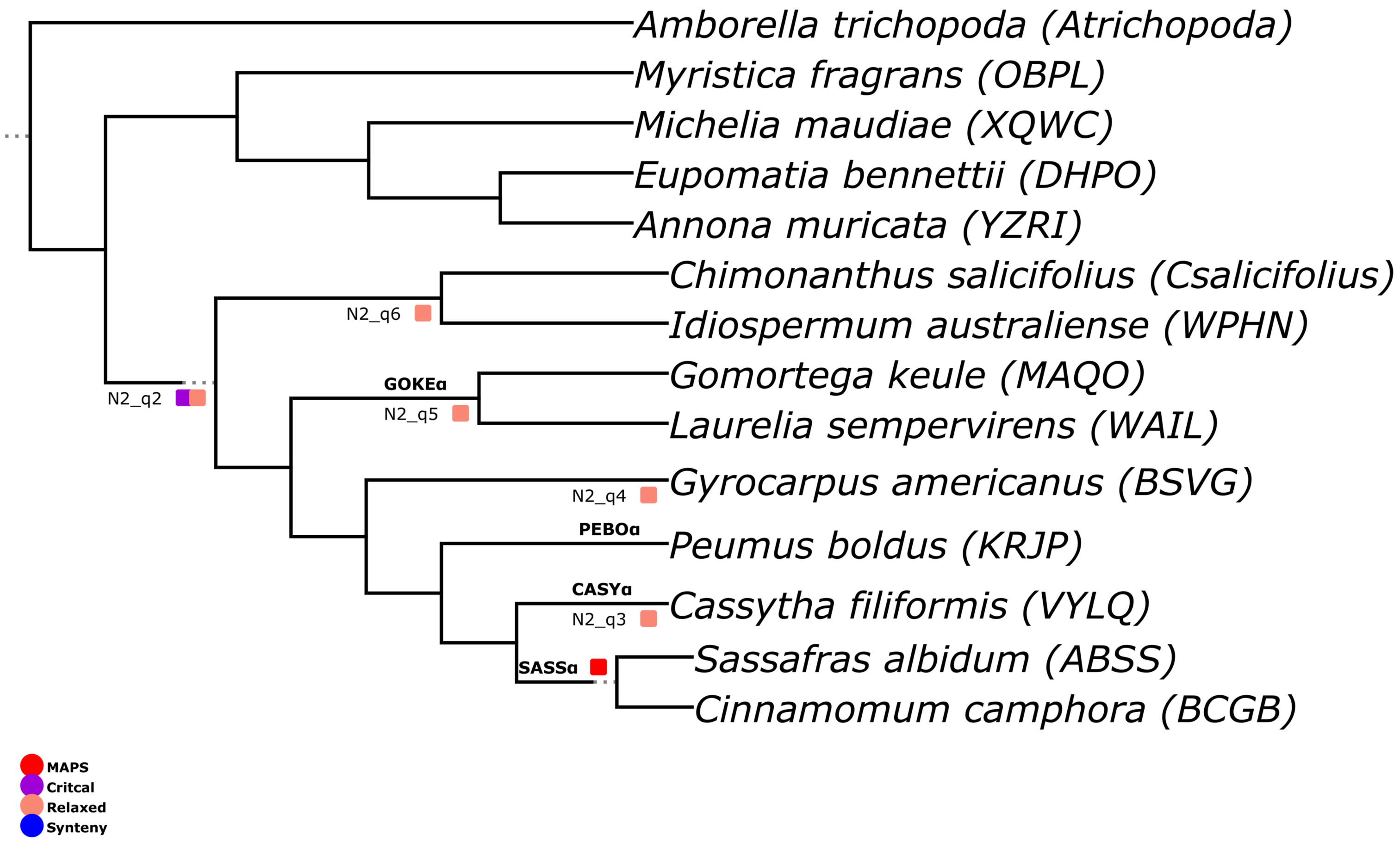


### Figure S35

### O1 TimeTree

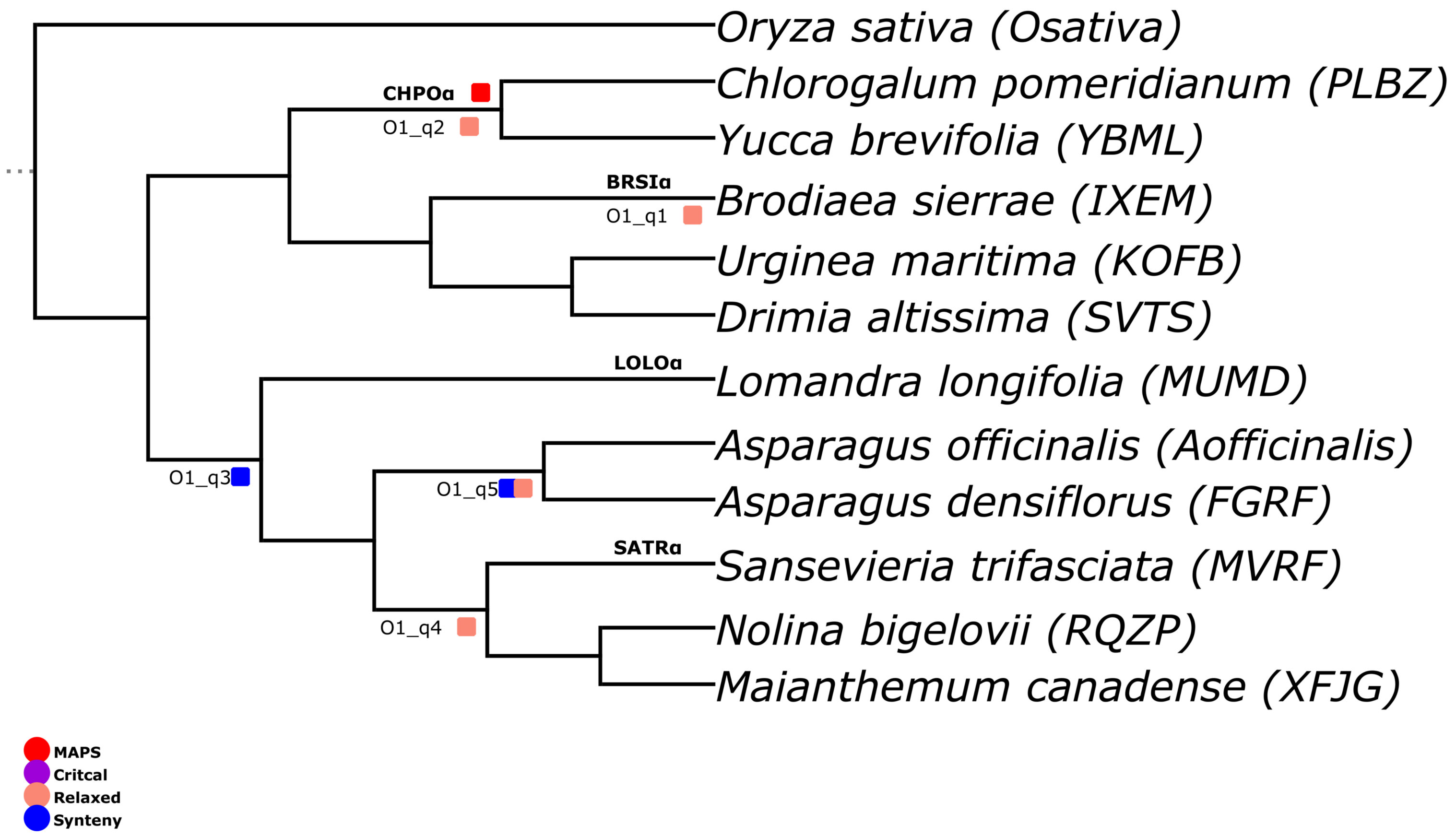


**Figure S36**

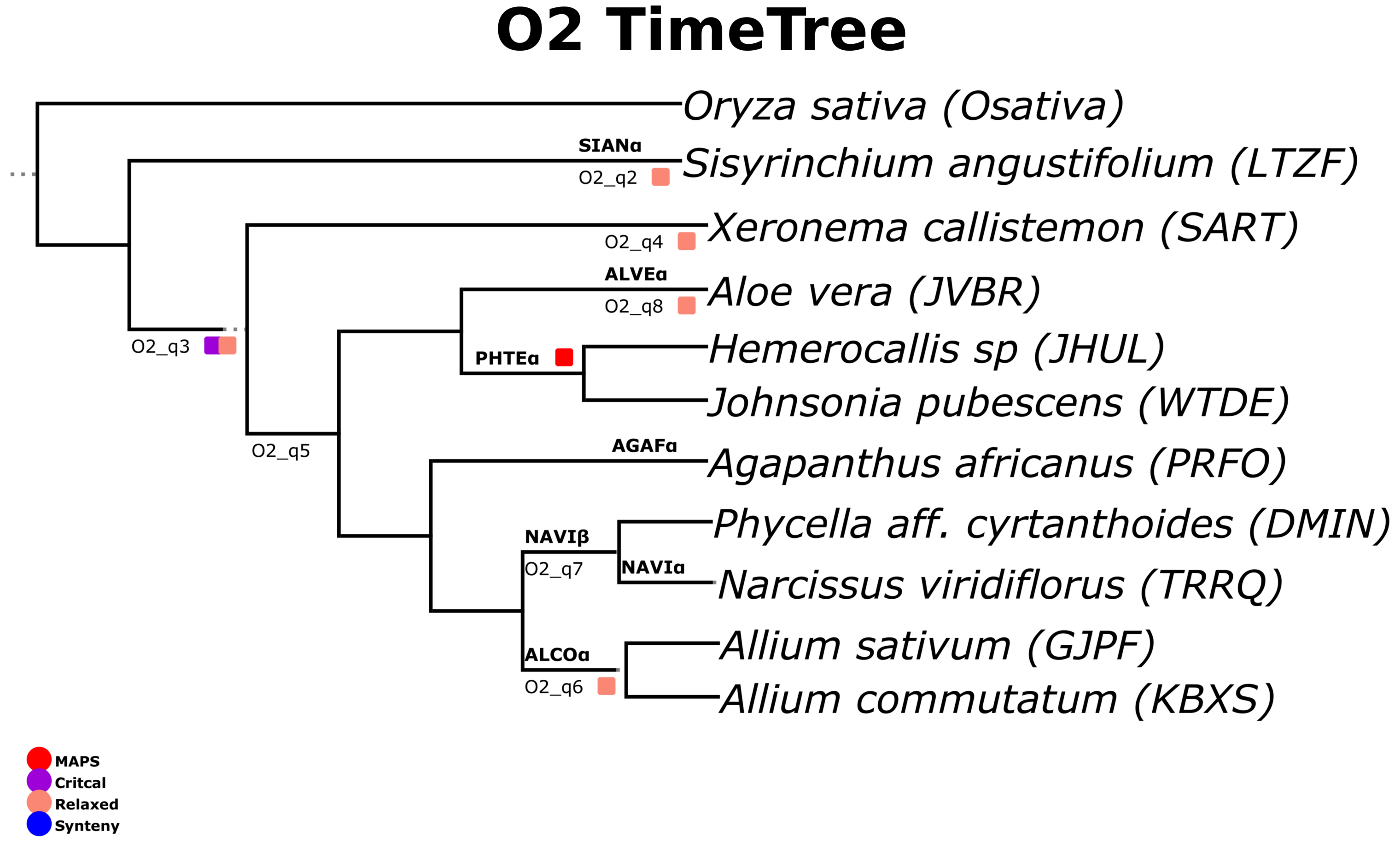
O2 TimeTree
Oryza sativa (Osativa)
SIANα
O2_q2
Sisyrinchium angustifolium (LTZF)
O2_q4
Xeronema callistemon (SART)
ALVEα
O2_q8
Aloe vera (JVBR)
O2_q3
PHTEα
Hemerocallis sp (JHUL)
Johnsonia pubescens (WTDE)
O2_q5
AGAFα
Agapanthus africanus (PRFO)
NAVIβ
O2_q7
NAVIα
Phycella aff. cyrtanthoides (DMIN)
Narcissus viridiflorus (TRRQ)
ALCOα
O2_q6
Allium sativum (GJPF)
Allium commutatum (KBXS)
MAPS
Critcal
Relaxed
Synteny

### Figure S37

### P1 TimeTree

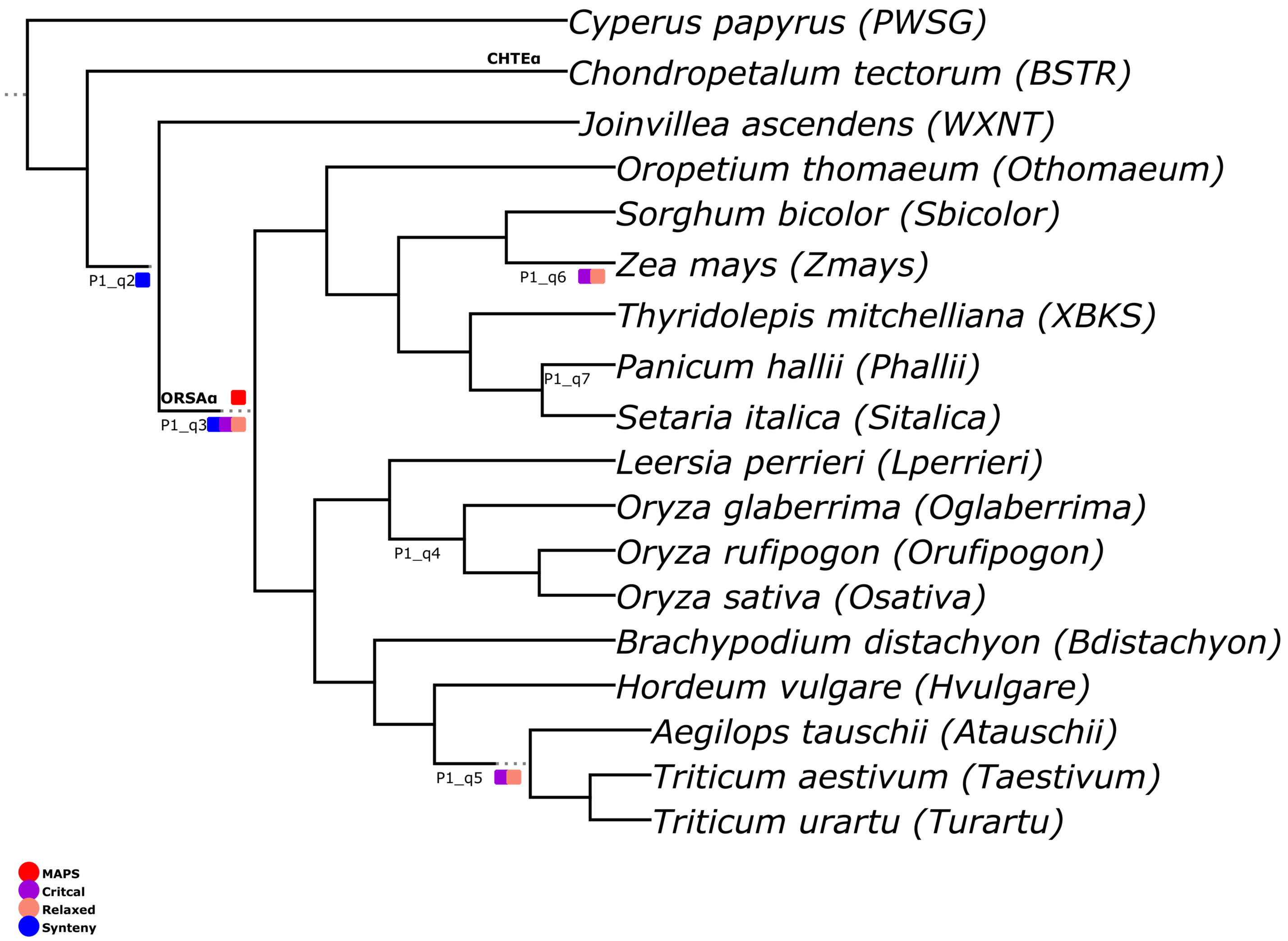


### Figure S38

### P2 TimeTree

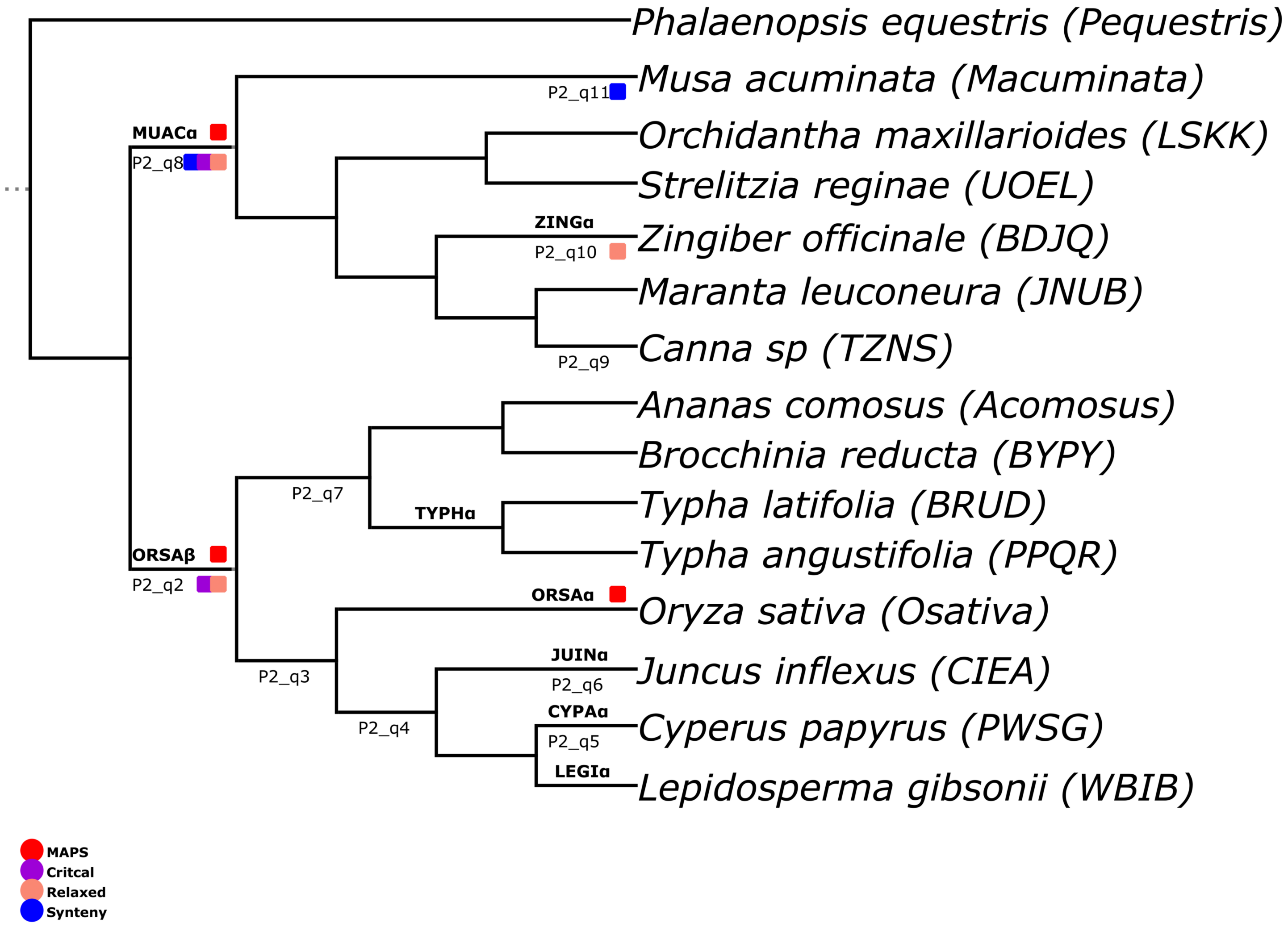


### Figure S39

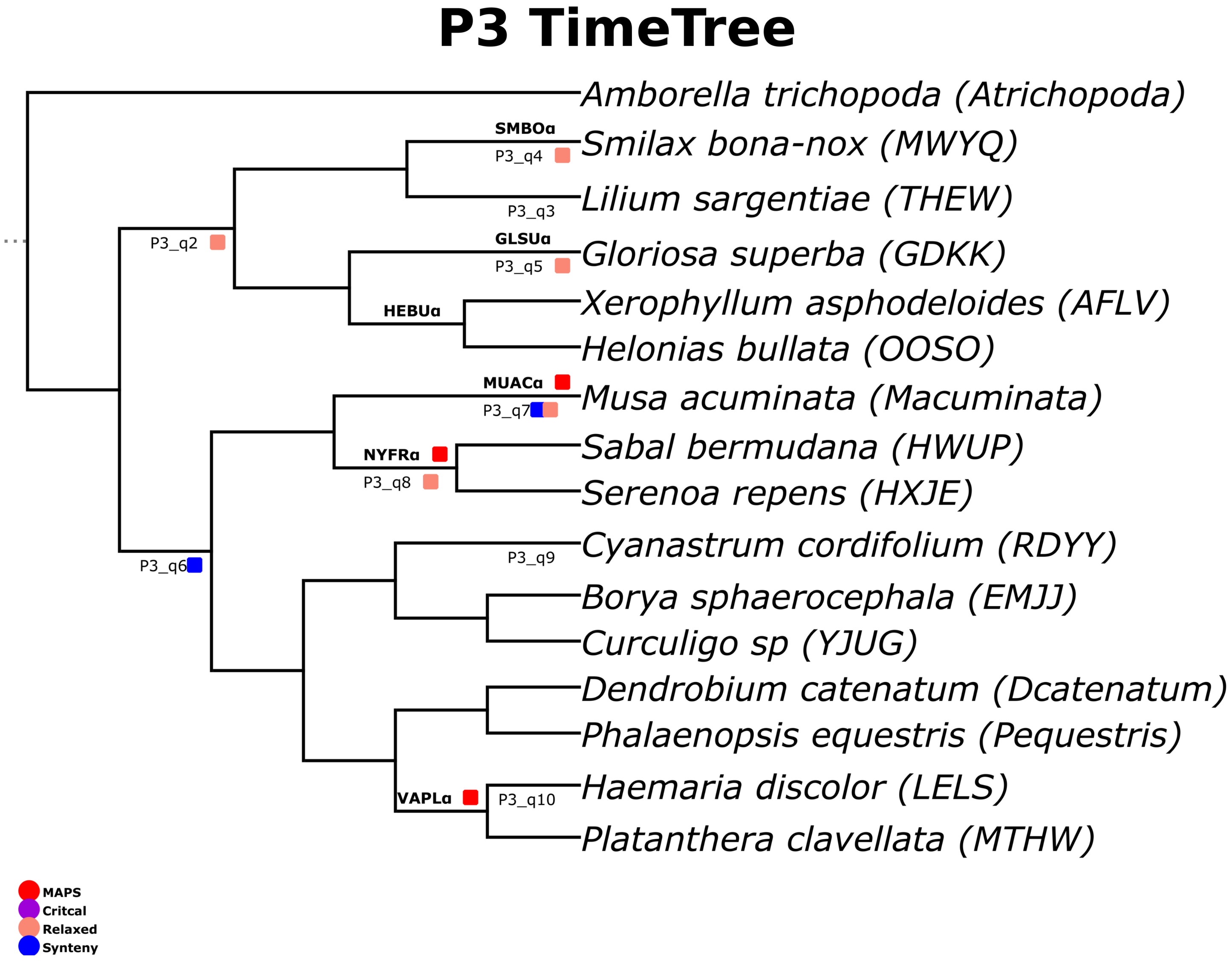
P3 TimeTree
Amborella trichopoda (Atrichopoda)
SMBOα
Smilax bona-nox (MWYQ)
P3_q4
Lilium sargentiae (THEW)
P3_q3
P3_q2
GLSUα
Gloriosa superba (GDKK)
P3_q5
Xerophyllum asphodeloides (AFLV)
HEBUα
Helonias bullata (OOSO)
MUACα
Musa acuminata (Macuminata)
P3_q7
Sabal bermudana (HWUP)
NYFRα
P3_q8
Serenoa repens (HXJE)
Cyanastrum cordifolium (RDYY)
P3_q9
P3_q6
Borya sphaerocephala (EMJJ)
Curculigo sp (YJUG)
Dendrobium catenatum (Dcatenatum)
Phalaenopsis equestris (Pequestris)
Haemaria discolor (LELS)
VAPLα
P3_q10
Platanthera clavellata (MTHW)
MAPS
Critcal
Relaxed
Synteny

### Figure S40

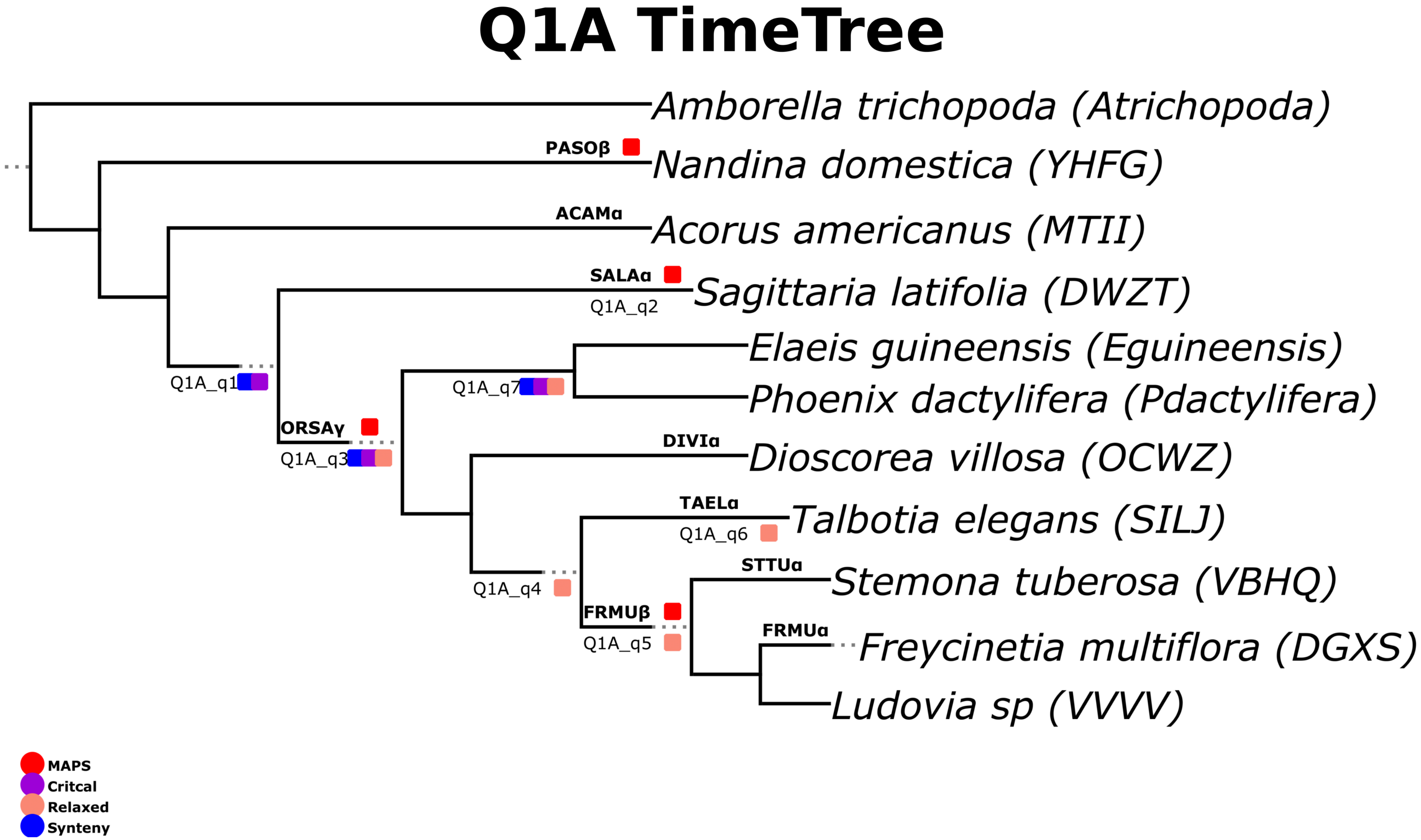
Q1A TimeTree
Amborella trichopoda (Atrichopoda)
PASOβ
Nandina domestica (YHFG)
ACAMα
Acorus americanus (MTII)
SALAα
Q1A_q2
Sagittaria latifolia (DWZT)
Elaeis guineensis (Eguineensis)
Phoenix dactylifera (Pdactylifera)
Q1A_q1
Q1A_q7
ORSAγ
Q1A_q3
DIVIα
Dioscorea villosa (OCWZ)
TAELα
Talbotia elegans (SILJ)
Q1A_q6
STTUα
Stemona tuberosa (VBHQ)
Q1A_q4
FRMUβ
FRMUα
Freycinetia multiflora (DGXS)
Q1A_q5
Ludovia sp (VVVV)
MAPS
Critcal
Relaxed
Synteny

### Figure S41

### Q1B TimeTree

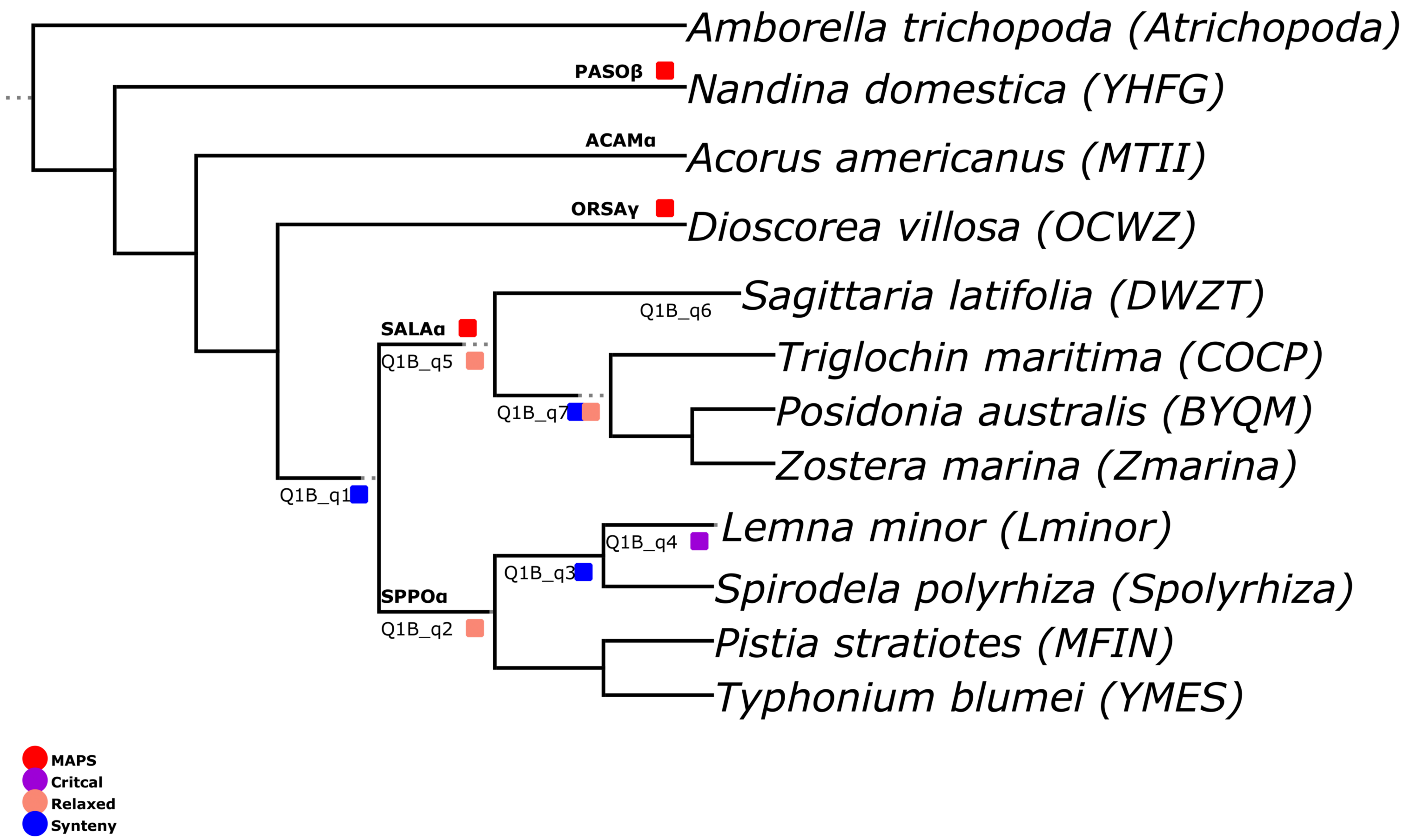


### Figure S42

### Q2 TimeTree

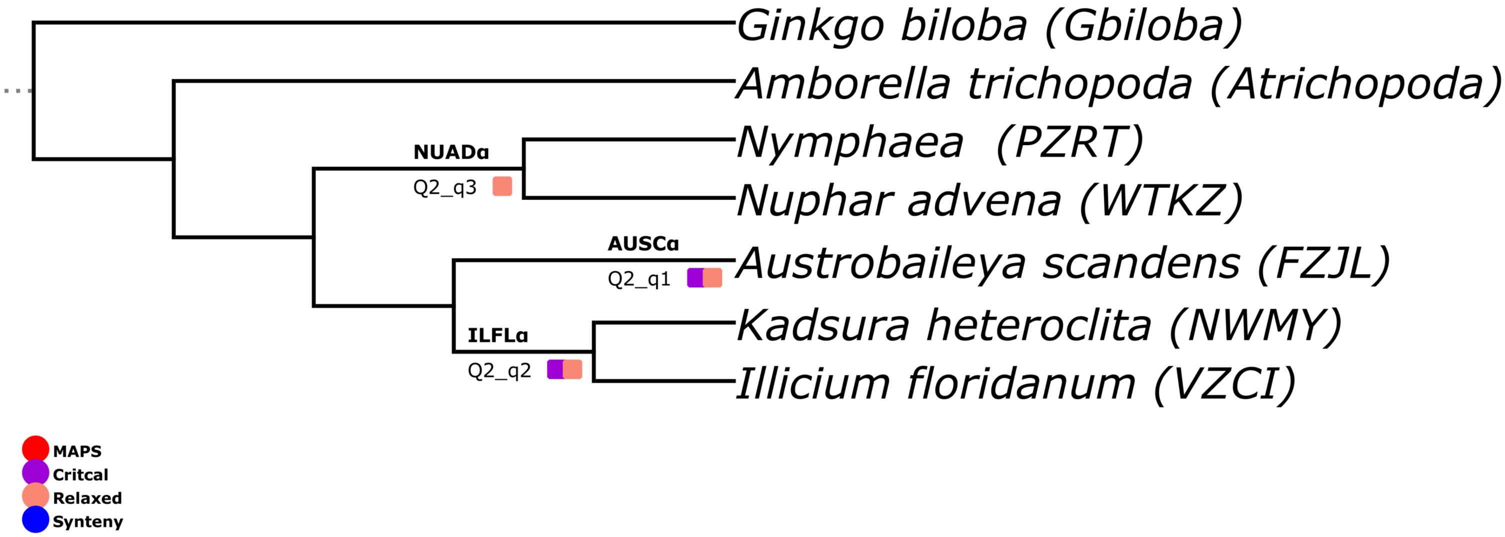


**Figure S43**

### A1 Janssens

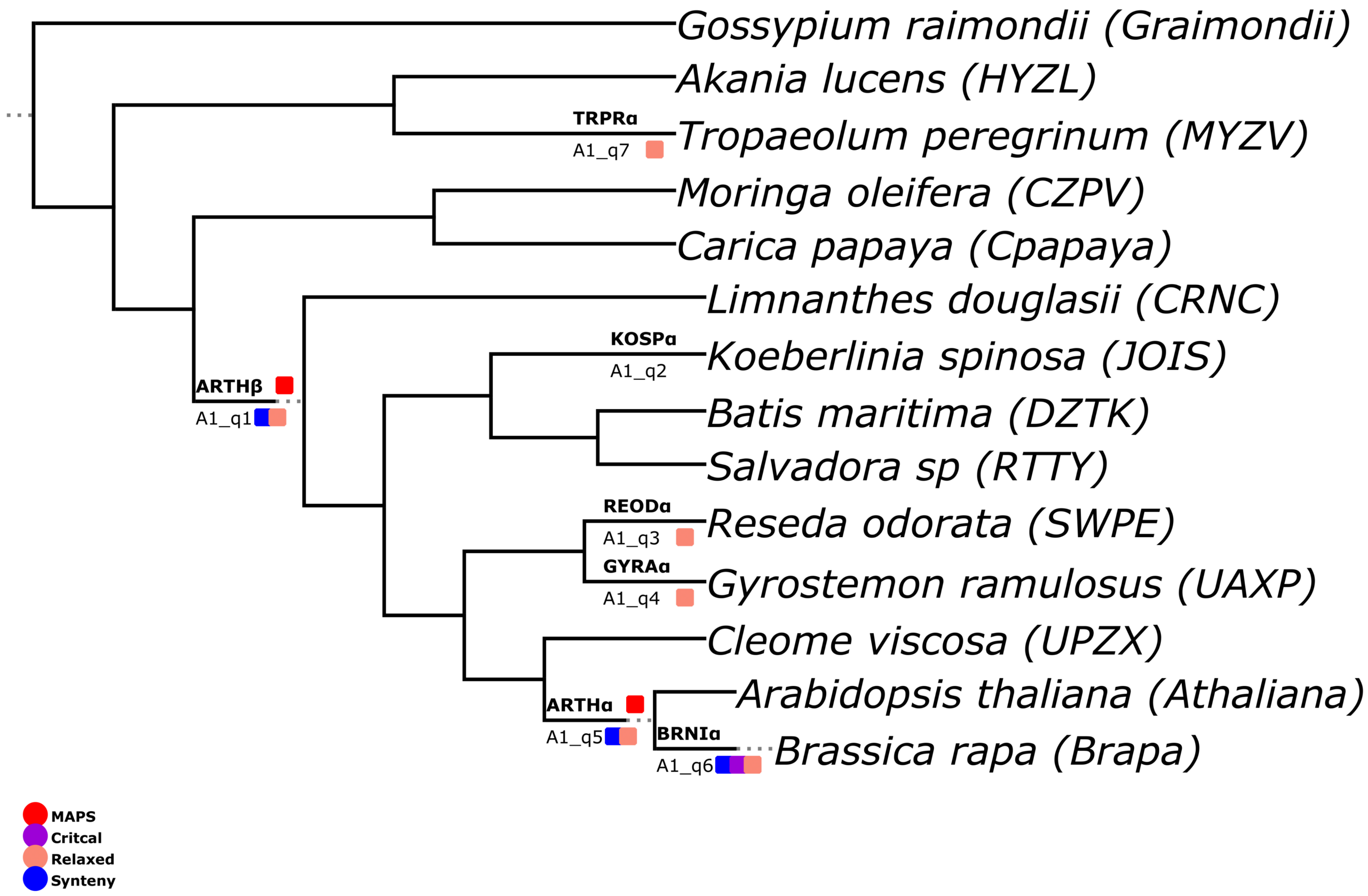


### Figure S44

### A2 Janssens

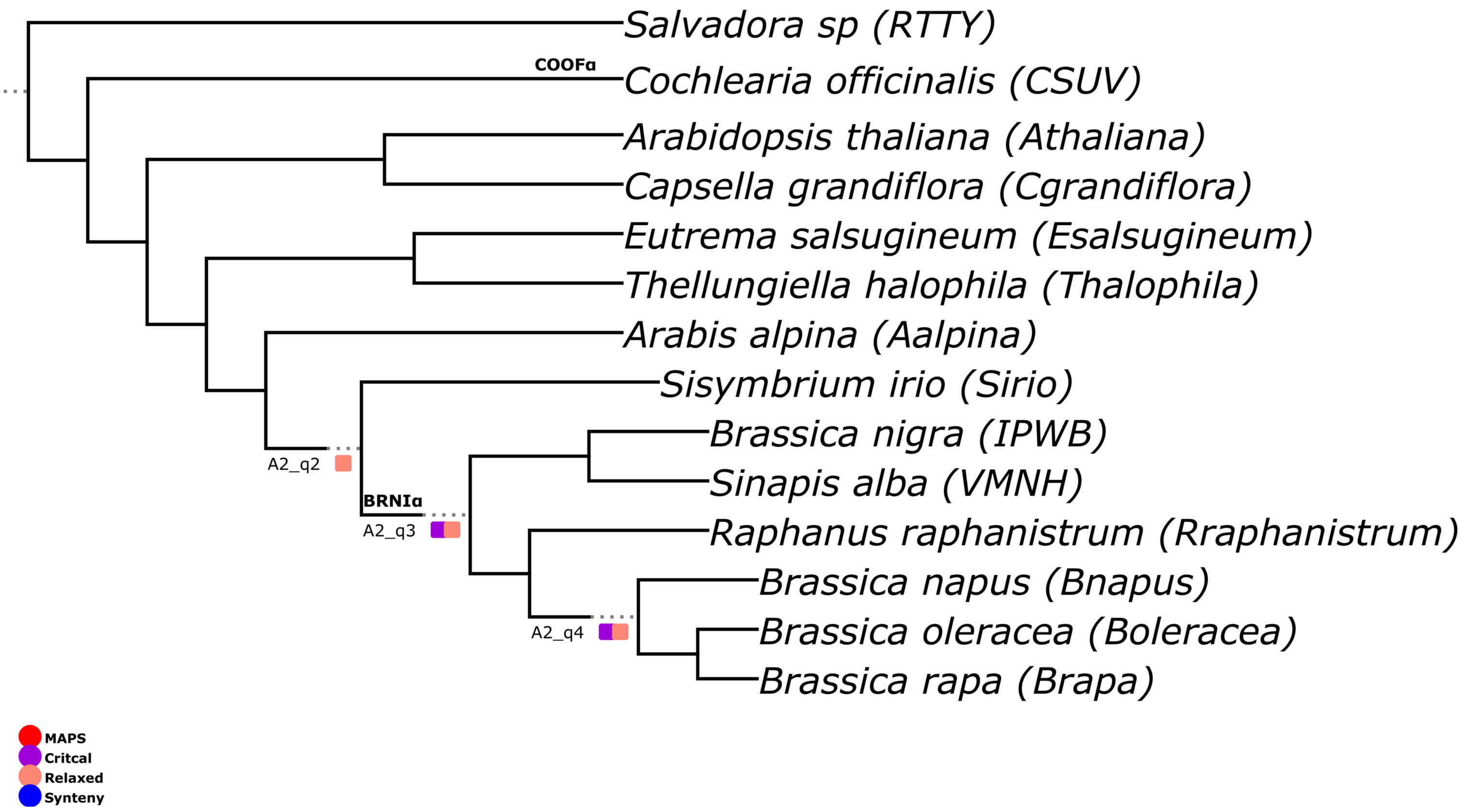


### Figure S45

### A3 Janssens

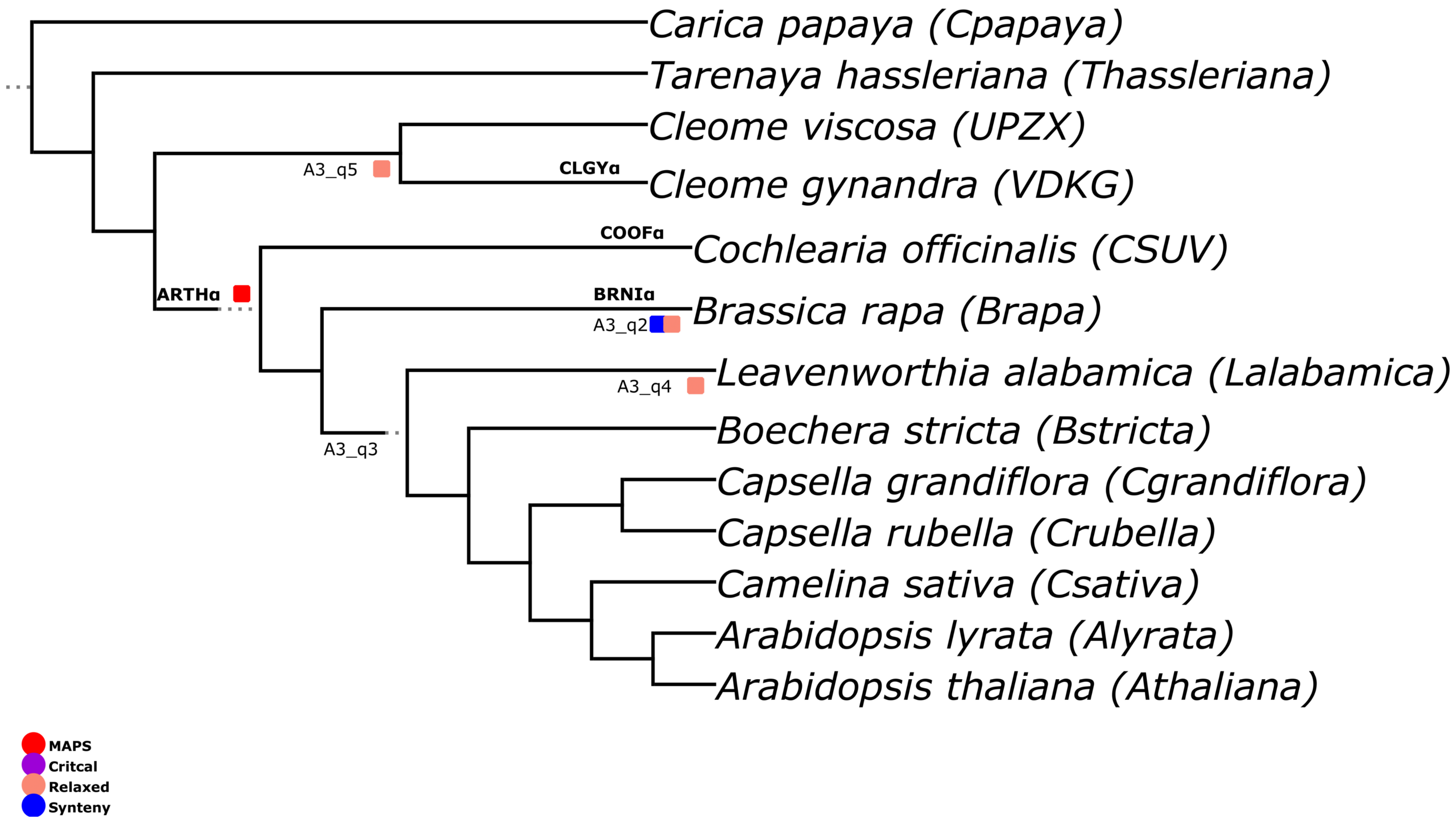


### Figure S46

### B1 Janssens

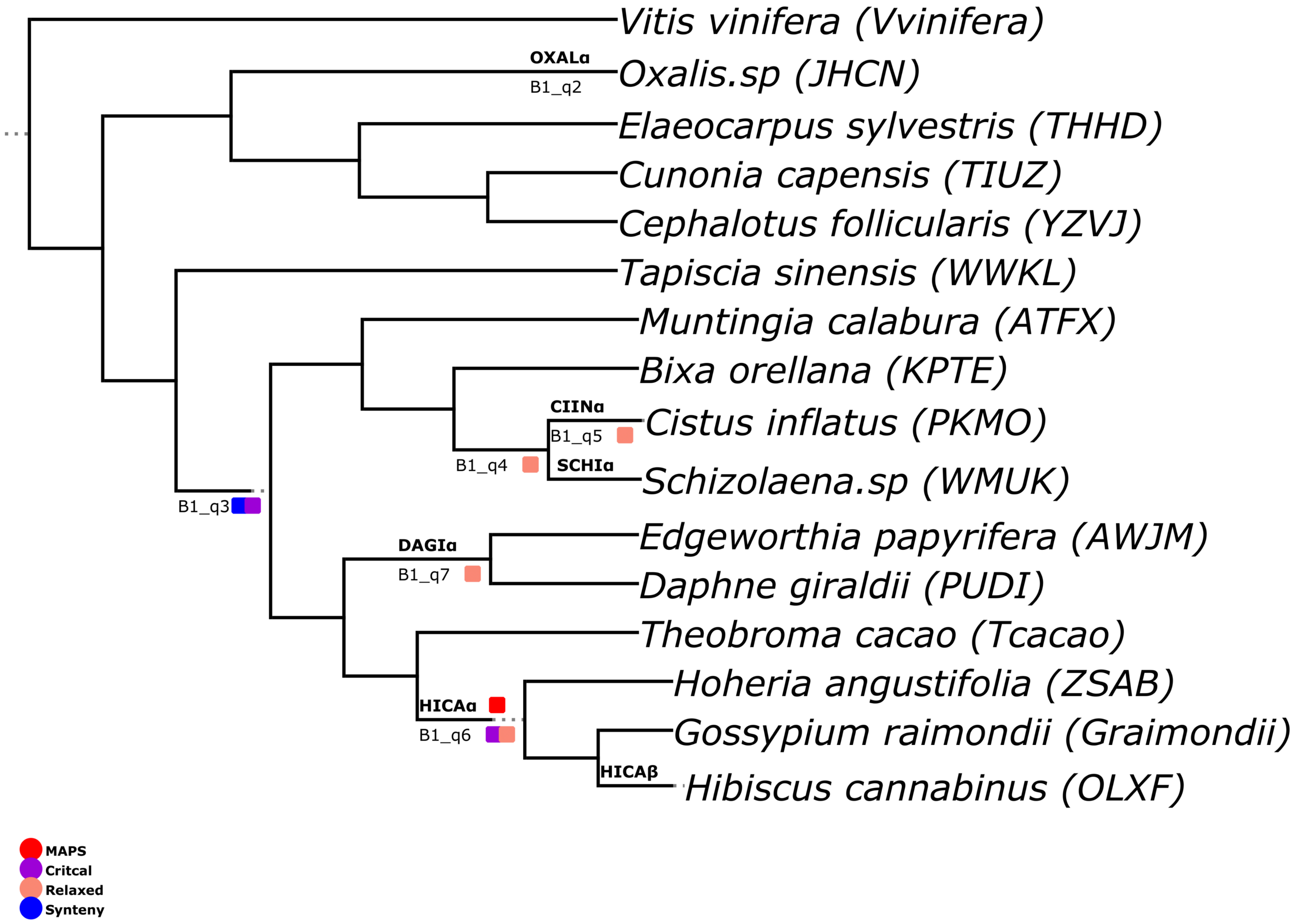


### Figure S47

### B2 Janssens

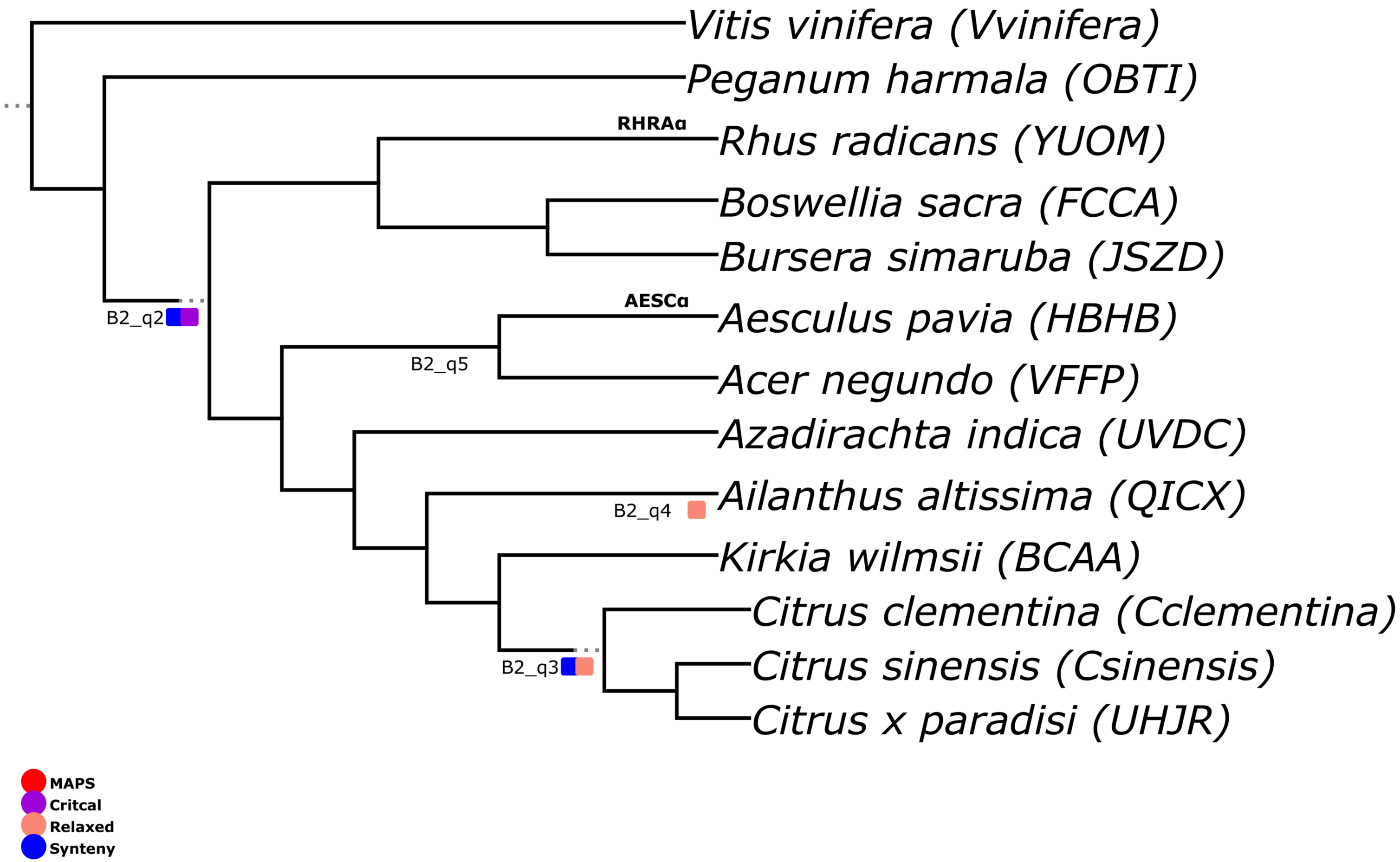


### Figure S48

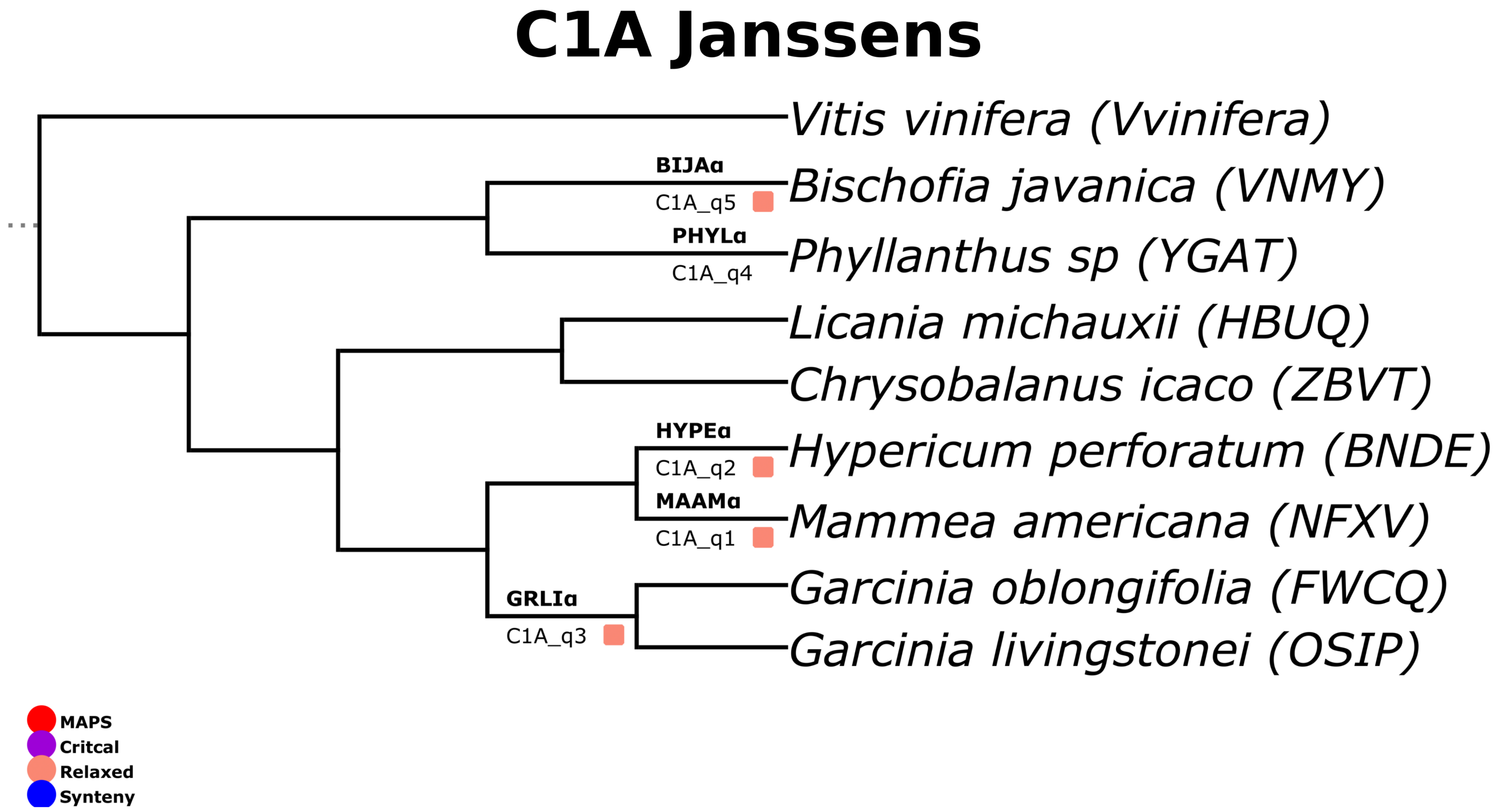
C1A Janssens
Vitis vinifera (Vvinifera)
BIJAα
C1A_q5
Bischofia javanica (VNMY)
PHYLα
C1A_q4
Phyllanthus sp (YGAT)
Licania michauxii (HBUQ)
Chrysobalanus icaco (ZBVT)
HYPEα
C1A_q2
Hypericum perforatum (BNDE)
MAAMα
C1A_q1
Mammea americana (NFXV)
GRLIα
C1A_q3
Garcinia oblongifolia (FWCQ)
Garcinia livingstonei (OSIP)
MAPS
Critcal
Relaxed
Synteny

### Figure S49

### C1B Janssens

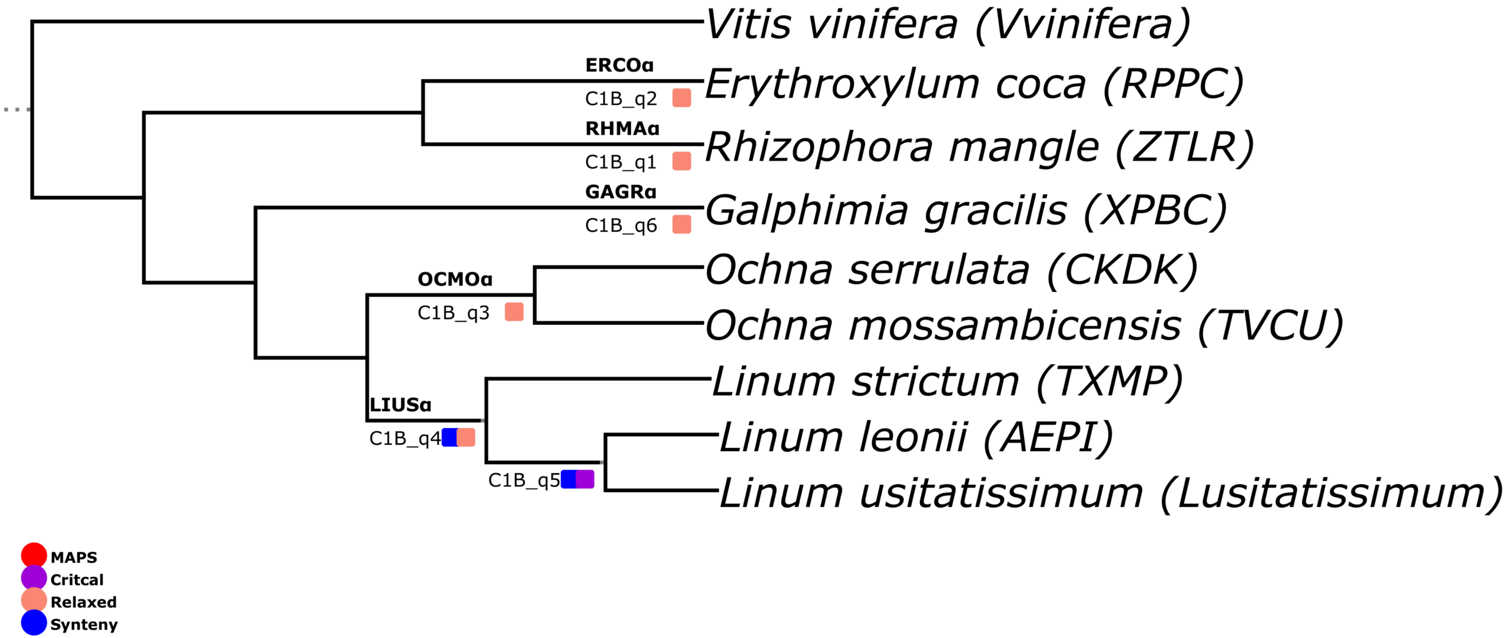


### Figure S50

### C2A Janssens

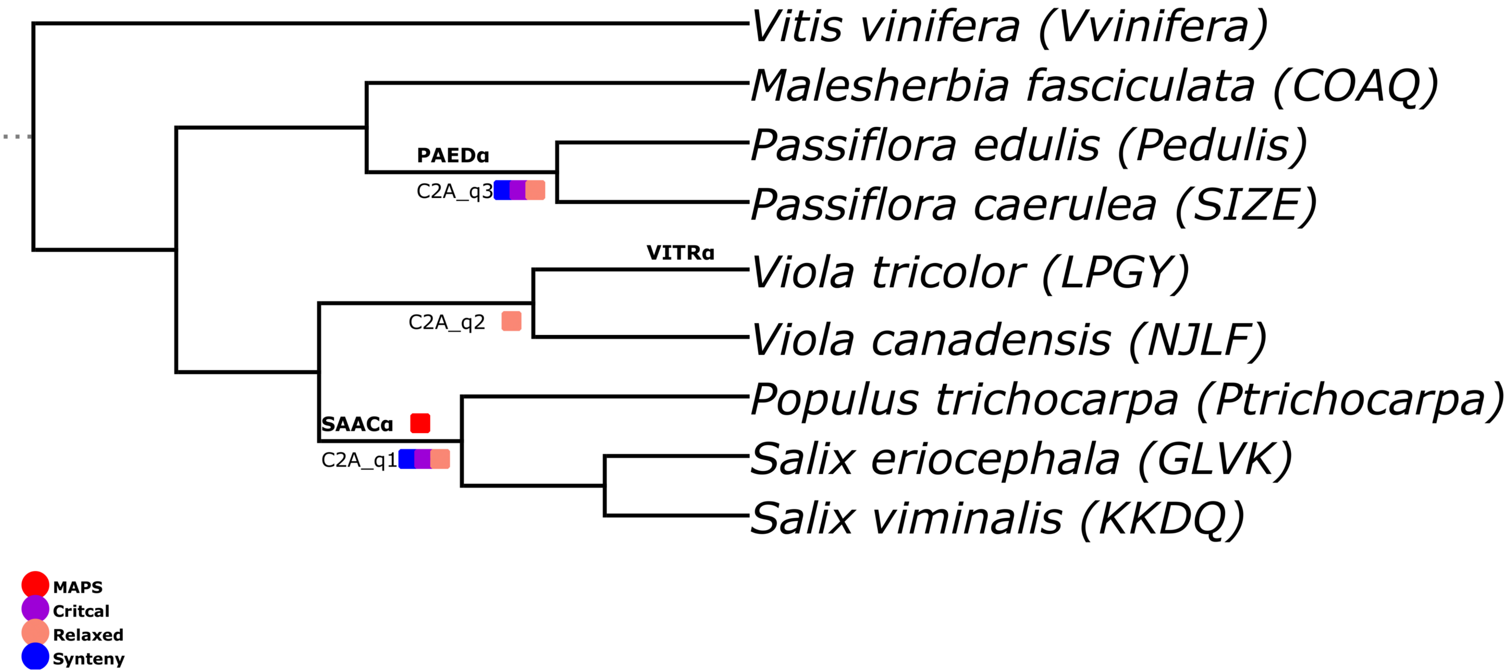


### Figure S51

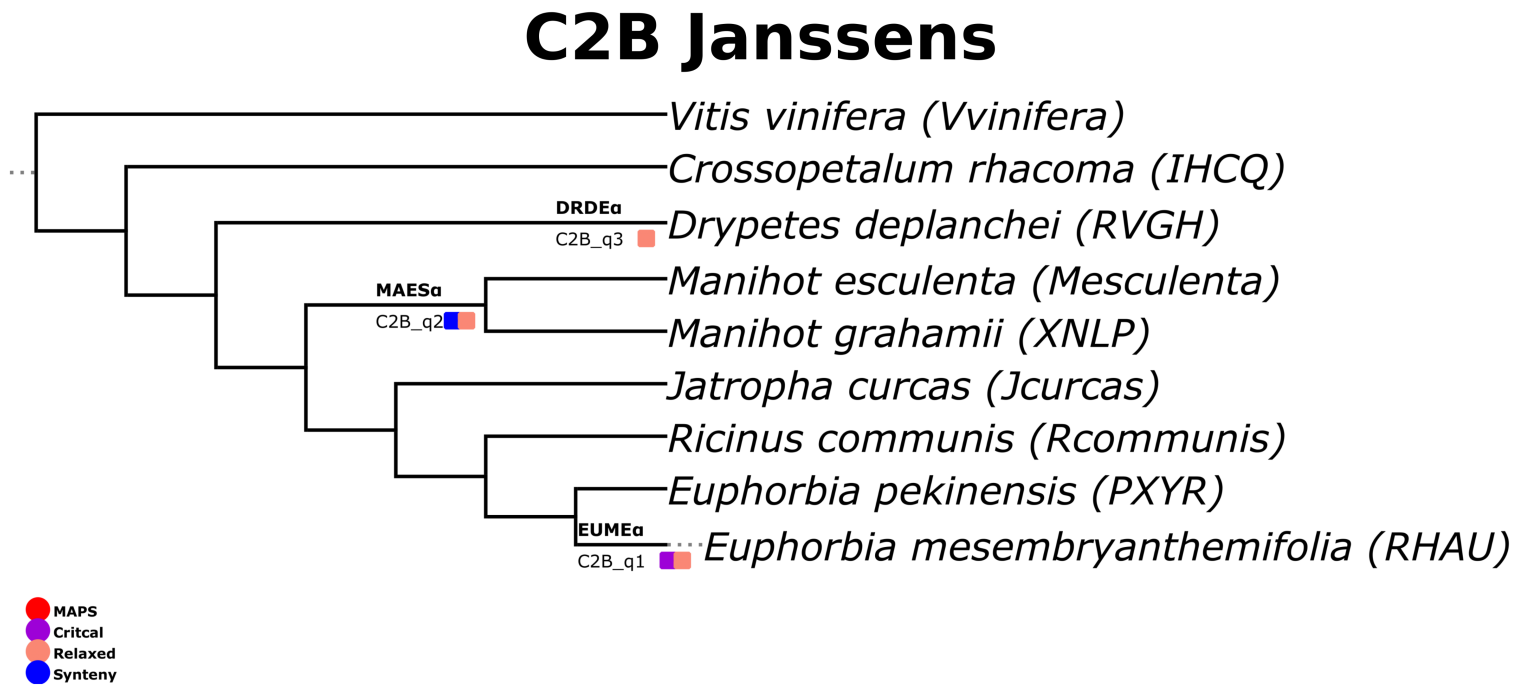
C2B Janssens
Vitis vinifera (Vvinifera)
Crossopetalum rhacoma (IHCQ)
DRDEα
C2B_q3
Drypetes deplanchei (RVGH)
MAESα
C2B_q2
Manihot esculenta (Mesculenta)
Manihot grahamii (XNLP)
Jatropha curcas (Jcurcas)
Ricinus communis (Rcommunis)
Euphorbia pekinensis (PXYR)
EUMEα
C2B_q1
Euphorbia mesembryanthemifolia (RHAU)
MAPS
Critcal
Relaxed
Synteny

### Figure S52

### D Janssens

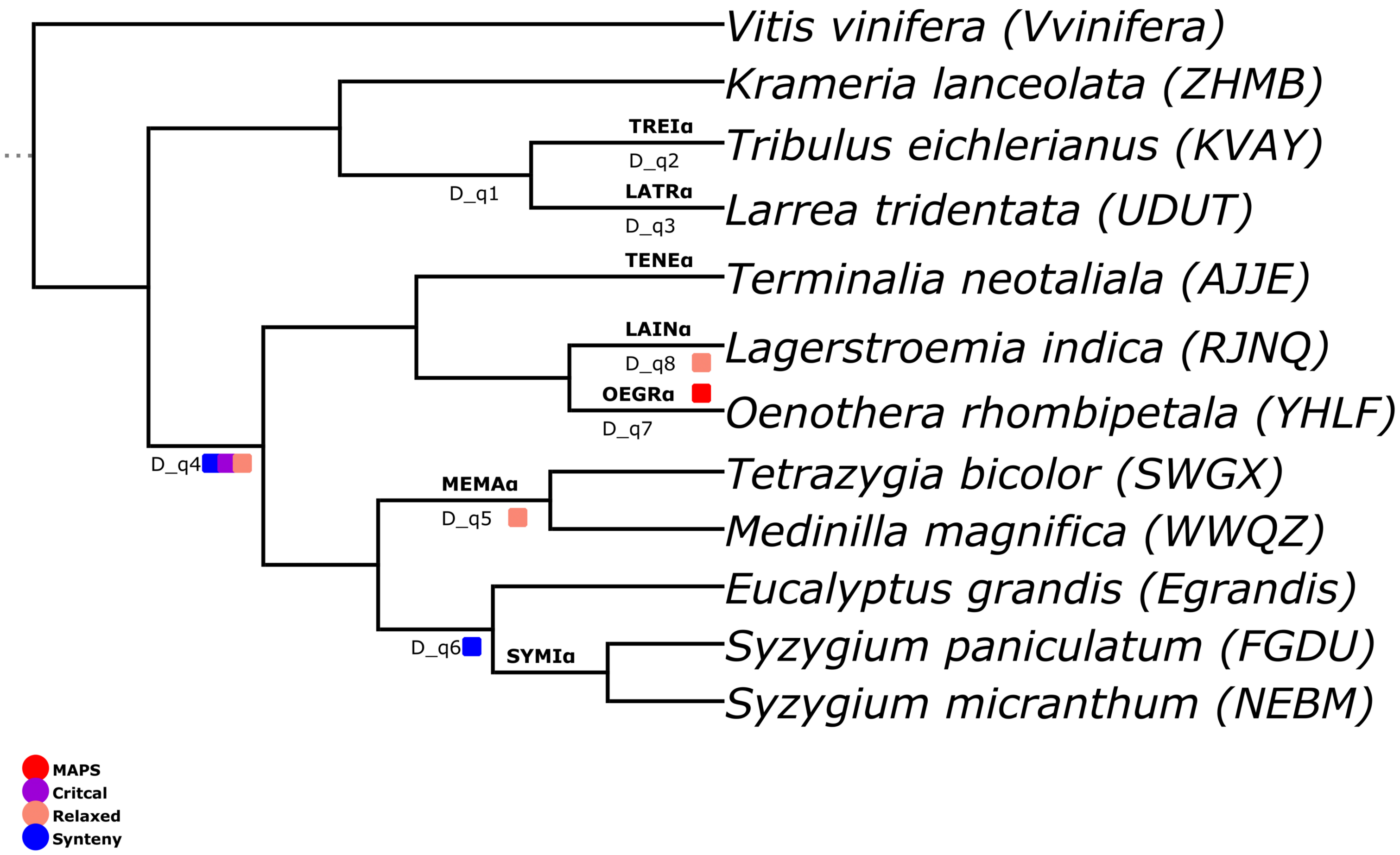


### Figure S53

### E1 Janssens

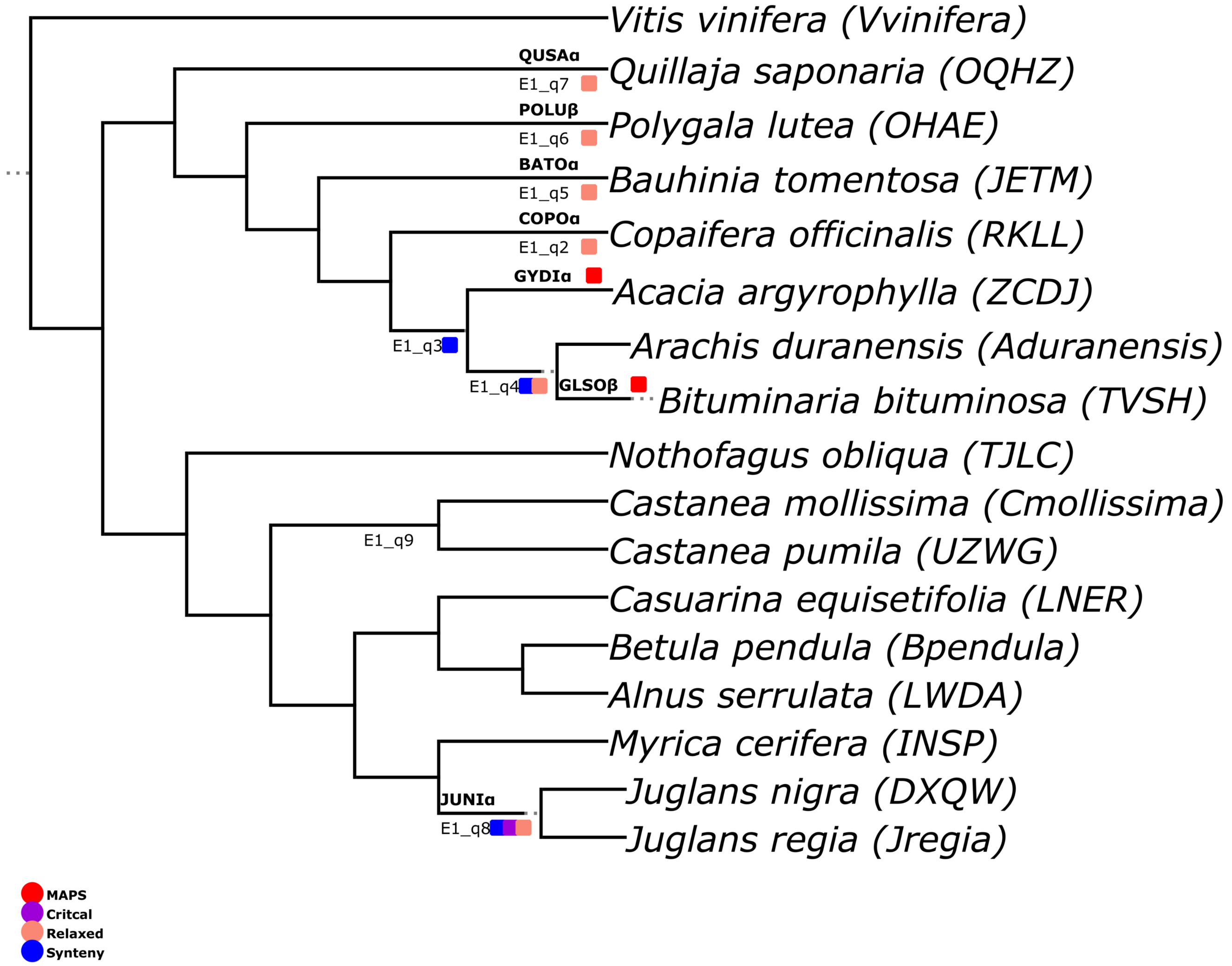


### Figure S54

### E2 Janssens

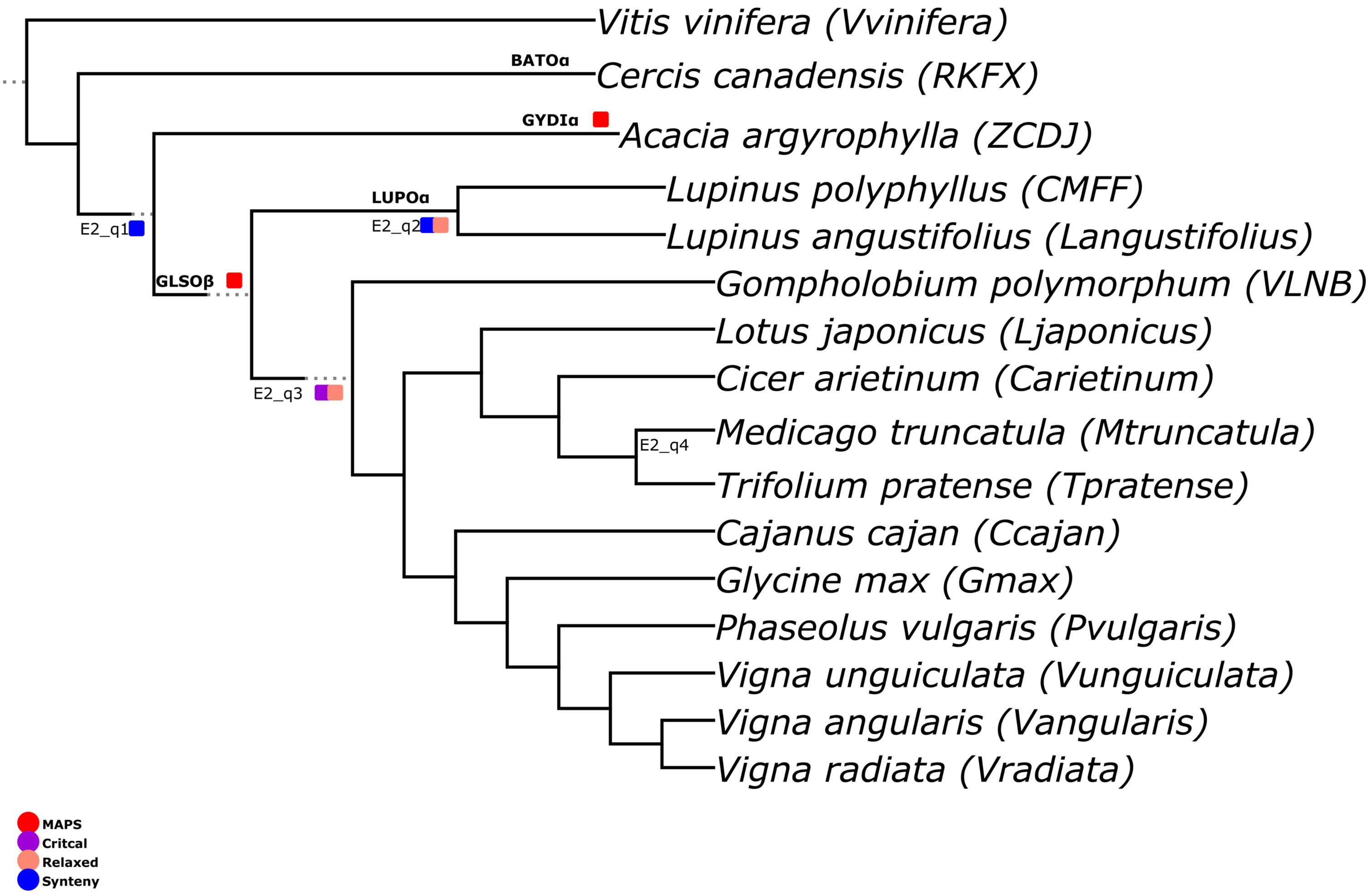


### Figure S55

### F1 Janssens

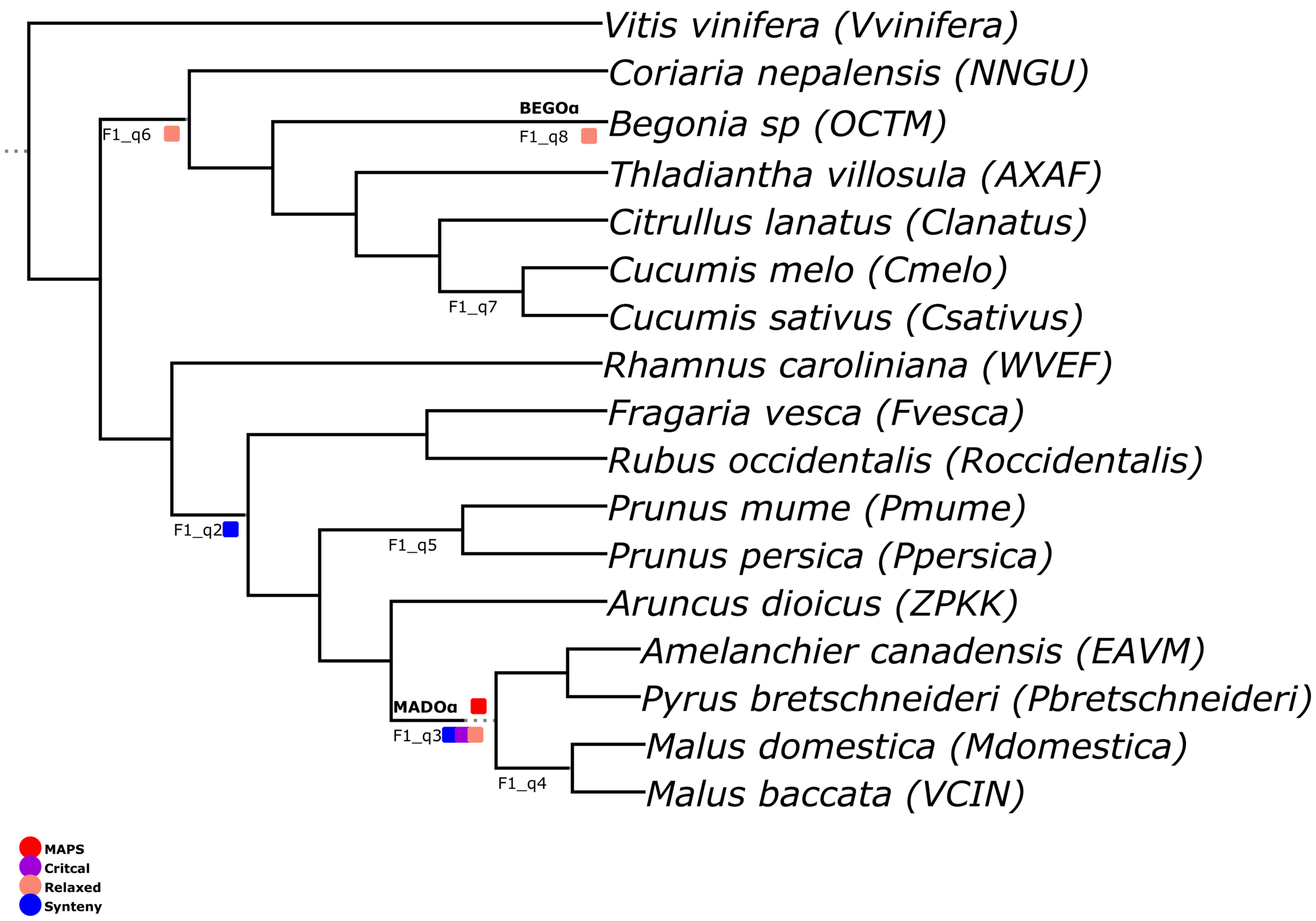


### Figure S56

### F2 Janssens

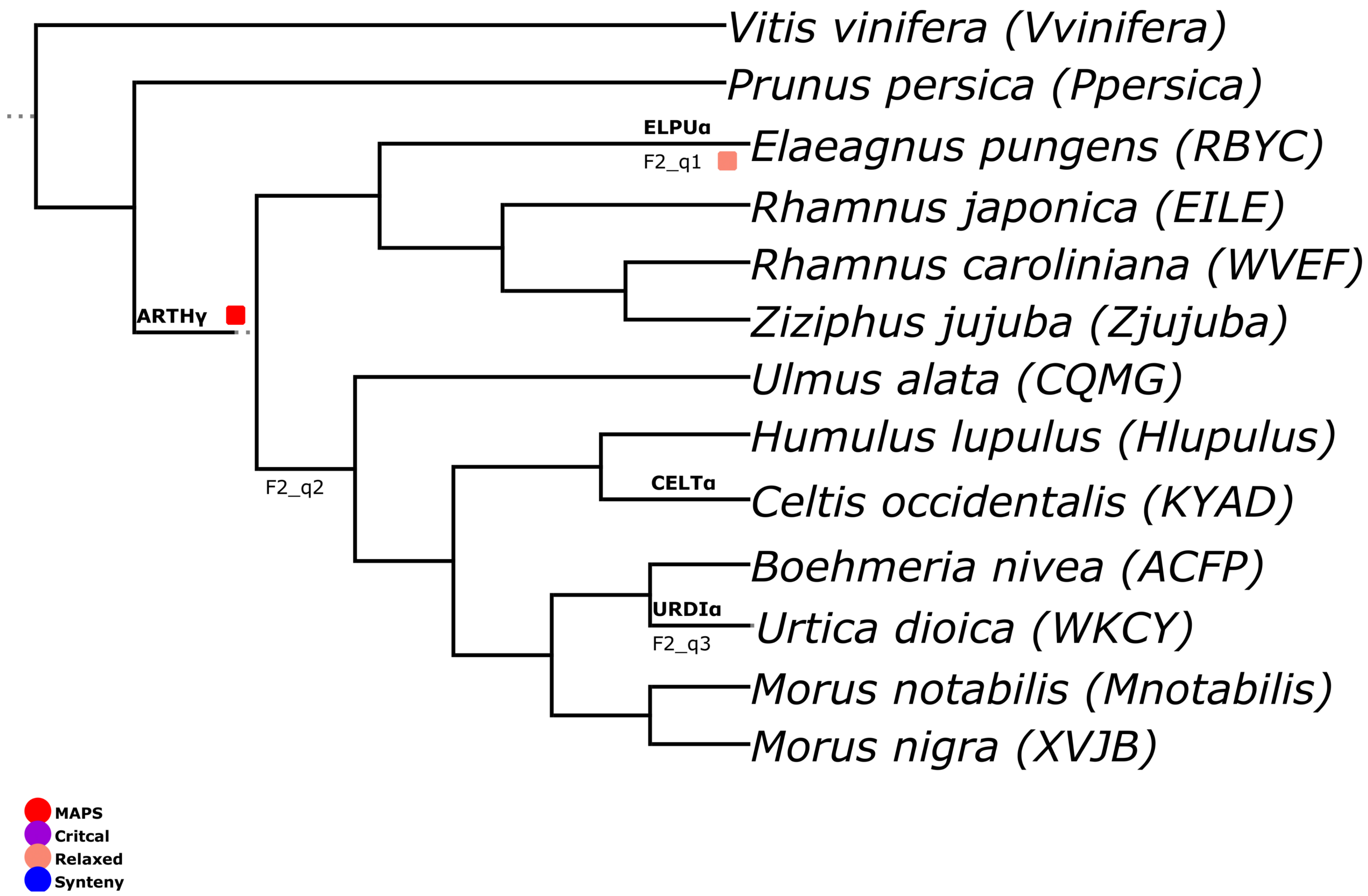


### Figure S57

### G1 Janssens

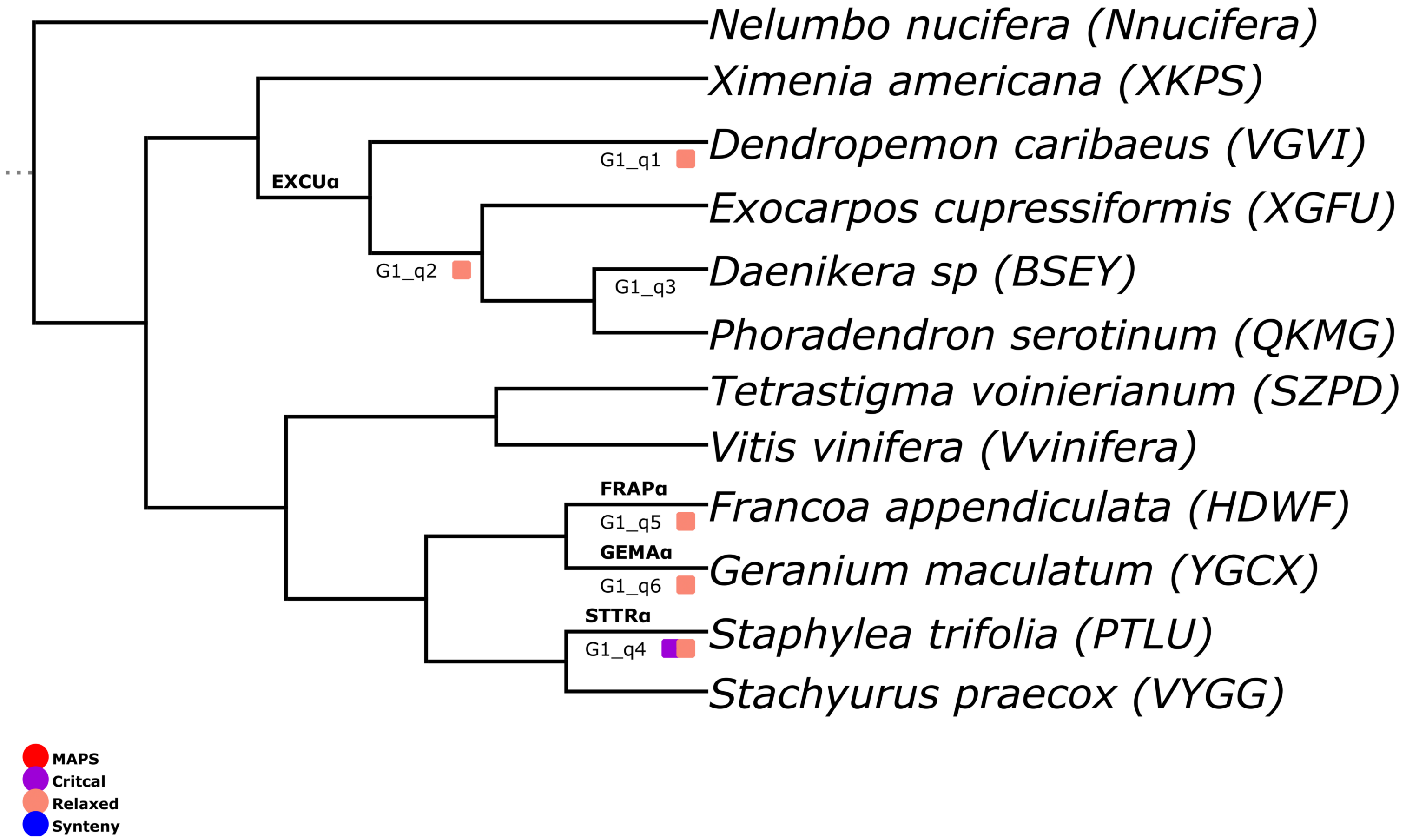


### Figure S58

### G2 Janssens

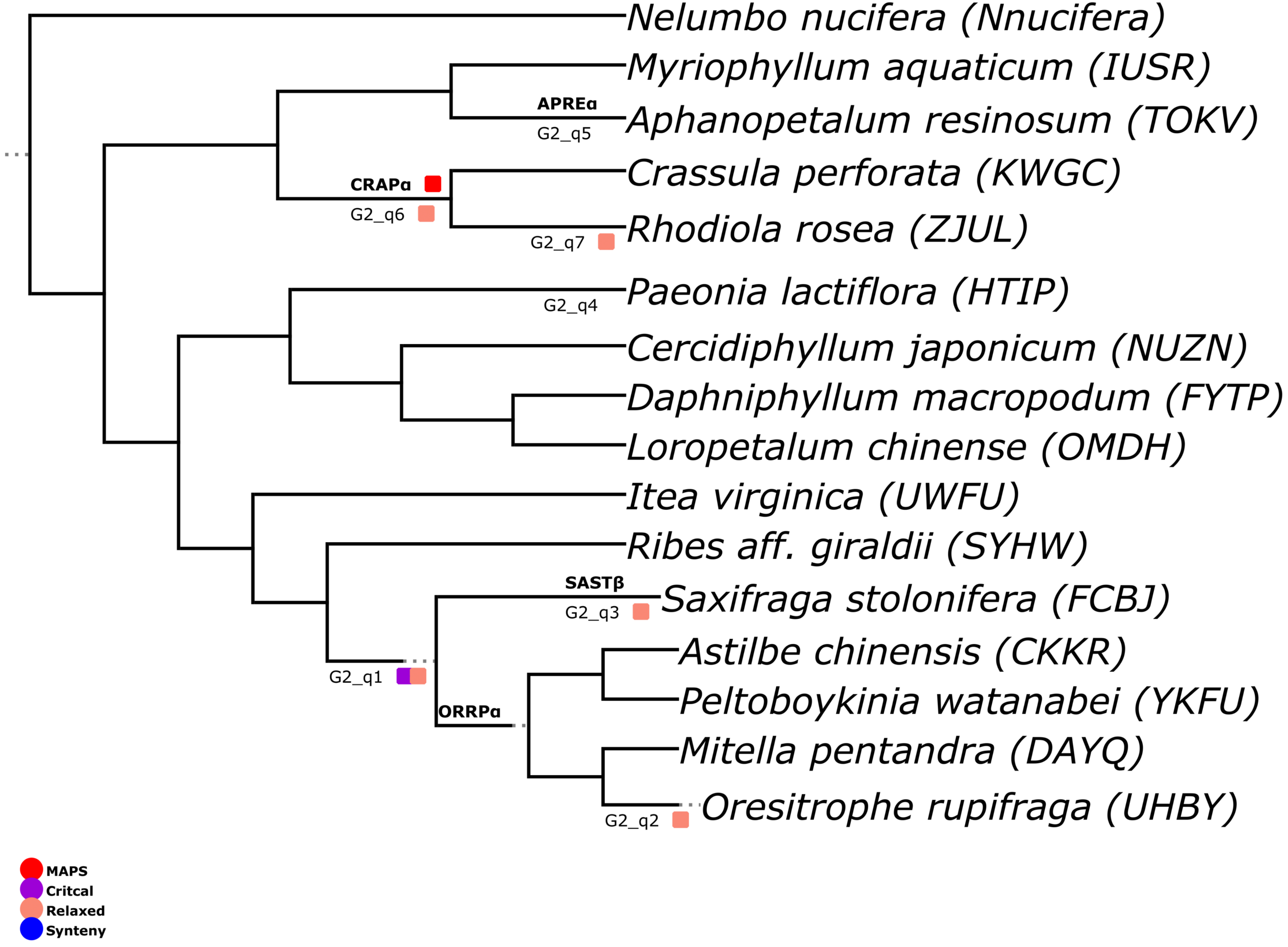


### Figure S59

### H1 Janssens

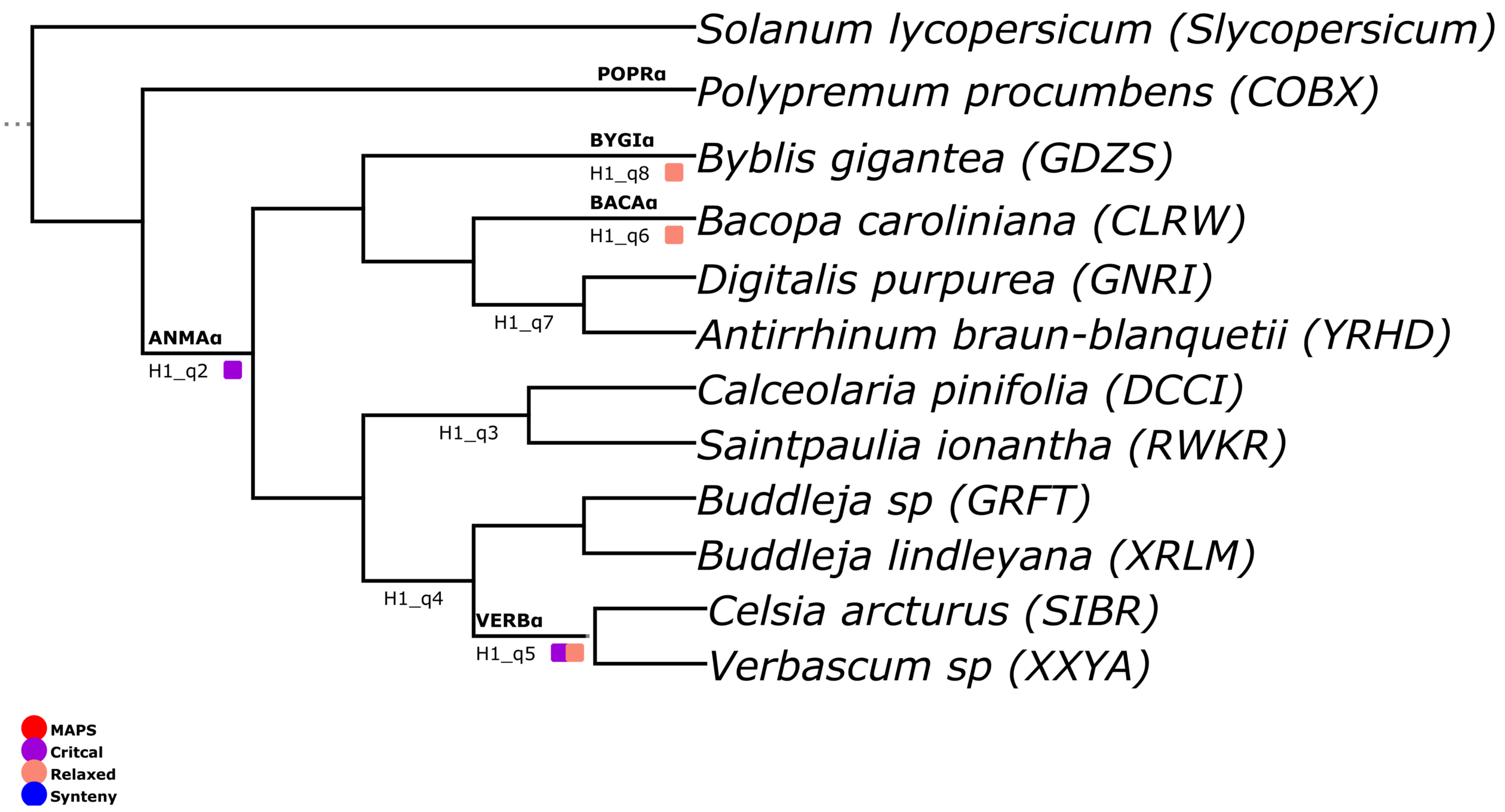


### Figure S60

### H2 Janssens

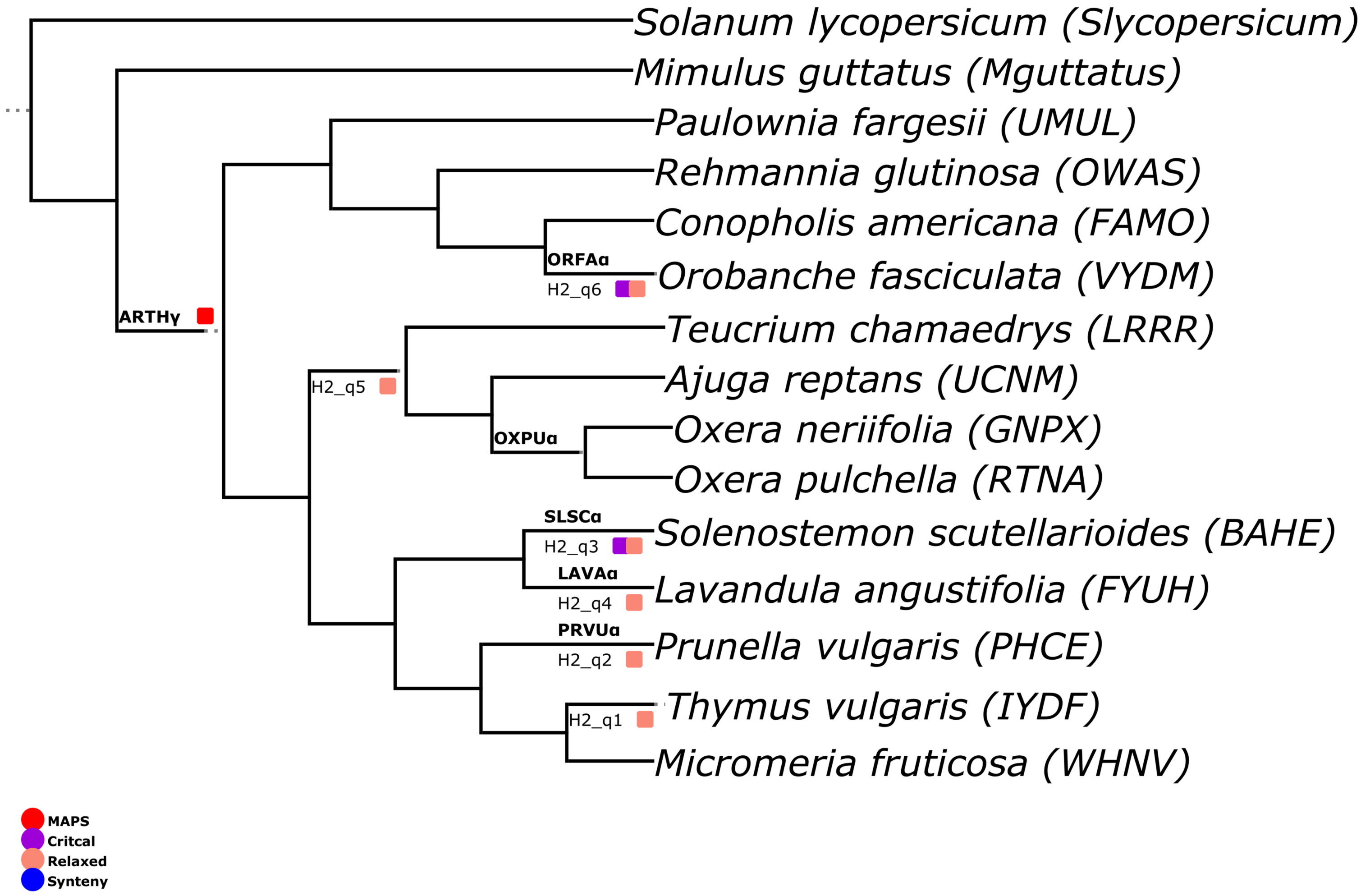


### Figure S61

### H3 Janssens

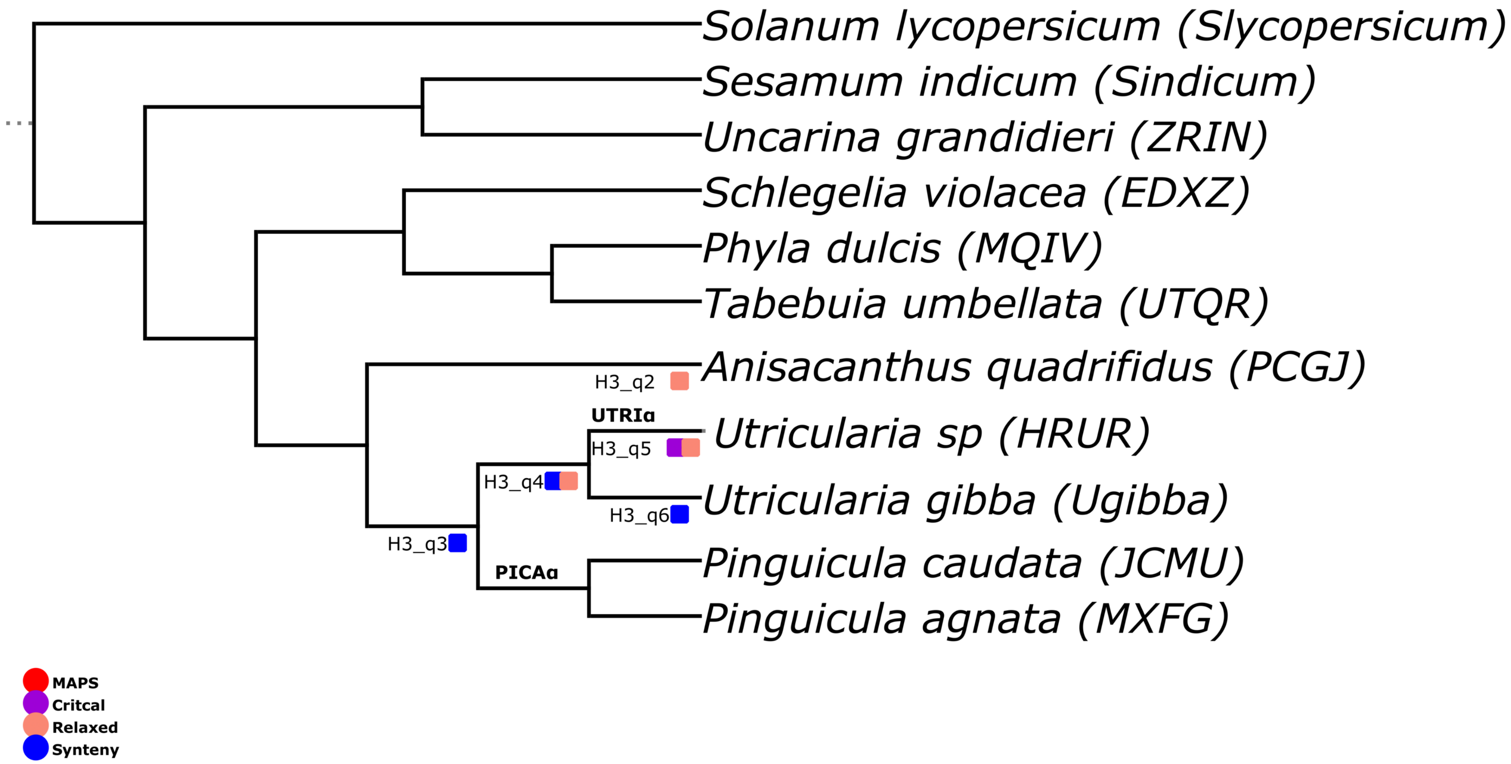


### Figure S62

### I1A Janssens

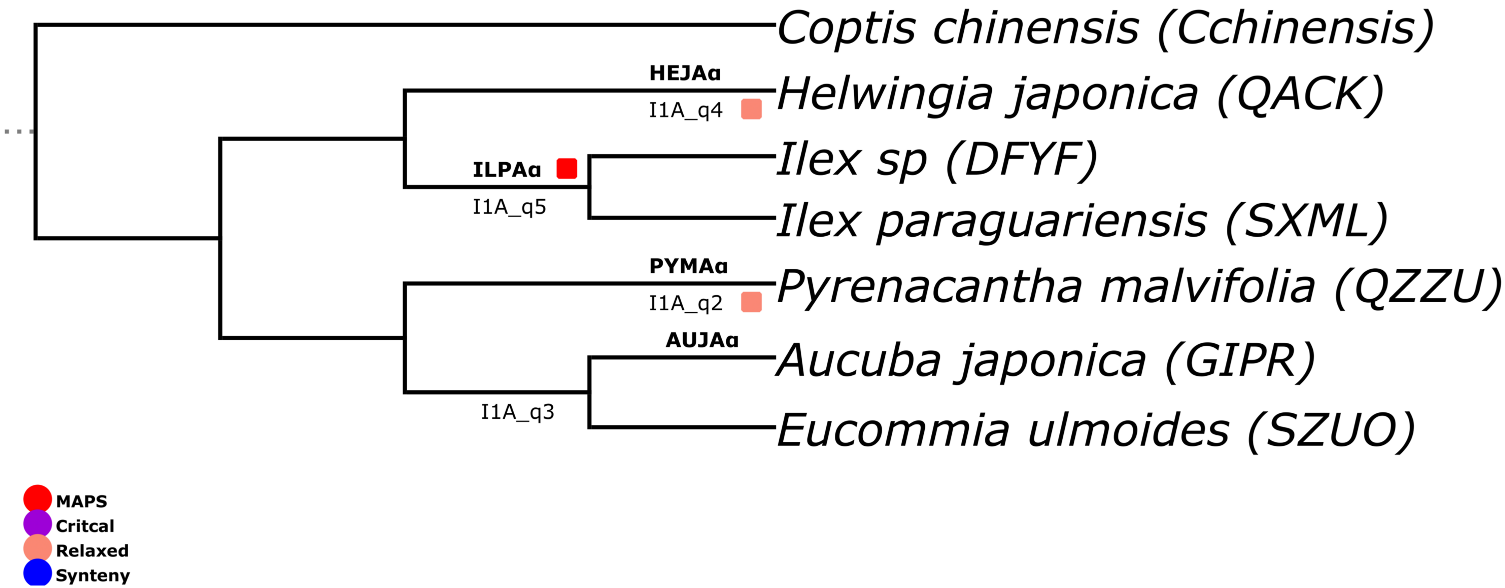


### Figure S63

### I1B Janssens

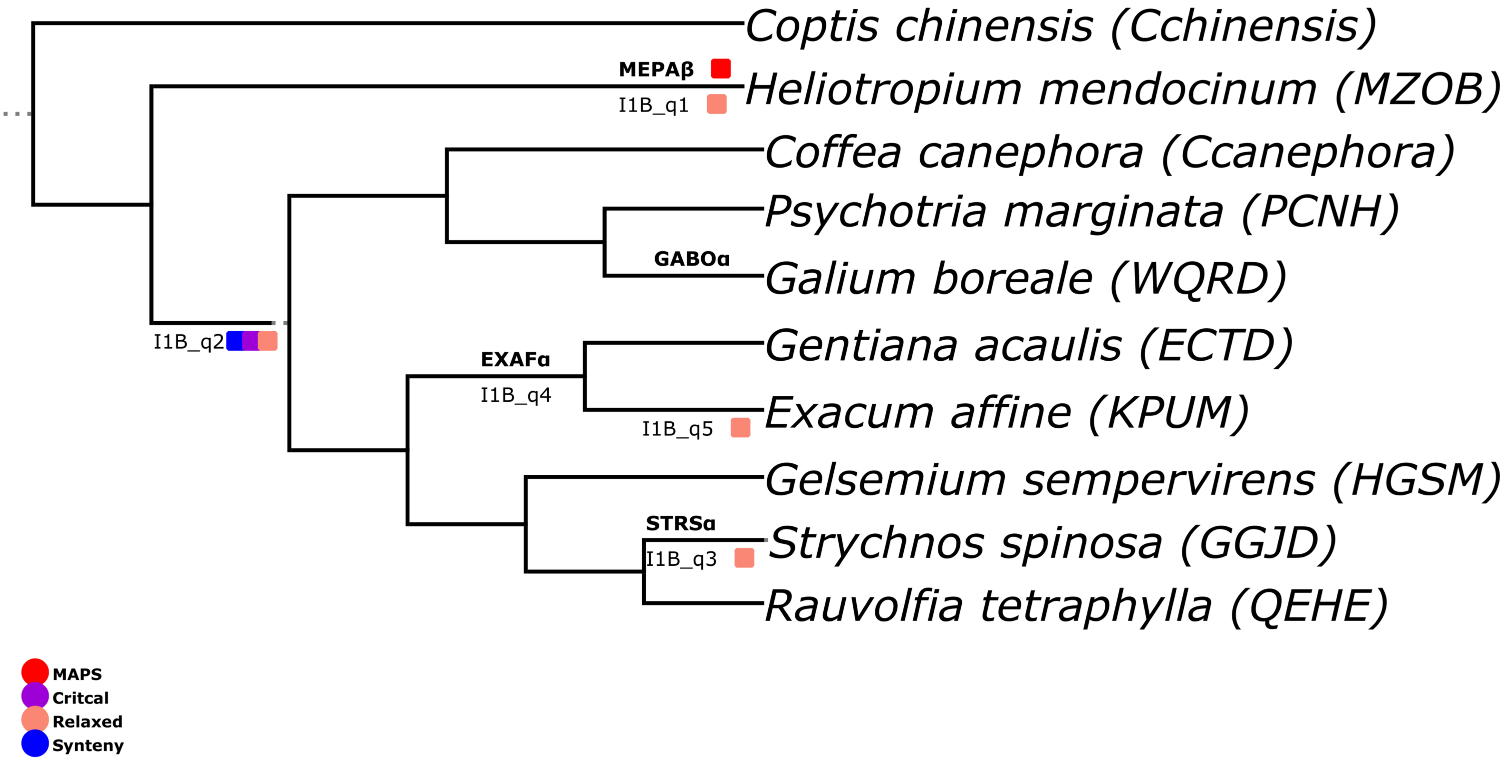


### Figure S64

### I2A Janssens

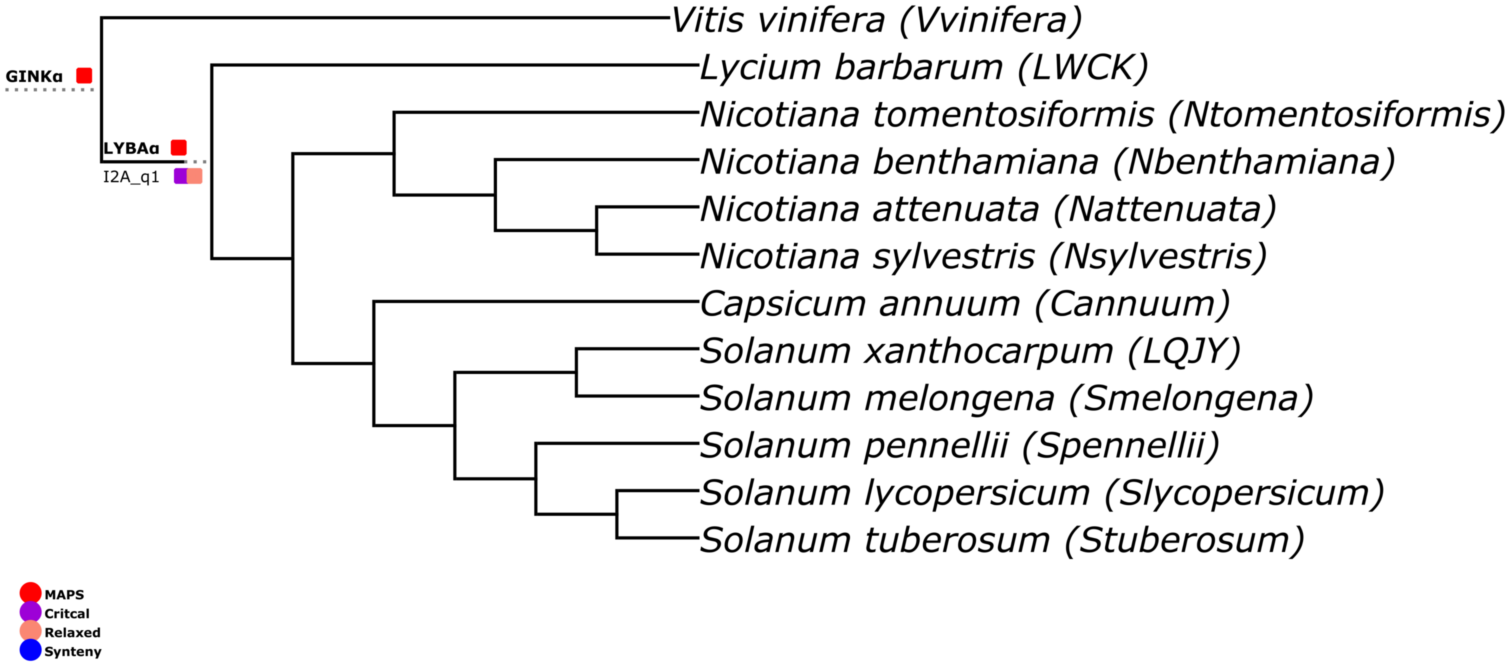


### Figure S65

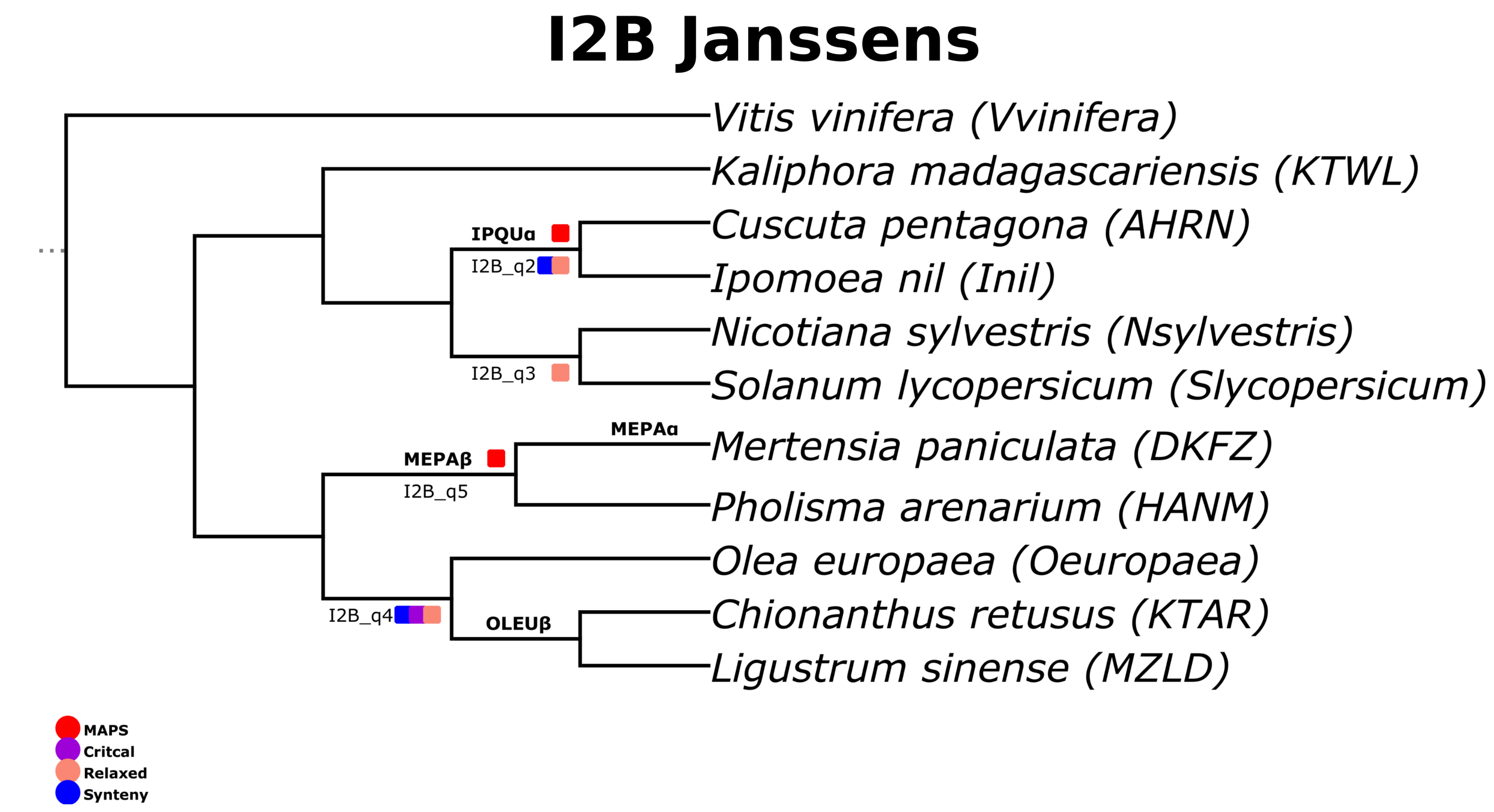
I2B Janssens
Vitis vinifera (Vvinifera)
Kaliphora madagascariensis (KTWL)
Cuscuta pentagona (AHRN)
Ipomoea nil (Inil)
Nicotiana sylvestris (Nsylvestris)
Solanum lycopersicum (Slycopersicum)
Mertensia paniculata (DKFZ)
Pholisma arenarium (HANM)
Olea europaea (Oeuropaea)
Chionanthus retusus (KTAR)
Ligustrum sinense (MZLD)
IPQUα
I2B_q2
I2B_q3
MEPAα
MEPAβ
I2B_q5
I2B_q4
OLEUβ
MAPS
Critcal
Relaxed
Synteny

### Figure S66

### J1 Janssens

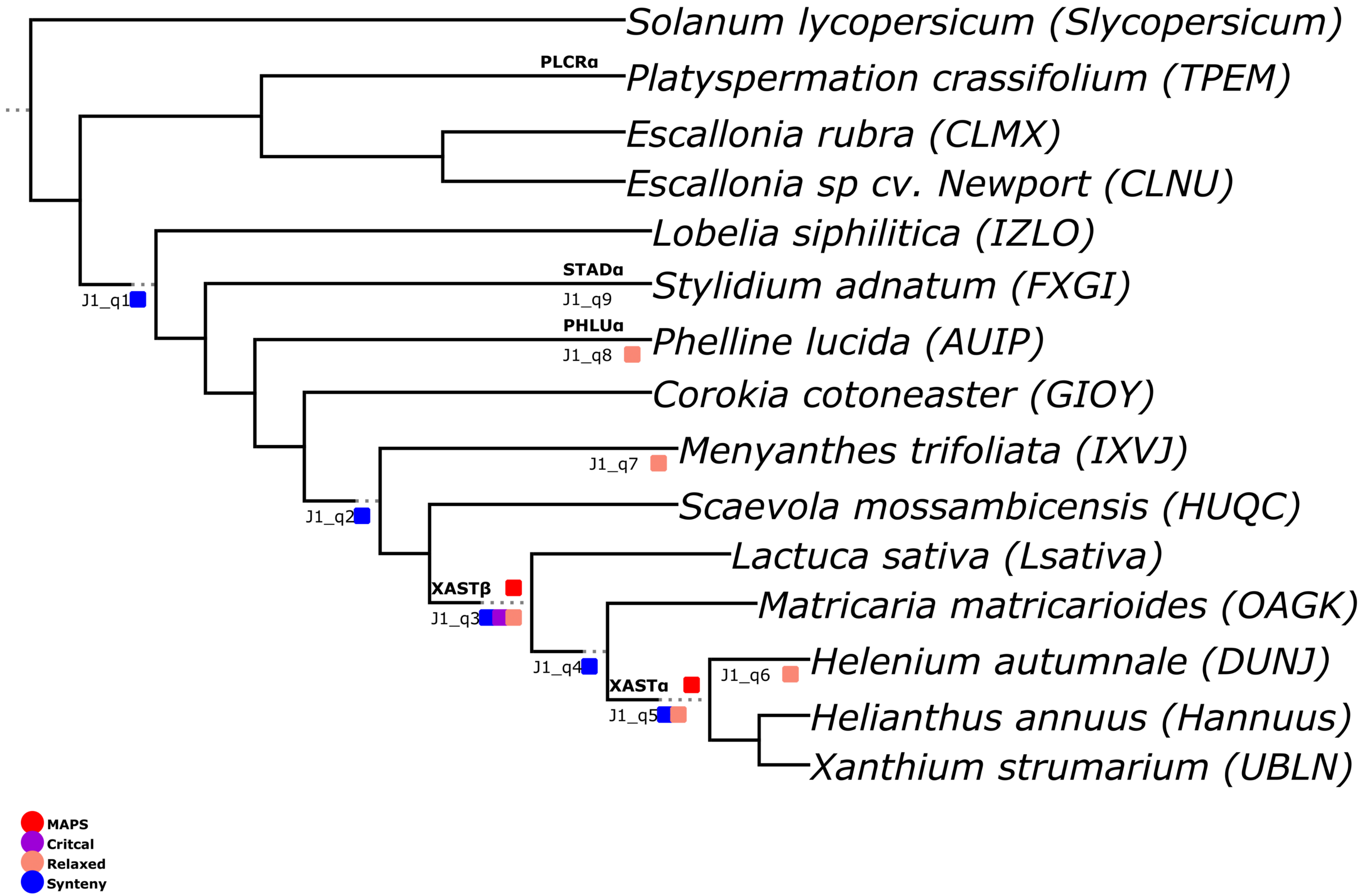


### Figure S67

### J2 Janssens

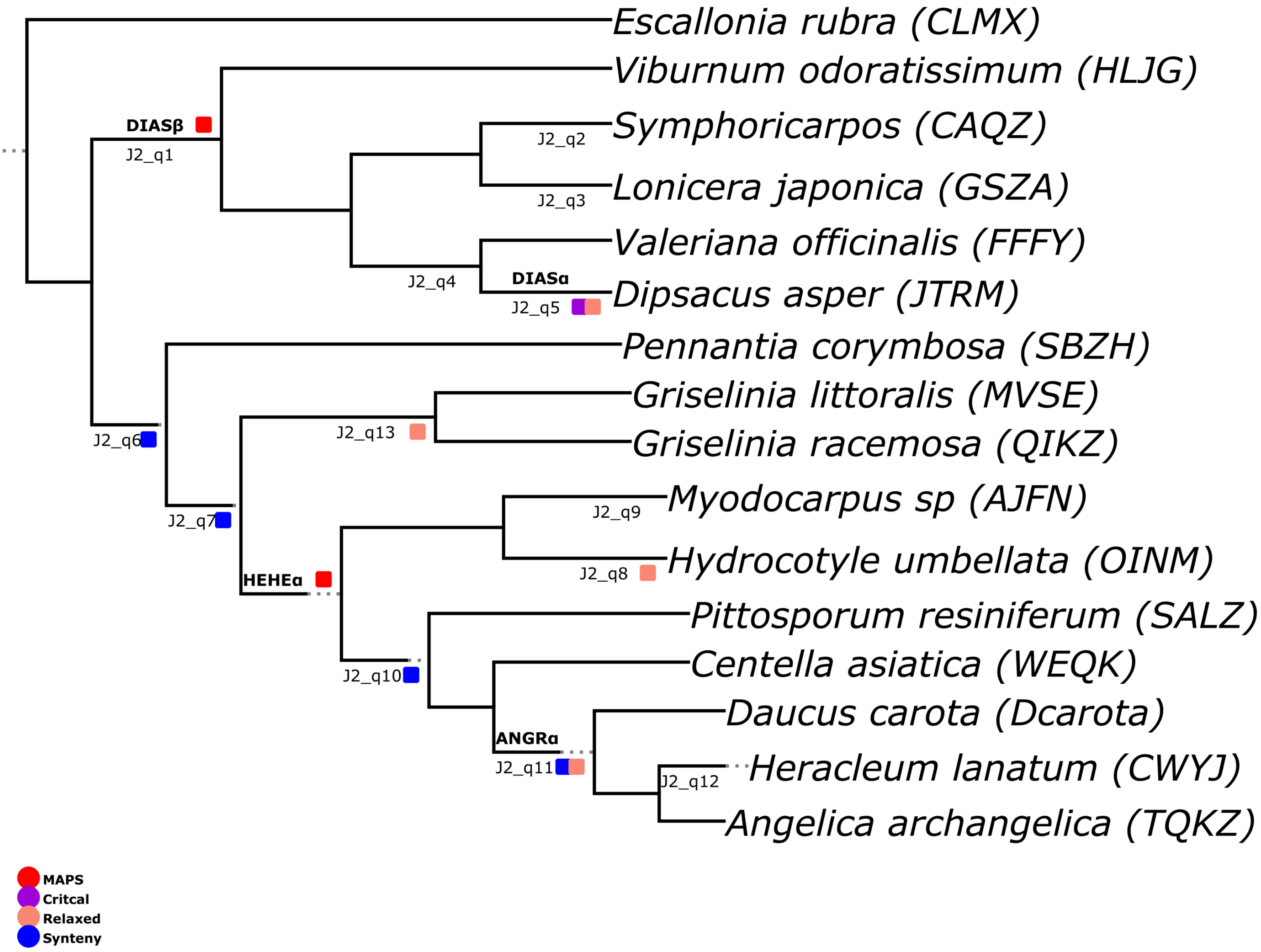


### Figure S68

### K1 Janssens

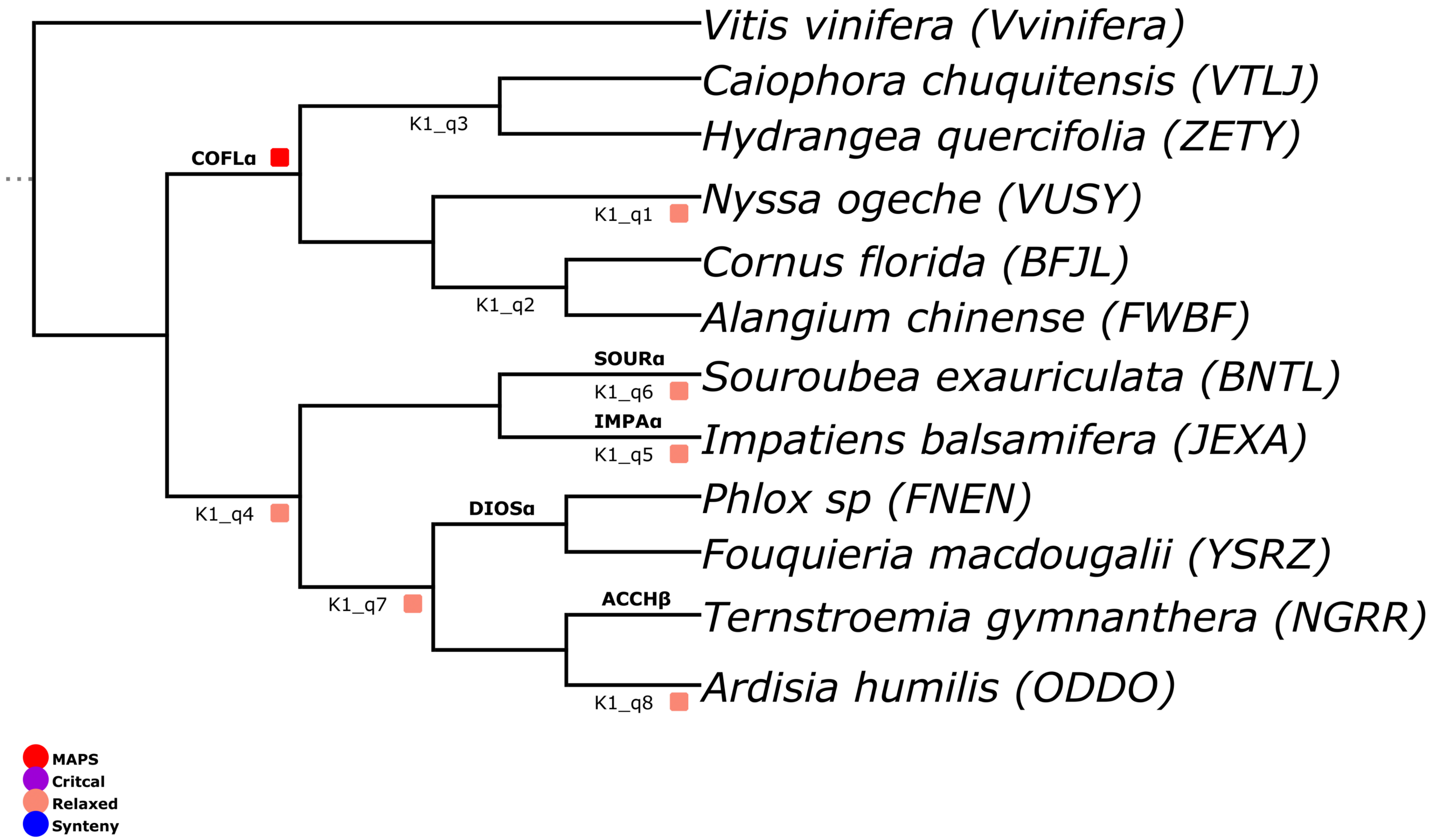


### Figure S69

### K2 Janssens

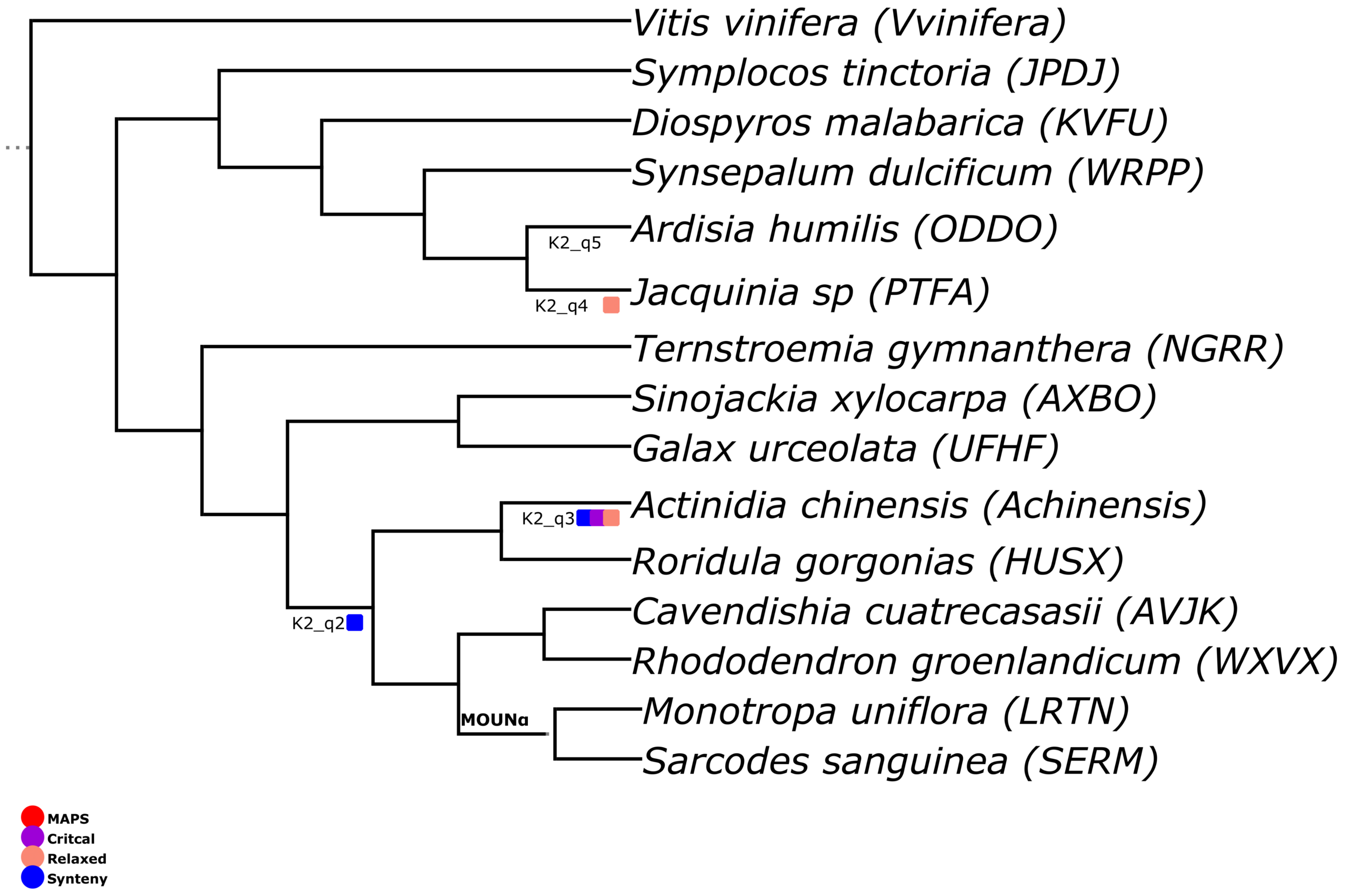


### Figure S70

### L1 Janssens

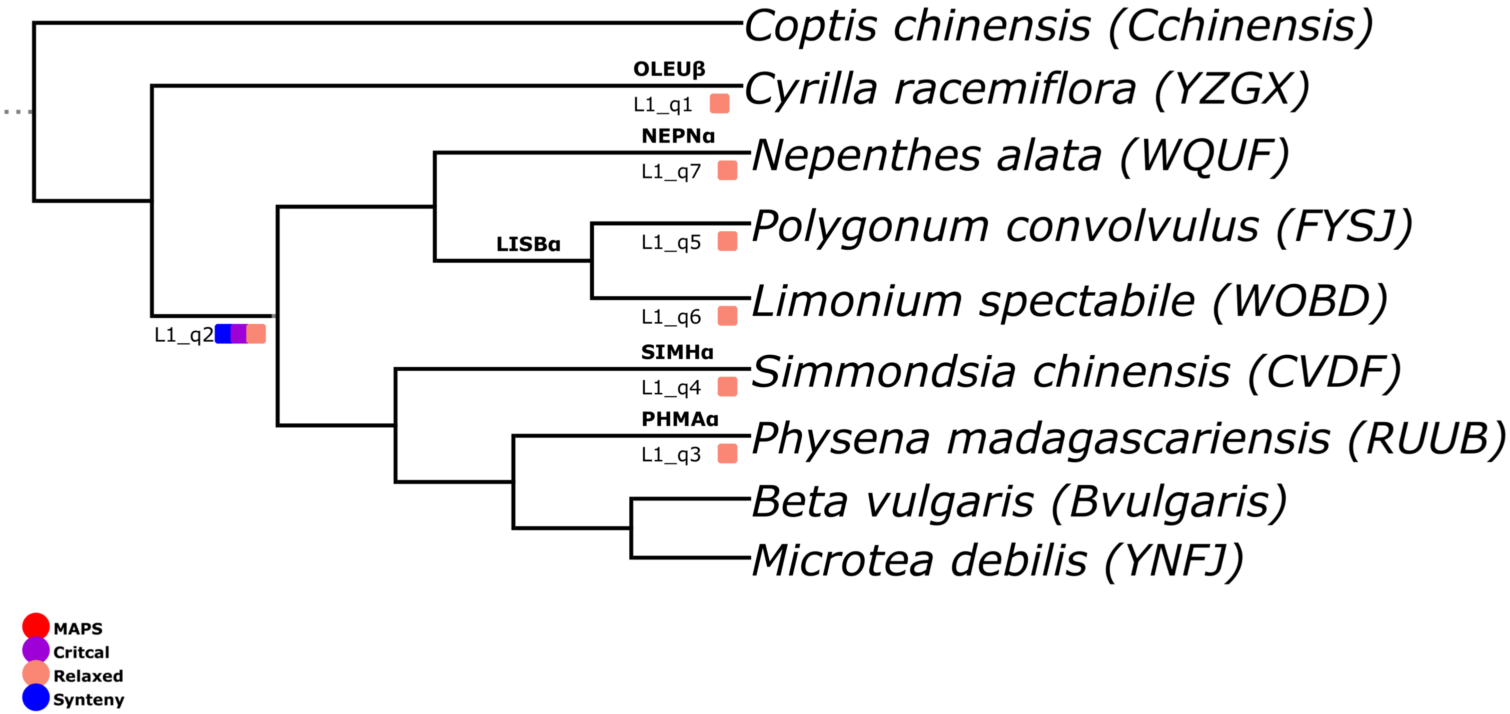


**Figure S71**

### L2 Janssens

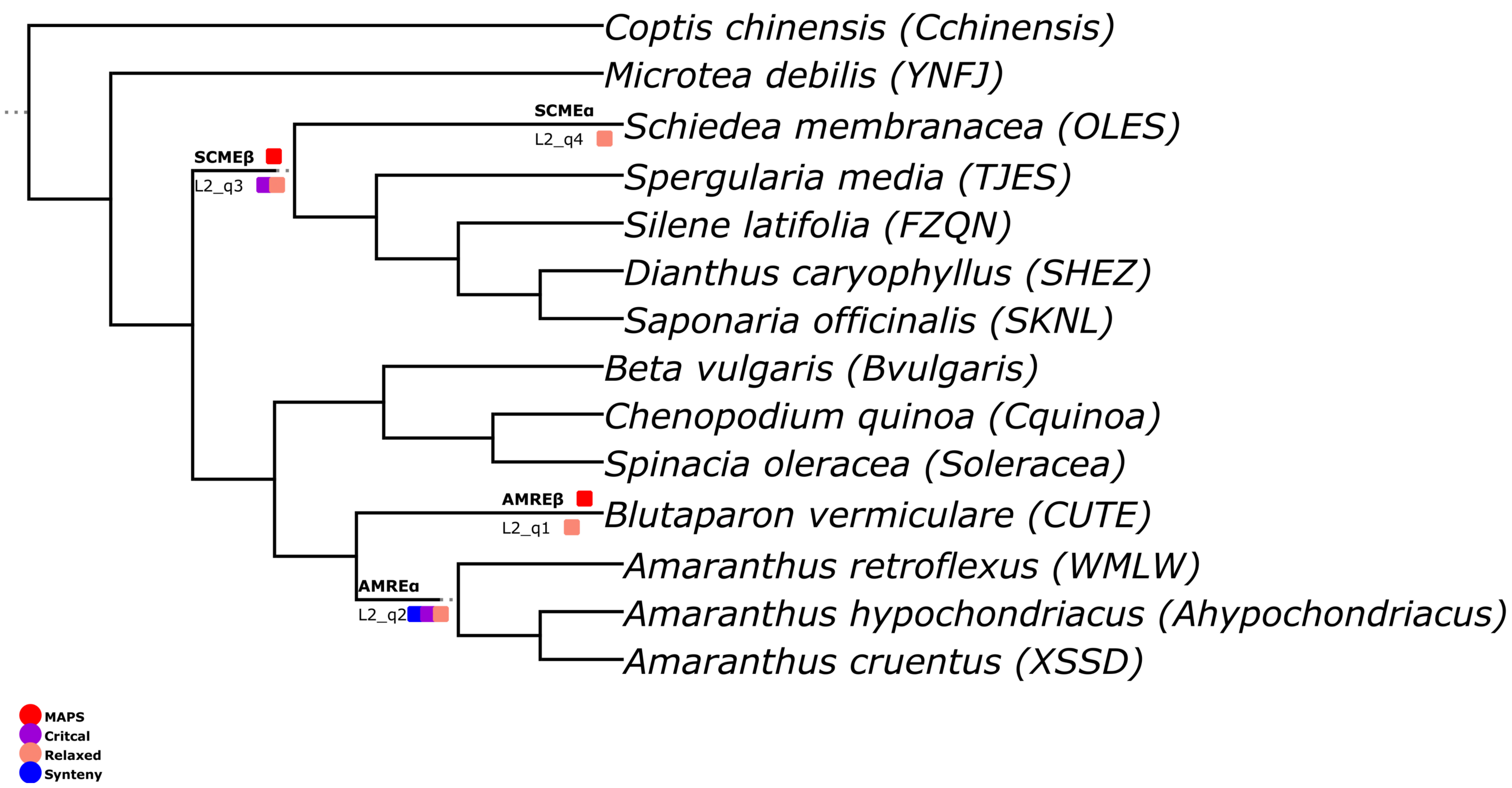


### Figure S72

### L3 Janssens

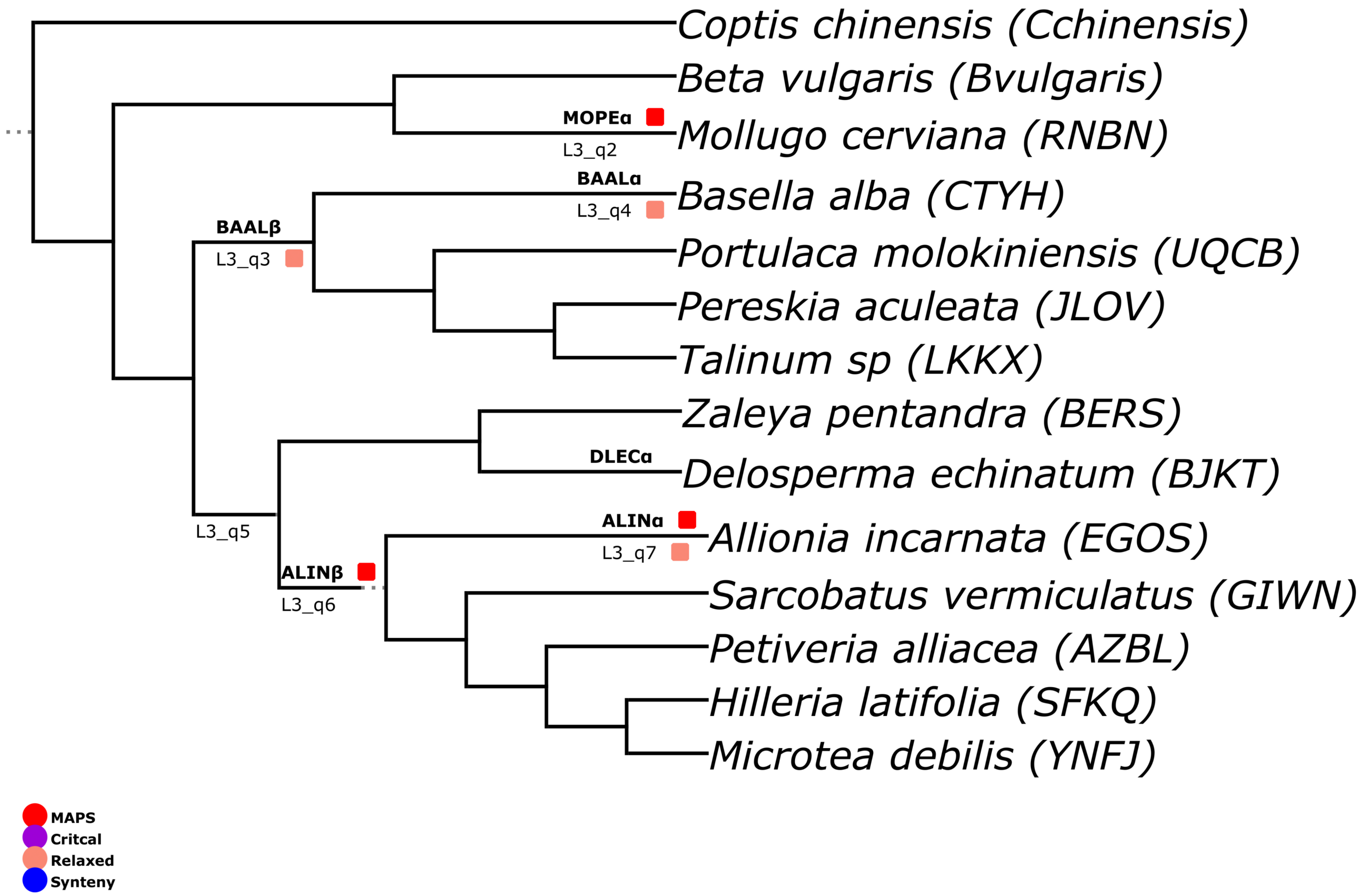


### Figure S73

### M1 Janssens

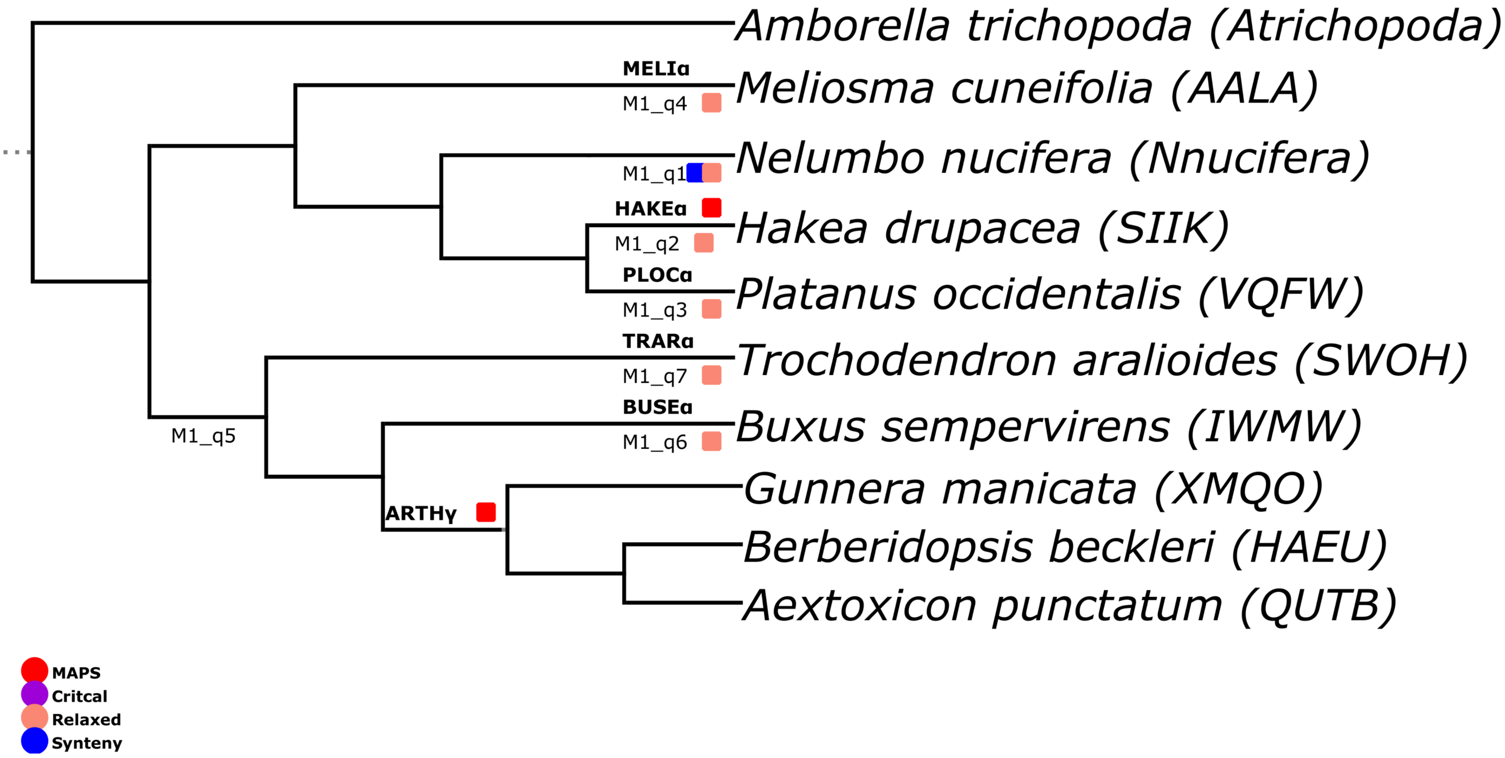


### Figure S74

### M2 Janssens

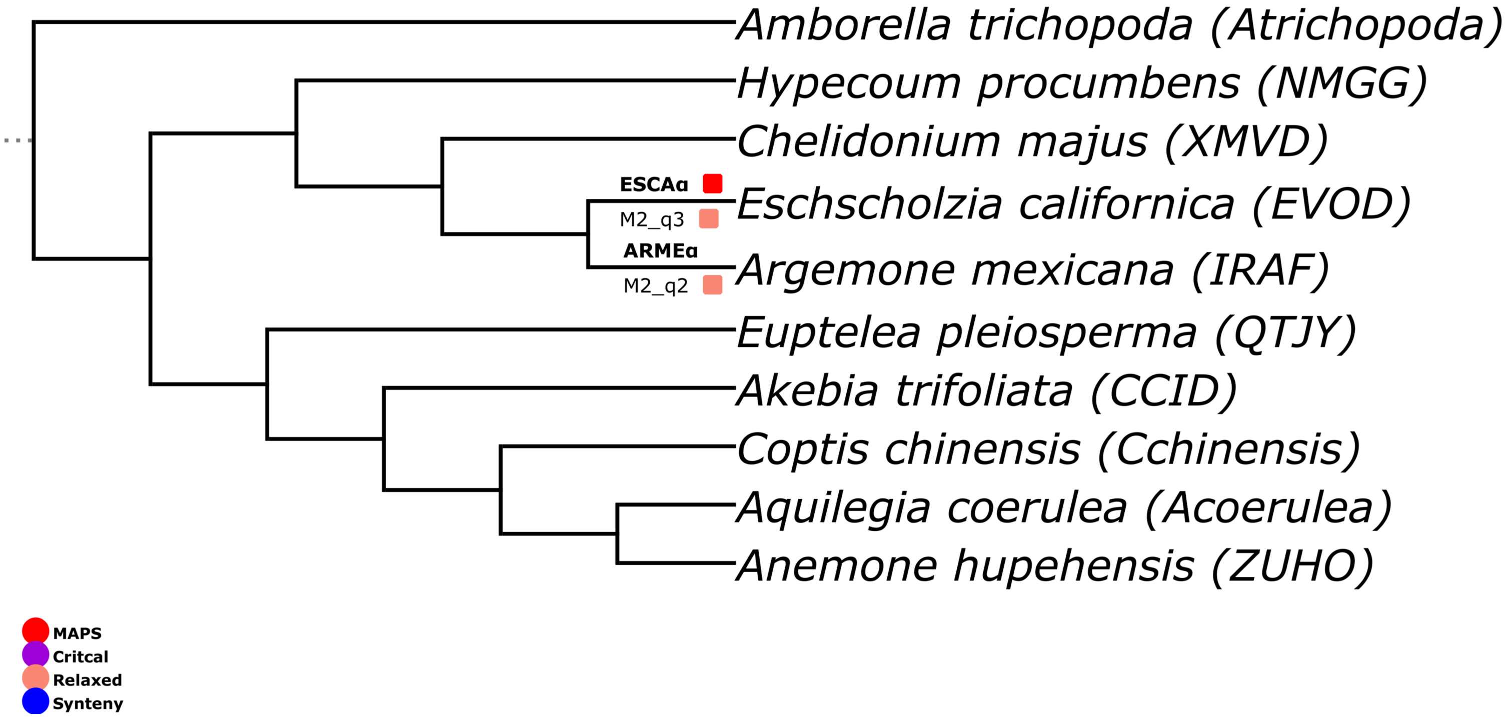


### Figure S75

### N1 Janssens

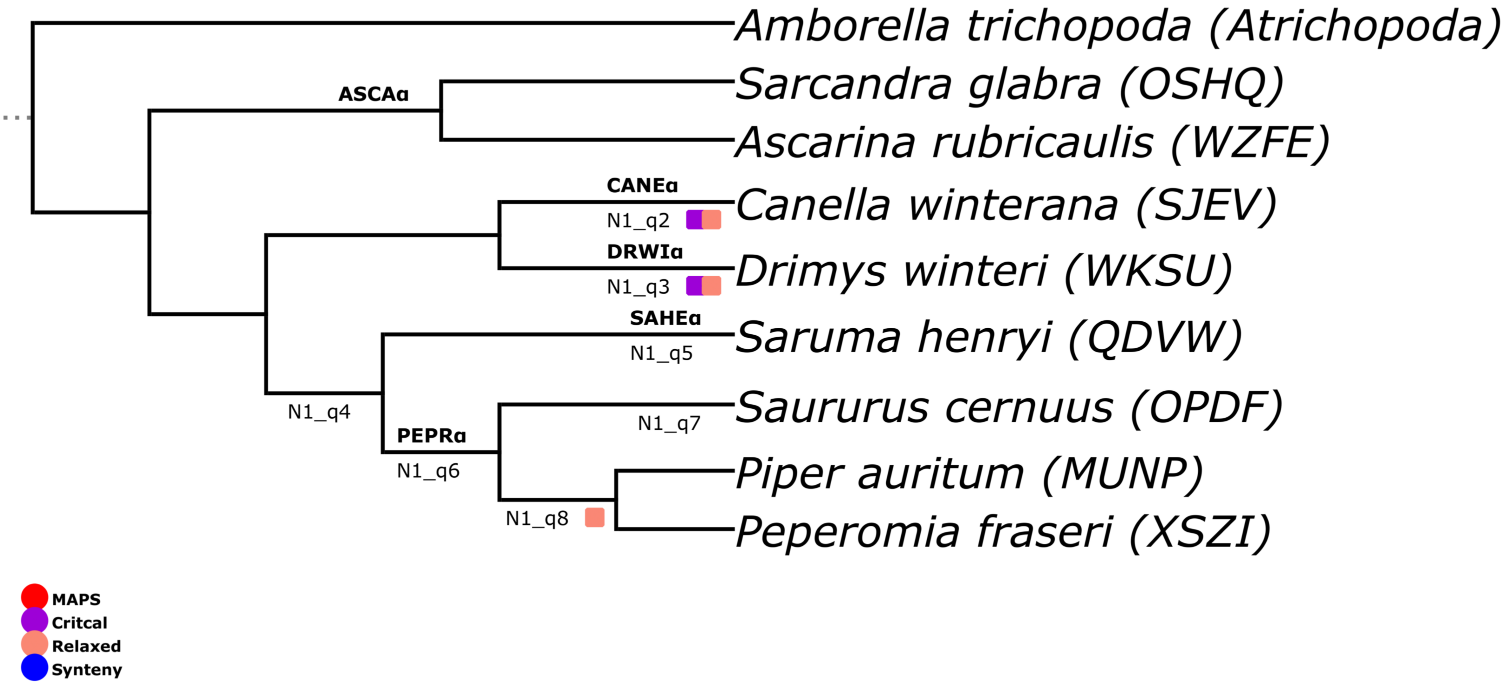


### Figure S76

### N2 Janssens

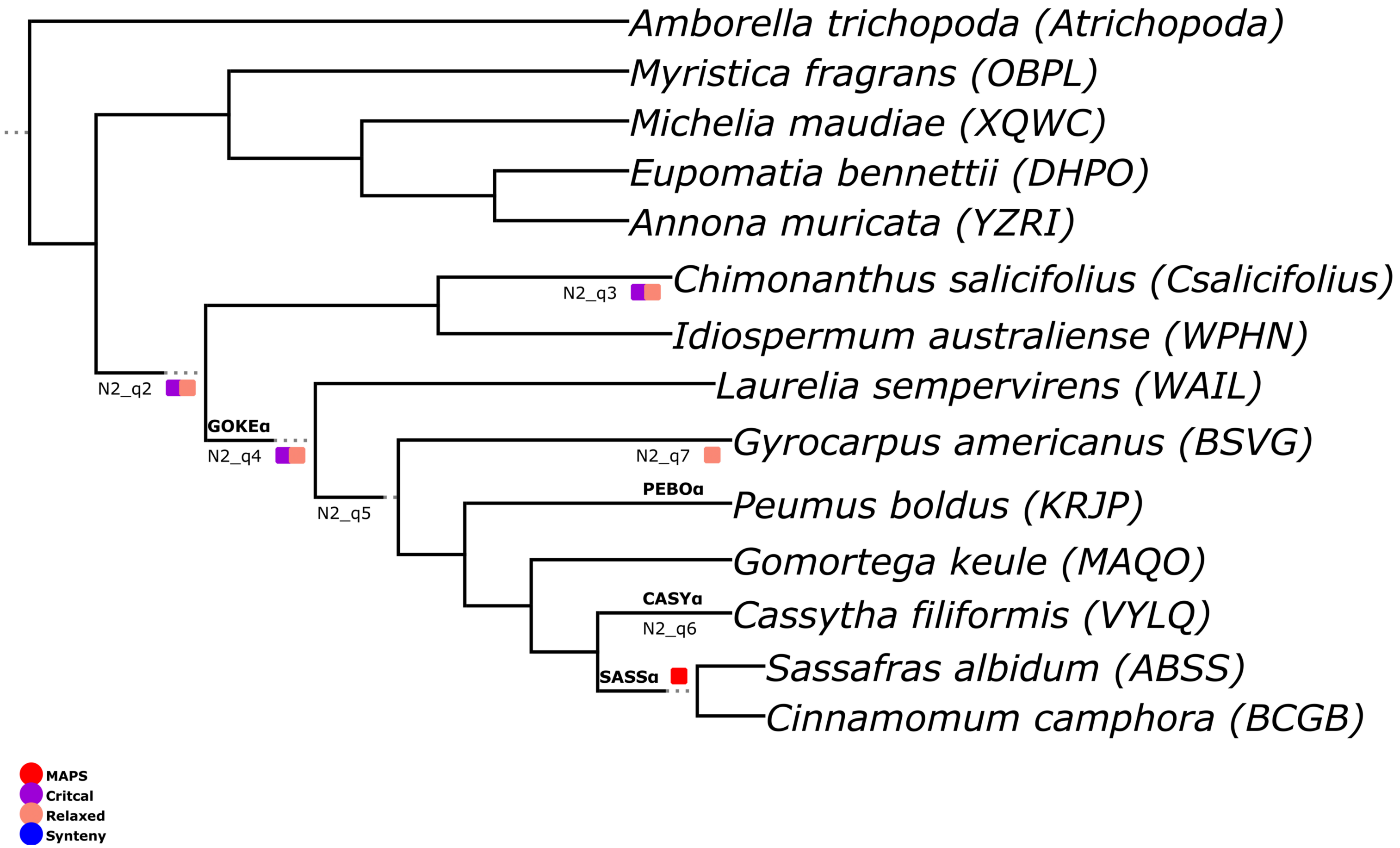


### Figure S77

### O1 Janssens

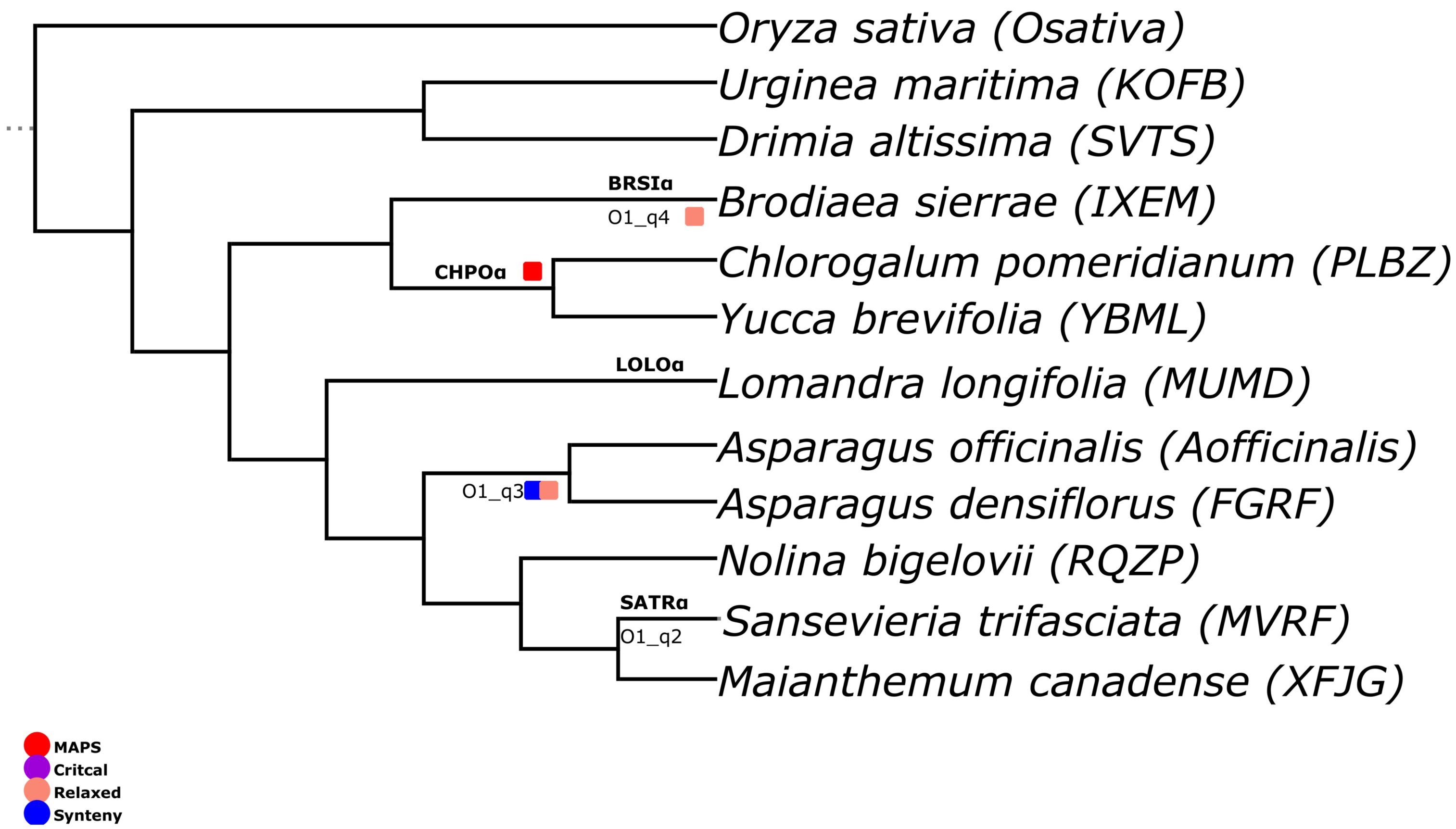


### Figure S78

### O2 Janssens

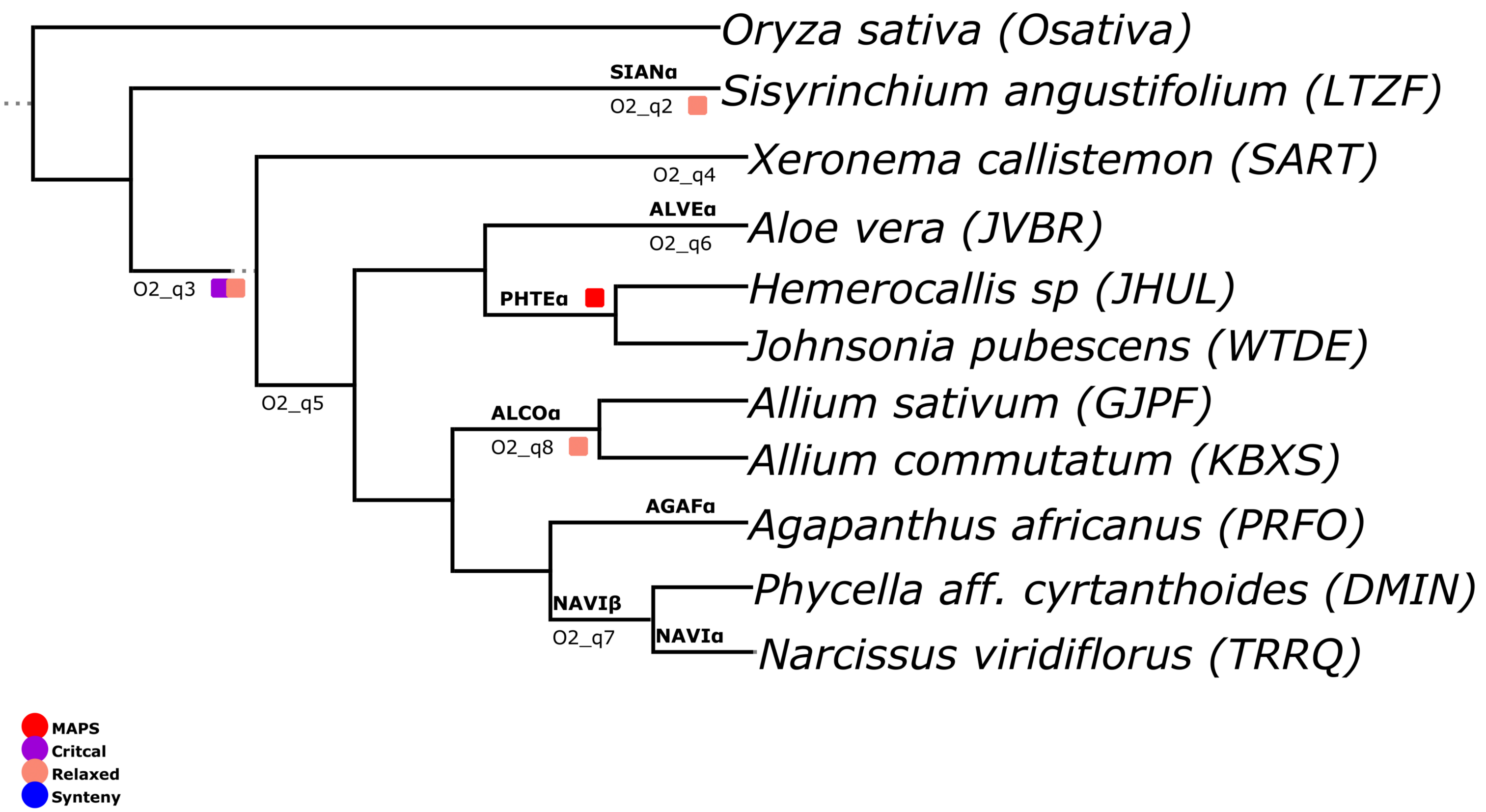


### Figure S79

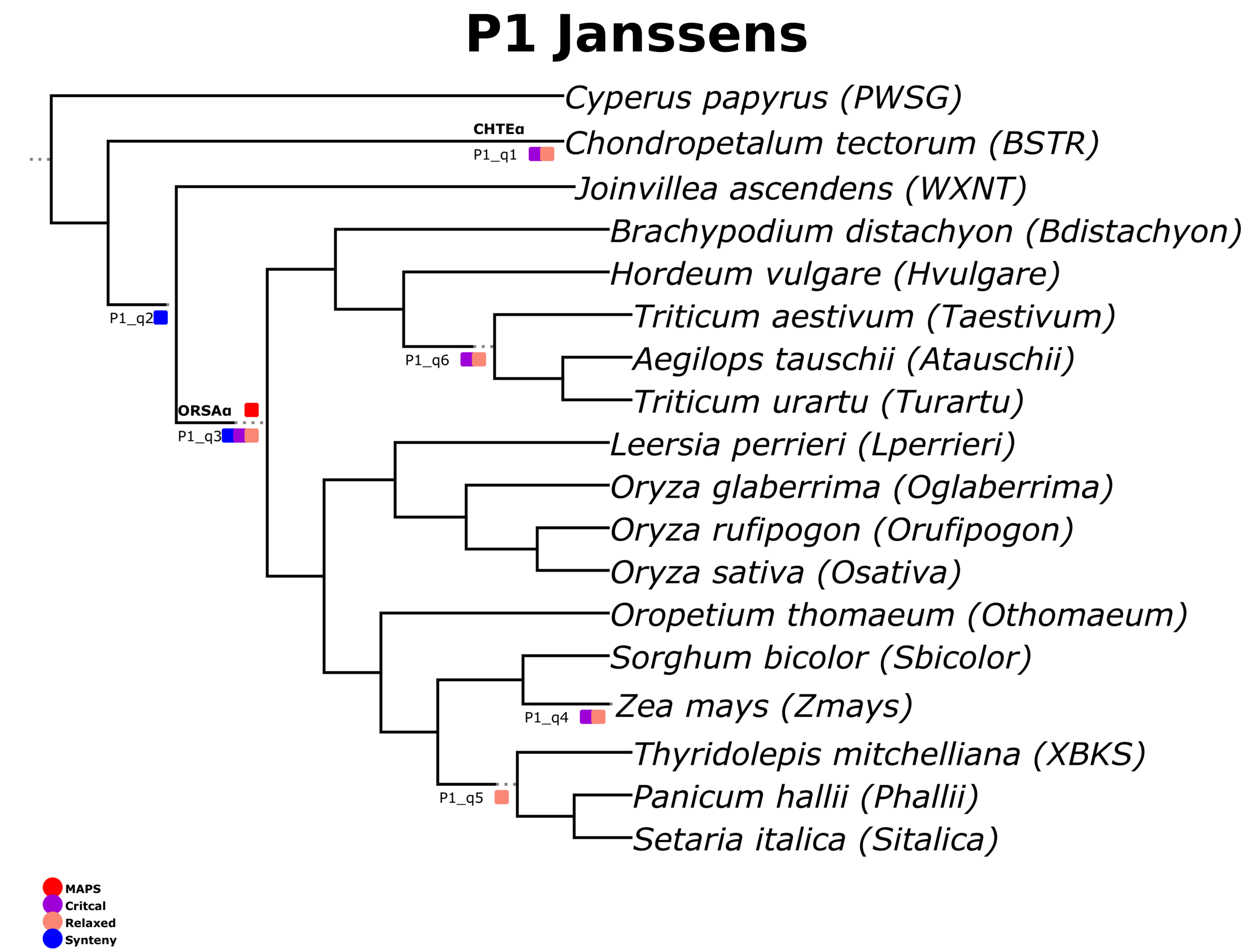
P1 Janssens
Cyperus papyrus (PWSG)
CHTEα
Chondropetalum tectorum (BSTR)
P1_q1
Joinvillea ascendens (WXNT)
Brachypodium distachyon (Bdistachyon)
Hordeum vulgare (Hvulgare)
Triticum aestivum (Taestivum)
P1_q2
P1_q6
Aegilops tauschii (Atauschii)
Triticum urartu (Turartu)
ORSAα
P1_q3
Leersia perrieri (Lperrieri)
Oryza glaberrima (Oglaberrima)
Oryza rufipogon (Orufipogon)
Oryza sativa (Osativa)
Oropetium thomaeum (Othomaeum)
Sorghum bicolor (Sbicolor)
Zea mays (Zmays)
P1_q4
Thyridolepis mitchelliana (XBKS)
P1_q5
Panicum hallii (Phallii)
Setaria italica (Sitalica)
MAPS
Critcal
Relaxed
Synteny

### Figure S80

### P2 Janssens

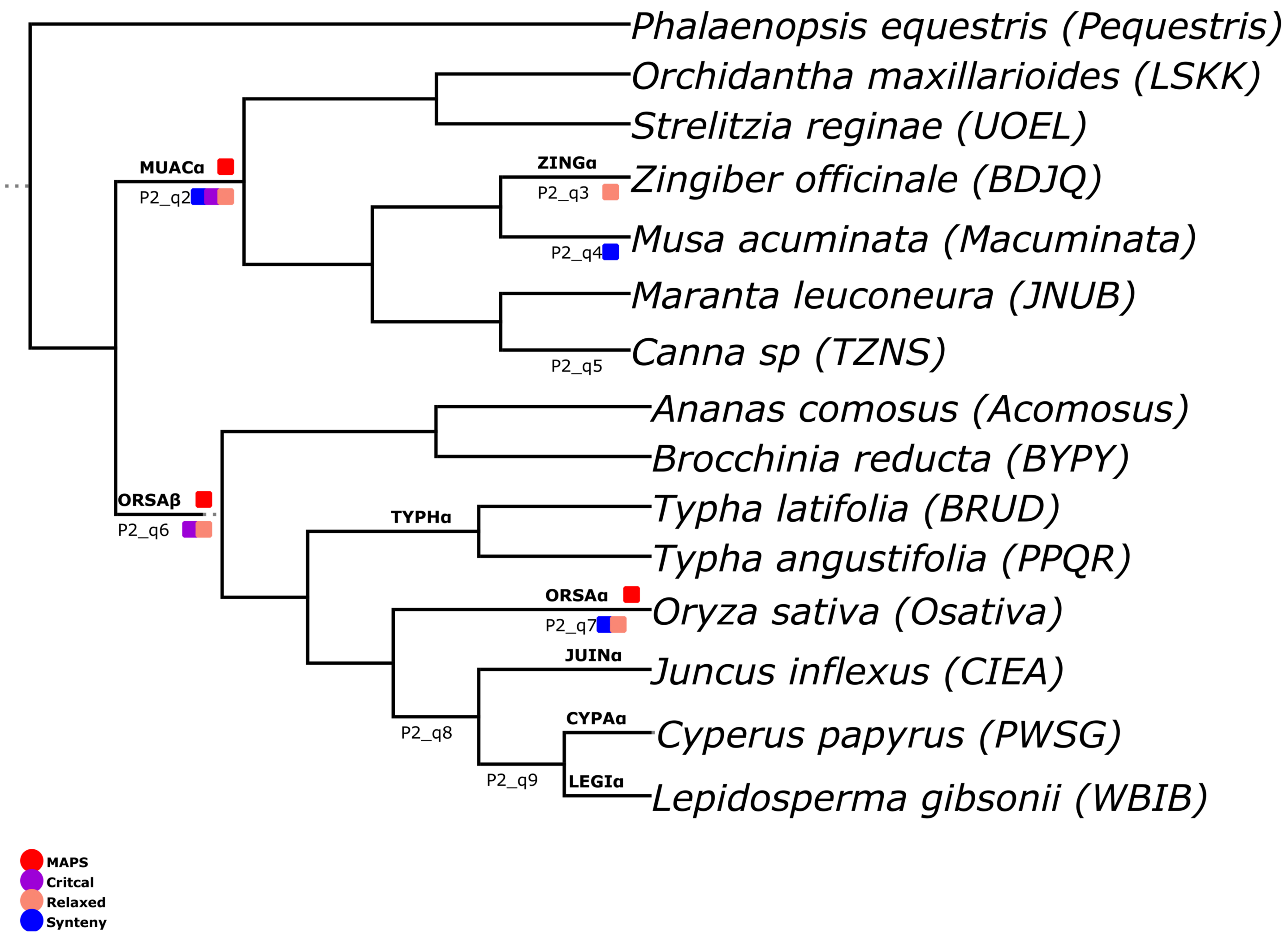


### Figure S81

### P3 Janssens

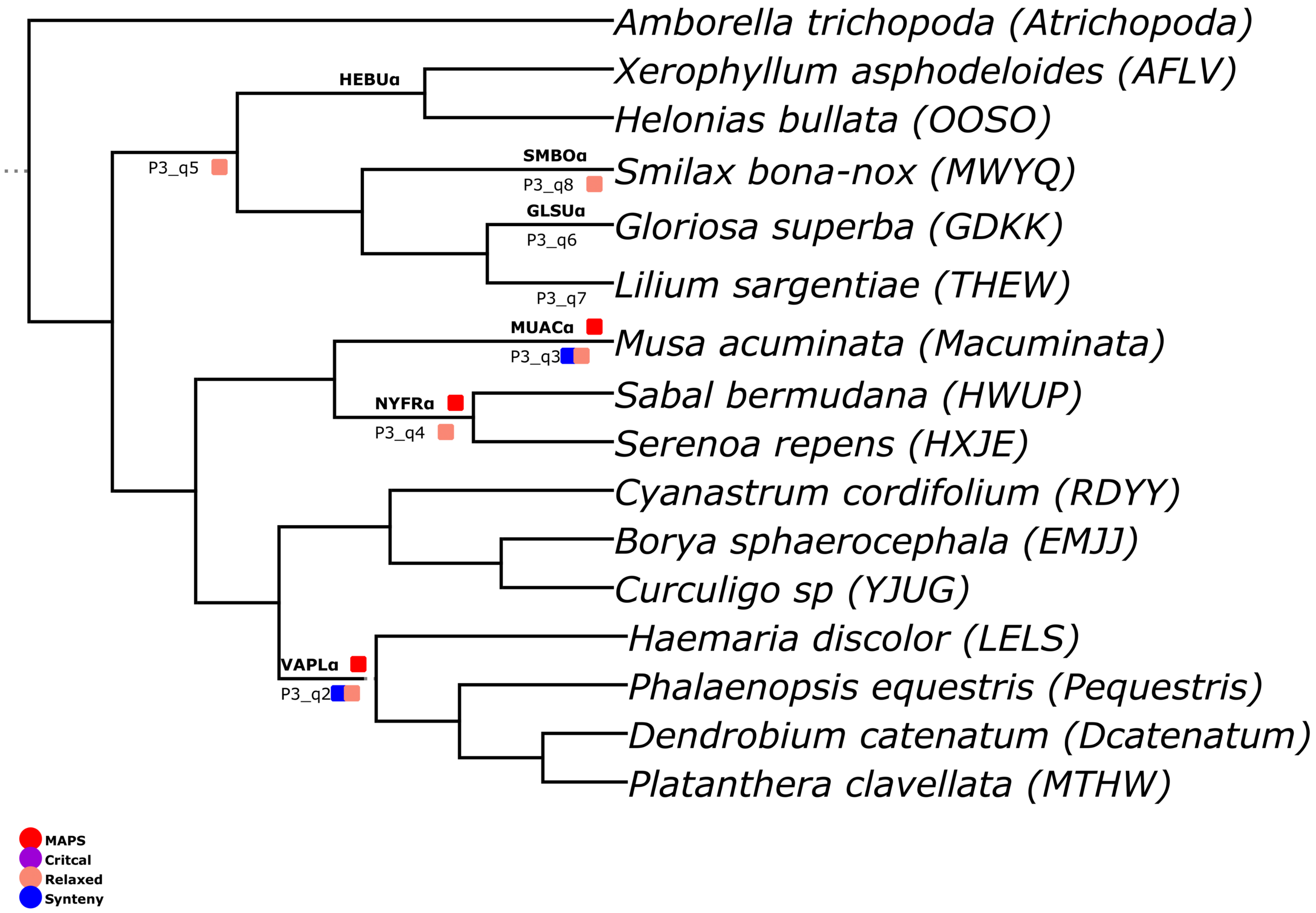


### Figure S82

### Q1A Janssens

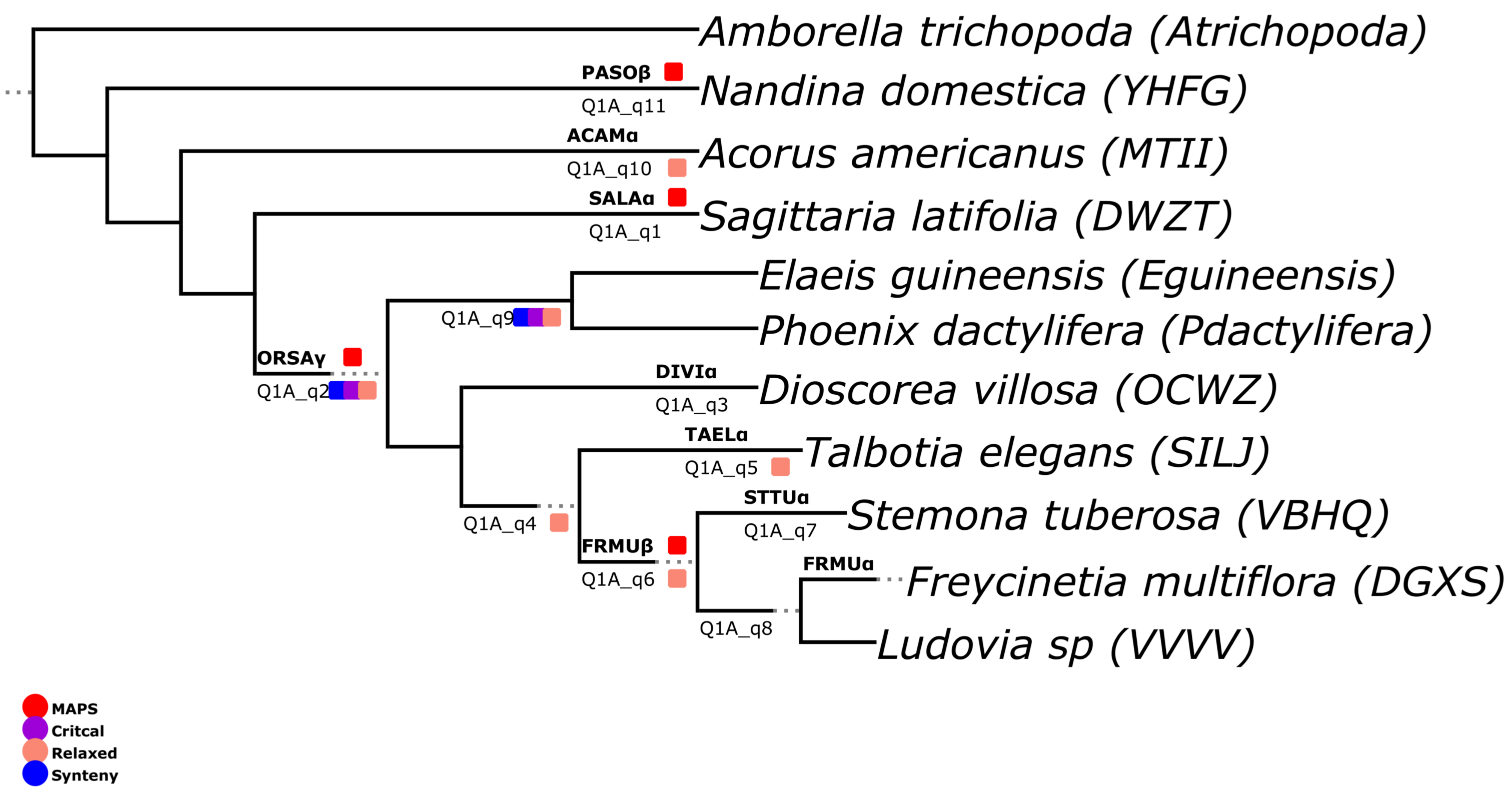


### Figure S83

### Q1B Janssens

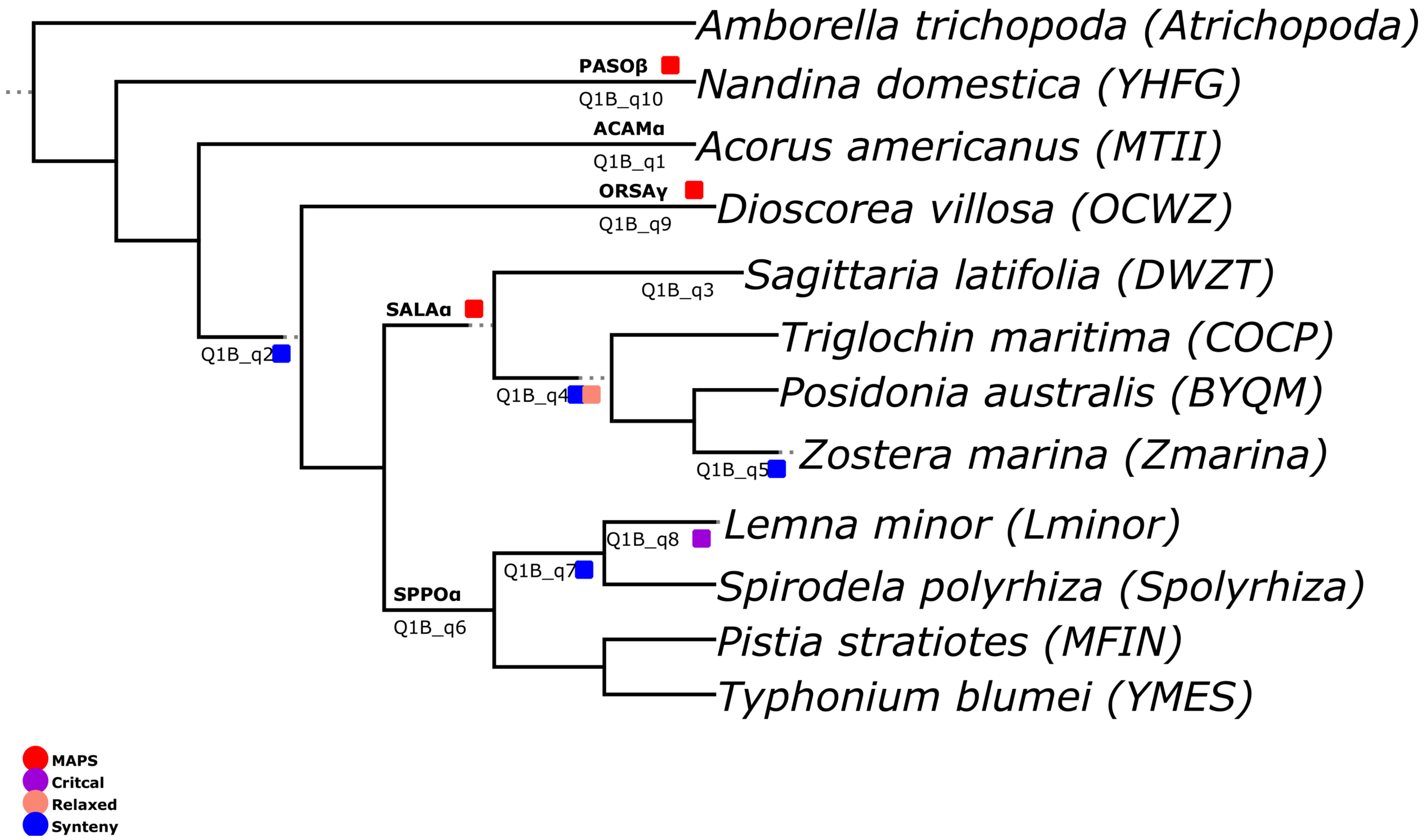


### Figure S84

### Q2 Janssens

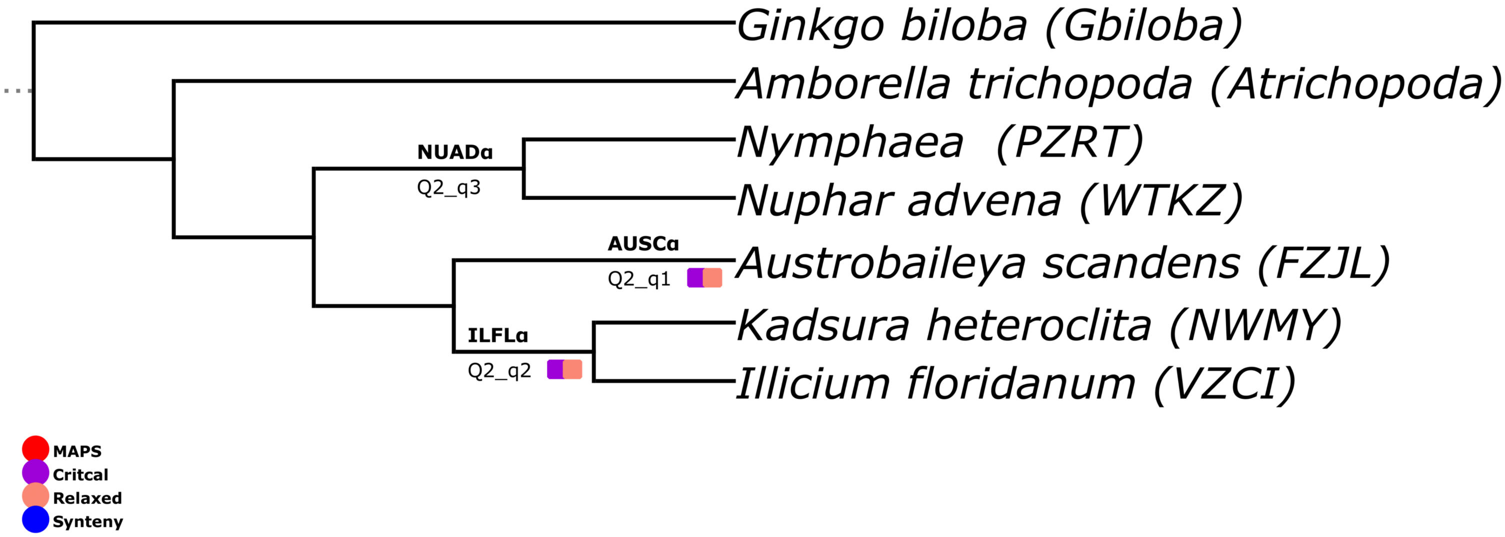


### Figure S85

**Figure S2-85:** Phylogenies for each analysis under the TimeTree and Janssens et al. (2020) phylogenies. Each tree is annotated with hypothesized WGD placements supported by synteny, WHALE models, or MAPS. MAPS inferences were based on the equivalent MRCA for species inferred to share the proposed WGD in the 1KP 2019 analysis. Annotations are as follows: MAPS (red), WHALE Critical (purple), relaxed model (peach), and the syntenic depth test (blue).
